# The double-edge sword of heterogeneous ripening pattern in winter wheat cultivar mixtures: A case study under post-anthesis water stress

**DOI:** 10.1101/705384

**Authors:** Abbas Haghshenas, Yahya Emam, Ali Reza Sepaskhah, Mohsen Edalat

## Abstract

Wheat cultivar mixtures with heterogeneous phenology has a less-explored potential to improve crop diversity, yield stability, and agronomic features particularly in response to the currently increased environmental stresses and uncertainties. To investigate the option of using wheat cultivar mixtures with different ripening patterns for mitigating the adverse effects of post-anthesis water stress, a two-year field experiment was conducted during 2014-15 and 2015-16 growing seasons at the research field of School of Agriculture, Shiraz University, Iran. The factorial experiment was a Randomized Complete Block Design with 3 replicates, in which 15 mixture treatments including monocultures and every 11 possible mixtures of four early- to middle-ripening wheat cultivars were grown under two normal and post-anthesis deficit-irrigation conditions. Measured traits and estimated indices included grain yield and its components, canopy temperature, soil water content, water productivity, susceptibility index, and water use efficiency. The results indicated that under the stressful condition of post-anthesis deficit-irrigation, heterogeneity in the ripening pattern of mixtures was declined. Consequently, dissimilarities in grain yields as well as various agronomic characters of mixture treatments were also lessened. This may be an evidence for the negative effect of water shortage stress on heterogeneity within agroecosystems. Although cultivar mixtures showed some casual advantages in some traits, such beneficial effects were not consistent across all conditions. Moreover, no cultivar mixture produced higher grain yield than the maximum monoculture. Despite the general expectation for beneficial ecological services from cultivar mixtures, in many cases disadvantageous blends were found which led to a considerable reduction in grain yield and water productivity. Therefore, it is suggested that unless the performance, and preferably the involved mechanisms, of cultivar mixtures are not fully understood, use of blends as an alternative for conventional high-input wheat cropping systems may lead to adverse results.

## 1. Introduction

In the modern intensive wheat production systems, intra-specific diversity is at its minimum historical level, as a consequence of decades of homogenization for improving yield and coping with environmental constrains (for instance see Chateil *et al*., 2013 and Bonnin *et al*., 2014). Among the multidimensional reductions in diversity is the intensified homogeneity in patterns of ripening and other phenological events, whose effects are rarely investigated; though the phenology or phenological modifications per se are extensively studied and practiced e.g. in breeding programs. For instance, as a reasonable response to the intensified drought stress and high temperatures during late season, earliness is considered as a vital solution in developing current genotypes into the vulnerable environments, based on the physiological mechanism of stress avoidance (Hu *et al*., 2005; Mondal *et al*., 2013; Mwadzingeni *et al*., 2016; Prieto *et al*., 2018). Therefore, it seems that irrespective the time of occurrence of phenological events (e.g. ripening), the role of decreased heterogeneity in incidence of phenological phases within canopy is less explored, especially under unpredictable environmental conditions.

To restore diversity to the conventional intensive wheat cropping systems, an increasing number of studies are carried out through designing multi-component canopy structures, either utilizing breeding techniques (e.g. multiline cultivars, Mundt, 2002; and composite cross populations suggested by Döring *et al*., 2015) or cultivar mixtures (i.e. blends of genetically uniform cultivars; e.g. see Dubin & Wolfe, 1994; Lopez, & Mundt, 2000; Gallandt *et al*., 2001; Kiær *et al*., 2009; Döring *et al*., 2015; Borg *et al*., 2018; and Reiss & Drinkwater, 2018). In addition to many research works focused on controlling effect of cultivar mixtures on biotic stresses (Wolfe, 1985; Newton *et al*., 1997; Mundt, 2002; Cox *et al*., 2004; Dubs *et al*., 2018; and Tratwal & Bocianowski, 2018), many studies show the results of comparisons between agro-ecological performances of cultivar mixtures and monocultures (e.g. see Baker, 1977; Stützel & Aufhammer, 1990; Smithson & LENNÉ, 1996; Jackson & Wennig, 1997; Thomas & Schaalje, 1997; Finckh *et al*., 2000; Swanston *et al*., 2005; Cowger & Weisz, 2008; Kaut *et al*., 2008; Mengistu *et al*., 2010; Dai *et al*., 2012; Chateil *et al*., 2013; Zając *et al*., 2014; Adu-Gyamfi *et al*., 2015; Barot *et al*., 2017; Newton *et al*., 2017; Lazzaro *et al*., 2018; and Fletcher *et al*., 2019). The theoretical principal beyond the idea of such studies which have made cultivar (or variety/ genotype) mixtures a promising U-turn towards improving sustainability in high-input cropping systems, is the well-known ecological services potentially provided by increasing diversity and dissimilarity (see Smithson & Lenné, 1996; Barot *et al*., 2017; and Reiss & Drinkwater, 2018). Accordingly, it is notable that there is an intrinsic difference between the higher level of heterogeneity in the traditional monoculture systems, and in the ideal sustainable stands purposed by the current studies, i.e. designing the canopy structure more wisely. Indeed, although the mixed and impure seed masses cultivated in the traditional (or subsistence) cropping systems were also resulted from a long-term selection by experienced growers, yet type and degree of heterogeneity within the canopy were not precisely selected or designed. However, supported by the current substantial body of knowledge, intelligent attempts are being made to design specific and purpose-oriented cultivar mixtures. For instance, according to the ecological bases which support the potential improved status of input uptake under enhanced levels of heterogeneity, performance of cultivar mixtures in efficient water consumption is investigated as a potential solution for water deficit conditions prevailed in semi-arid areas (Haghshenas *et al.,* 2013; Fang *et al*., 2014; Adu-Gyamfi *et al*., 2015; Wang *et al*., 2016).

One kind of dissimilarities among the mixture components considered for designing cultivar blends are phenological differences, which are physiologically vital enough to mainly determine the fate of interaction between the crop and environment. Accordingly, Haghshenas *et al*., (2013) evaluated the option of mitigating the intensified post-anthesis competition within the mixed canopy of two early- and middle-ripening wheat cultivars, and reported that the intra-specific competition under various seasons and irrigation conditions was several percent lower in mixtures, compared with monocultures; however, this relative advantage did not lead to significantly higher grain yield and post-anthesis water use efficiency. Besides, the equal mixing ratio (1:1) of the two cultivars was consistently similar to the highest yielding monoculture. Fletcher *et al*. (2019) also investigated the idea of using cultivar mixtures for buffering the risk of early or late flowering of short- to long-duration wheat cultivars under various locations and sowing dates. They reported a 6.4% over-yielding in mixtures across the sowing dates. Moreover, it was concluded that wheat cultivar mixtures can provide growers strategy to manage the uncertain risks of heat and drought stress in frost and heat-prone dryland environments. Despite such instances, yet the concept of phenological heterogeneity seems to have a considerable unexplored capacity to be utilized for penetrating the black box of unstable homogenized monocultures. In the present study, mixtures of early- to middle-ripening wheat cultivars were used to evaluate the effect of inducing heterogeneity in the pattern of late season phenological phases on wheat performance under different water availability conditions (full-irrigation and 50% deficit-irrigation) during post-anthesis.

## 2. Materials and Methods

### 2.1. Field experiments

The field experiment was conducted during 2014-15 and 2015-16 growing seasons (see Haghshenas and Emam, 2019) at the research field of School of Agriculture, Shiraz University, Iran (29°73’ N latitude and 52°59’ E longitude at an altitude of 1,810 masl). Mixture treatment were 15 mixing ratios of four early- to middle-ripening wheat cultivars [Chamran (1), Sirvan (2), Pishtaz (3), and Shiraz (4), respectively] including the 4 monocultures and their every 11 possible mixtures, which were grown with 3 replicates under two full-irrigation and post-anthesis deficit-irrigation conditions (See Fig. S1 part A). The experimental design was RCBD (Randomized Complete Block Design) in which all the 90 (2×2 meter) plots were arranged in a lattice configuration with 1 meter distances. Plant density was 450 plants/m^2^ and seeds were mixed with equal ratios (i.e. 1:1, 1:1:1, and 1:1:1:1 for the 2-, 3-, and 4-component blends, respectively) considering their 1000-grain weights and germination percentages. The planting dates in the first and second growing seasons were November 20 and November 5, respectively; and based on the soil test, only 150 kg nitrogen/ha (as urea) was applied in three equal splits i.e. at planting, early tillering, and anthesis. No pesticide was used and weeding was done by hand. Based on the local practices, irrigation interval was set at ten days, and the amount of irrigation water (i.e. equal to crop evapotranspiration) was estimated as (Shahrokhnia and Sepaskhah, 2013):

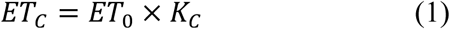

where *ET_C_* is the crop potential evapotranspiration, *ET*_0_ is the reference potential evapotranspiration, and *K_C_* is the crop coefficient (which was taken from Shahrokhnia and Sepaskhah, 2013). *ET*_0_ was estimated using Fao-56 Penman-Monteith model with local corrected coefficients (Razzaghi and Sepaskhah, 2012; Shahrokhnia and Sepaskhah, 2013). In the deficit-irrigated treatment, water was supplied similar to well-irrigation condition until anthesis and afterwards, water was supplied up to 50% *ET_C_*. Irrigation was carried out using hose and flowmeter.

### 2.2. Quantitative monitoring of ripening

Since determining and reporting the ripening trend in cultivar mixtures with heterogeneous phenology based on the conventional qualitative approaches are difficult, laborious, and challenging task, an independent novel image-based method was developed for recording the comparative progress of ripening (see Haghshenas and Emam, 2019). In this method, the ripening trend was determined as the binomial decreasing trend of the well-known canopy coverage (CC) index calculated for each plot, and the ripening date was reported as the date (or accumulated thermal time, ATT) at which CC became zero in the late season. Canopy coverage was calculated as:

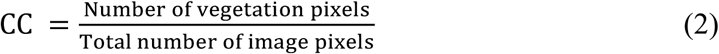

(see Guevara-Escobar *et al*., 2005; and Wang *et al*., 2016)

The diurnal temperatures for calculating accumulated thermal time (ATT) and growing degree days (GDD) were obtained from the weather station located near the experimental site (Haghshenas and Emam, 2019). The accumulated thermal time was calculated by summing the average diurnal temperatures (°C) from sowing to ripening i.e. CC=0). Besides, the growth and developmental phases including early spike development of the 4 cultivars in the monocultures were also recorded based on the conventional phenology (see Haghshenas and Emam, 2019; and Fig. S2).

### 2.3. Measurements and calculations

Based on the study purposes, the main focus of the field measurements was put on the post-anthesis period. From the first deficit-irrigation practice (around anthesis), canopy temperatures were measured using an infrared thermometer (Terminator^®^ TIR 8861) at solar noon, both exactly before and also 48 hours after irrigation. Average of ten readings were recorded for each experimental plot and the overall duration of measurement was about 30 minutes in each date. Although it was attempted to record the data under calm air condition and clear sky, at the time of analyzing it was found that regardless of the date and season, there was a strong declining trend in the recorded temperatures from the first plot to the last one (see Fig. S3). Since plots were randomly arranged in the experimental design and also were visited every time in a regular path, this decreasing trend was attributed to the natural decline in the afternoon air temperature. Therefore, the temperature values of each date were normalized as following:

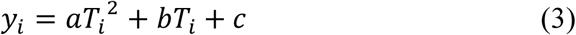

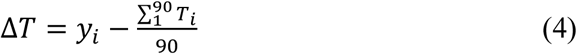

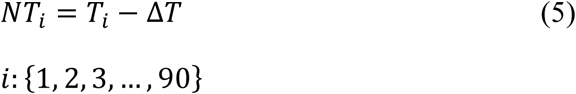

where *i* is the plot number, *y*_*i*_ is the predicted temperature based on the binomial equation of the decreasing trend (the point on the equation trend line), *T*_*i*_ is the recorded temperature of plot *i*, Δ*T* is the difference between *y*_*i*_ and averaged temperature of all plots, and finally, *NT*_*i*_ is the normalized canopy temperature. It is notable that in the present study, the comparative values were important rather than the absolute canopy temperatures. In order to improve analyses, the recorded canopy temperatures across the post-anthesis period were arranged in three groups as follows: (i) Pre-irrigation temperature; which shows the average values of the measured temperatures at the irrigation dates i.e. exactly before irrigation; (ii) Post-irrigation temperature; which is the averaged values of temperatures recorded 48 hours after irrigation; and (iii) Mean temperature: average of pre- and post-irrigation temperatures, or the average canopy temperatures across the post-anthesis period.

From the first deficit-irrigation to the end of season, soil water contents (SWC) of the treatments at the end of irrigation intervals were measured using the gravimetric method (soil samples were oven dried at 110 °C for 24 hours). Having high number of plots (90 plots) and too limited time, sampling was restricted to two depths of 0-30 and 30-60 cm, from two out of the three replicates of the experiment, i.e. 60 plots). Multiplying the gravimetric values by the soil bulk densities of the corresponding depths, soil water content is reported as volumetric percentage.

Late in the season, plants were harvested from the center of plots (equal to overall row length of 3 meter, or 0.6 m^2^ per plot) and grain yield and its components were determined using a laboratory thresher and weighing balance. In order to count grain number m^−2^, an image processing system (Visual Grain Analyzer, TM) was used, by which almost every grains were counted with high precision. Moreover, since the number of spikes m^−2^ were determined much sooner than applying the deficit-irrigation treatments, it was calculated with 6 replicates.

Water productivity (kg m^−3^) was estimated by dividing the grain yield (kg ha^−1^) by the total volume of input water (m^3^ ha^−1^) as follows (almost similar to the method used by Mosaffa and Sepaskhah, 2019, for calculating irrigation water productivity):

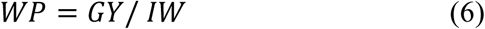

where WP is the water productivity, GY is the grain yield, and IW is the input water, which was in turn calculated as the summation of the volume of irrigation water applied across the season and effective rainfall. Volume of effective rainfall was estimated using the S.C.S. (U.S. Soil Conservation Service) method (SCS, 1972). Due to difference in ripening time for different cultivars in the mixtures, for calculation of the applied water the following point should be considered. The point is that the monocultures of the 4 early- to middle-ripening cultivars had different ripening patterns, and according to the present experiences, the 1^st^ and 2^nd^ cultivars often require one irrigation less than the others. However, verifying this requirement in a replicated experiment where the ripening of individual plots of a given cultivar may be potentially different due to their relative locations in the field, may be challenging. The problem is expected to become even more complicated in cultivar mixtures. For this reason, the below procedure was followed:

a. At the last irrigation (i.e. the 3^rd^ irrigation after anthesis), all plots were irrigated with the estimated volume for applying the well- or deficit-irrigation, regardless of the degree of their ripening.
b. As a routine practice in the present study, all plots were imaged exactly before, and also 72 hours after irrigation. Assessing the pair images (taken before and after irrigation), the overall greenness of each plot was visually evaluated, and it was verified if the plot had required to be irrigated or not. In the latter case, the volume of the last irrigated water was not included in the calculations of input water (for more information about the effect of late season irrigation on the comparative canopy coverage and greenness of the 4 same cultivars, see Haghshenas and Emam, 2019).

Water use efficiency was calculated using the following equation:

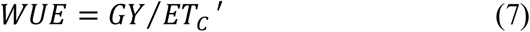

where *WUE* is the water use efficiency (kg ha^−1^ mm^−1^), *GY* is the grain yield (kg ha^−1^), and *ET_C_*′ is the actual evapotranspiration (mm) estimated using the water balance equation in the same way reported by Gao *et al*. (2009):

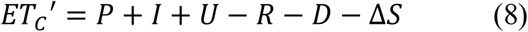

where P is the effective precipitation (mm), I is the irrigation quota (mm), U is the upward capillary flow (mm), R is the runoff (mm), D is the deep percolation from root zone (mm), determined by using hydraulic conductivity, as reported by Sepaskhah and Ilampour (1995), and ΔS is the change of water stored in the 0-60 cm soil layer (mm). Amounts of the upward flow, the runoff, and effective precipitation were negligible for the two seasons (there was no precipitation after flowering).

As a common analysis in cultivar mixture studies, various attributes of the mixtures were compared with the averaged values of the corresponding monocultures. In the present study, such comparisons were named Analysis of Deviation from Monocultures (or ADM), and the results were reported as the deviation percentage i.e. the difference percentage between the actual measured values and the predictions estimated as the mean of values of the corresponding monocultures.

Susceptibility index (SI, Blum *et al*., 1989) was used for comparing the susceptibility of grain yield and yield components of monocultures and mixtures to the irrigation treatment. This index was calculated as:

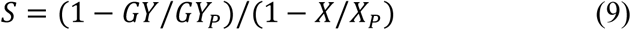

where *GY* is the grain yield under deficit-irrigation, *GY_P_* is the grain yield under well-irrigation, and *X* and *X_P_* are the average yield over all 15 mixture treatments under deficit- and well-irrigation conditions, respectively.

In order to have an overall understanding of the mixtures performance across the 4 environmental conditions (two irrigation treatments × two seasons), giving equal weights to higher yield and less variation, a simple index named *consistent desirable index* (CDI) was developed and used as follows:

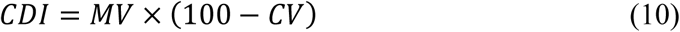

where *MV* is the mean value (of grain yield or other variables) across the 4 conditions, and *CV* is the corresponding coefficient of variation (%). Afterwards, the ranking of mixture treatments in the lists of *MV*, 100 − *CV*, and *CDI* were determined. Similar analyses were also carried out for other measured traits.

Statistical analyses including ANOVA and mean comparison analysis were carried out using IBM SPSS Statistics for Windows, Version 19.0 (Armonk, NY: IBM Corp.). Mean comparisons related to RCBD were performed using LSD and Tukey’s tests. Besides, various orthogonal tests were carried out for comparing several dual conditions created by mixing of the four cultivars. Accordingly, based on Levene’s test of homogeneity of variances, appropriate degrees of freedom and P-values of the dual-grouped treatments were used for comparisons. Data mining was carried out using RapidMiner studio version 9.2. All charts and figures were made and edited by Microsoft Excel 2016 and Adobe Illustrator CC 2019.

## 3. Results

As shown in Fig. 1, during the post-anthesis period in both seasons, diurnal mean air temperature and relative humidity had an increasing and decreasing trend, respectively, and this pattern contributed to higher reference potential evapotranspiration (ET_0_). Total seasonal rainfall was 205 and 267 mm in the first and second years, respectively, which almost occurred before anthesis.

**Figure 1.**
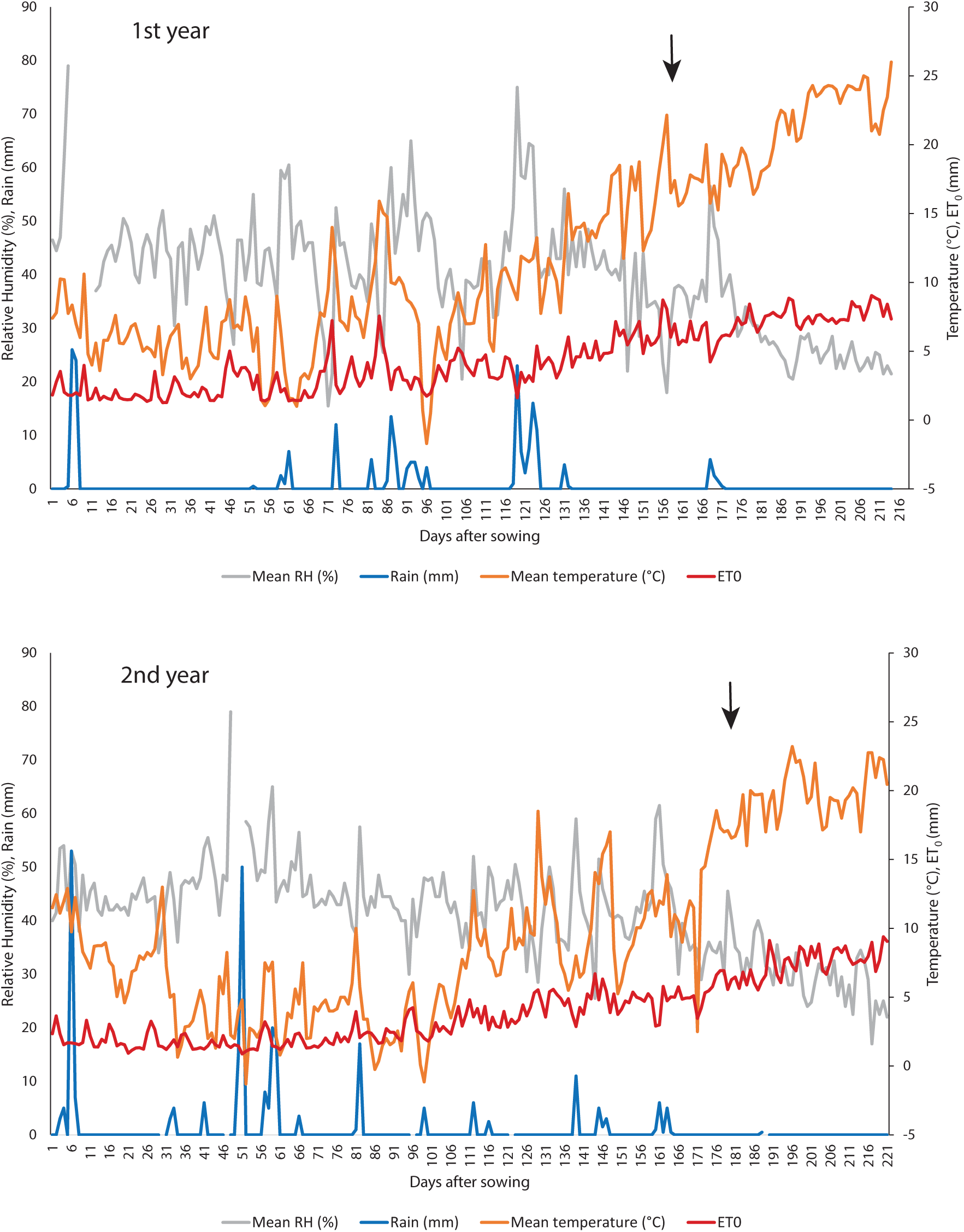
The meteorological data of the two growing seasons (i.e. 2014-15 and 2015-16). The environmental condisions during post-anthesis period was potentially more stressfull, as there were no rainfal, relative air humidity decreased, mean diurnal temperature, and consequently ET0 had increasing trends. The black arrows show the flowering time of the early ripening cultivar i.e. the 1st cultivar.

### 3.1. Ripening pattern

The comparative image-based patterns of ripening in mixture treatments (i.e. the 15 treatments including monocultures) are presented in Fig. 2. Accordingly, the following results are obtained:

- In general, the comparative thermal time requirements for ripening of the 4 cultivars in monocultures followed the expected pattern of the 1^st^ < 2^nd^ < 3^rd^ < 4^th^ cultivars.
- None of the mixtures had a ripening date out of the range of the monocultures; therefore, irrespective the season and irrigation treatment, the ripening time of the 1^st^ and 4^th^ cultivars were two extremes.
- Under deficit-irrigation conditions, the differences between the ripening patterns of the mixture treatments were decreased, and the ripening dates were more similar to each other.

**Figure 2.**
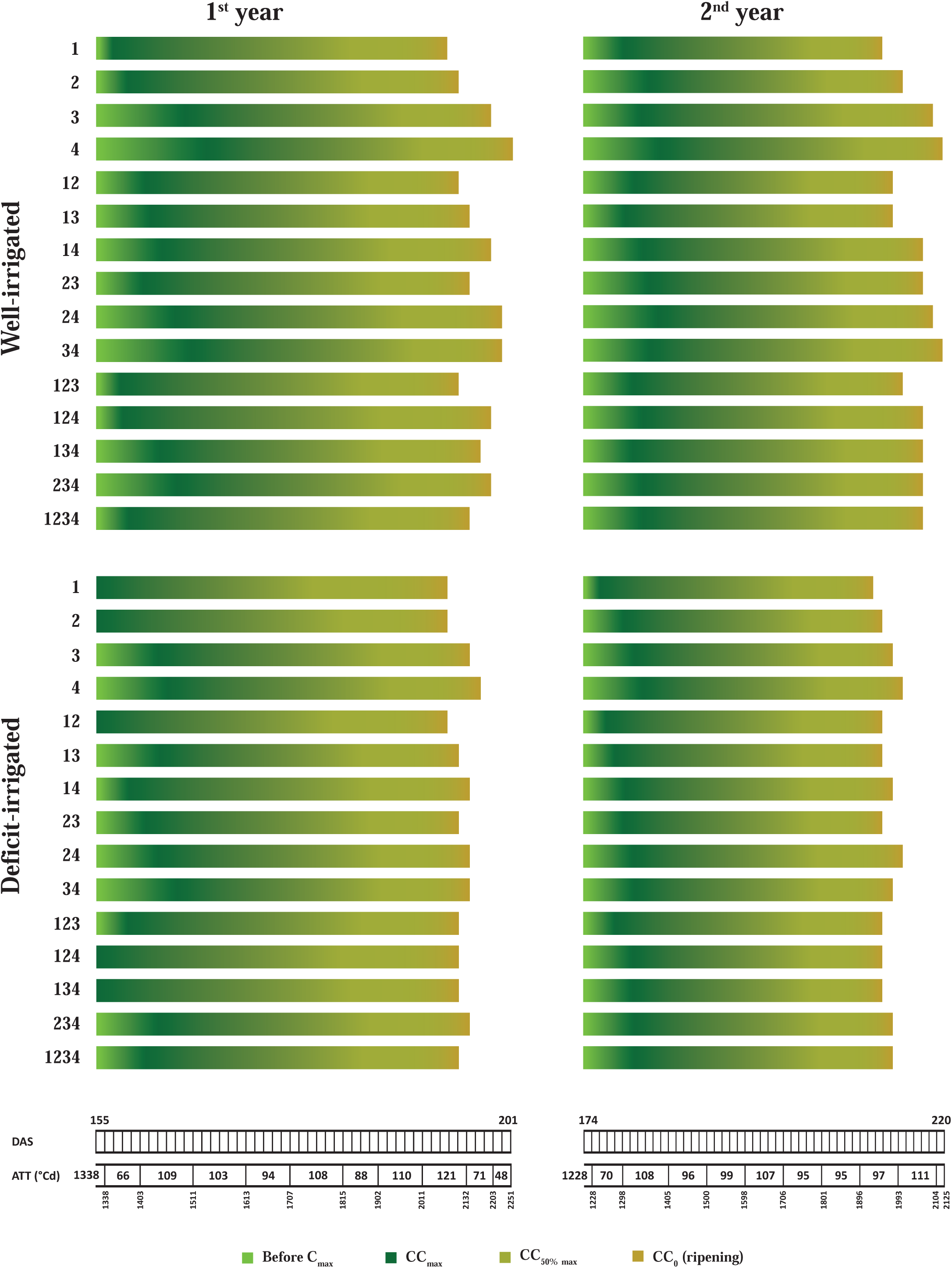
Comparative phenologies of wheat culitvar mixtures across two years and well- and deficit-irrigation conditions. DAS: days after sowing; ATT: accumulated thermal time (°Cd)

The main effects of year, mixture treatments, and also irrigation treatment on ripening were significant (Table 1). In general, ripening (which was determined as accumulated thermal time, ATT, required for reaching to CC_0_ phase) was occurred earlier under deficit-irrigation. Also in the second year, well-irrigated plots were ripened earlier, compared with that obtained in the first year (Table 2). With an exception for the case of well-irrigated treatment in the first year, it was found that irrespective of the environmental conditions (i.e. irrigation and year), there was no significant difference in the ATTs required for ripening of neither the 1^st^ and 2^nd^ cultivars, nor the 3^rd^ and 4^th^ cultivars. Therefore, it seems that the monocultures may be categorized in two major ripening groups, whose comparative effects on various crop properties were evaluated later using orthogonal comparisons.

**Table 1.** Analysis of variance for ripening (accumulated thermal time, ATT).

| Source of variation | Ripening (ATT) |  |  |  |  |
| --- | --- | --- | --- | --- | --- |
|  | df | SS | MS | F | P-value |
| Year | 1 | 650409.6 | 650409.6 | 1185.884 | 0.000 |
| Mixture treatments (Mix.) | 14 | 140821 | 10058.65 | 18.34 | 0.000 |
| Irrigation (Irr.) | 1 | 123160.2 | 123160.2 | 224.556 | 0.000 |
| Replicate (Rep.) | 2 | 266.997 | 133.498 | 0.243 | 0.784 |
| Year × Mix. | 14 | 4903.444 | 350.246 | 0.639 | 0.828 |
| Year × Irr. | 1 | 7091.138 | 7091.138 | 12.929 | 0.000 |
| Mix. × Irr. | 14 | 23144.35 | 1653.167 | 3.014 | 0.001 |
| Year × Mix. × Irr. | 14 | 5189.822 | 370.702 | 0.676 | 0.794 |
| Error | 118 | 64718.27 | 548.46 | - | - |
| % C.V. (coefficient of variation) |  | 3.58 |  |  |  |

**Table 2.**
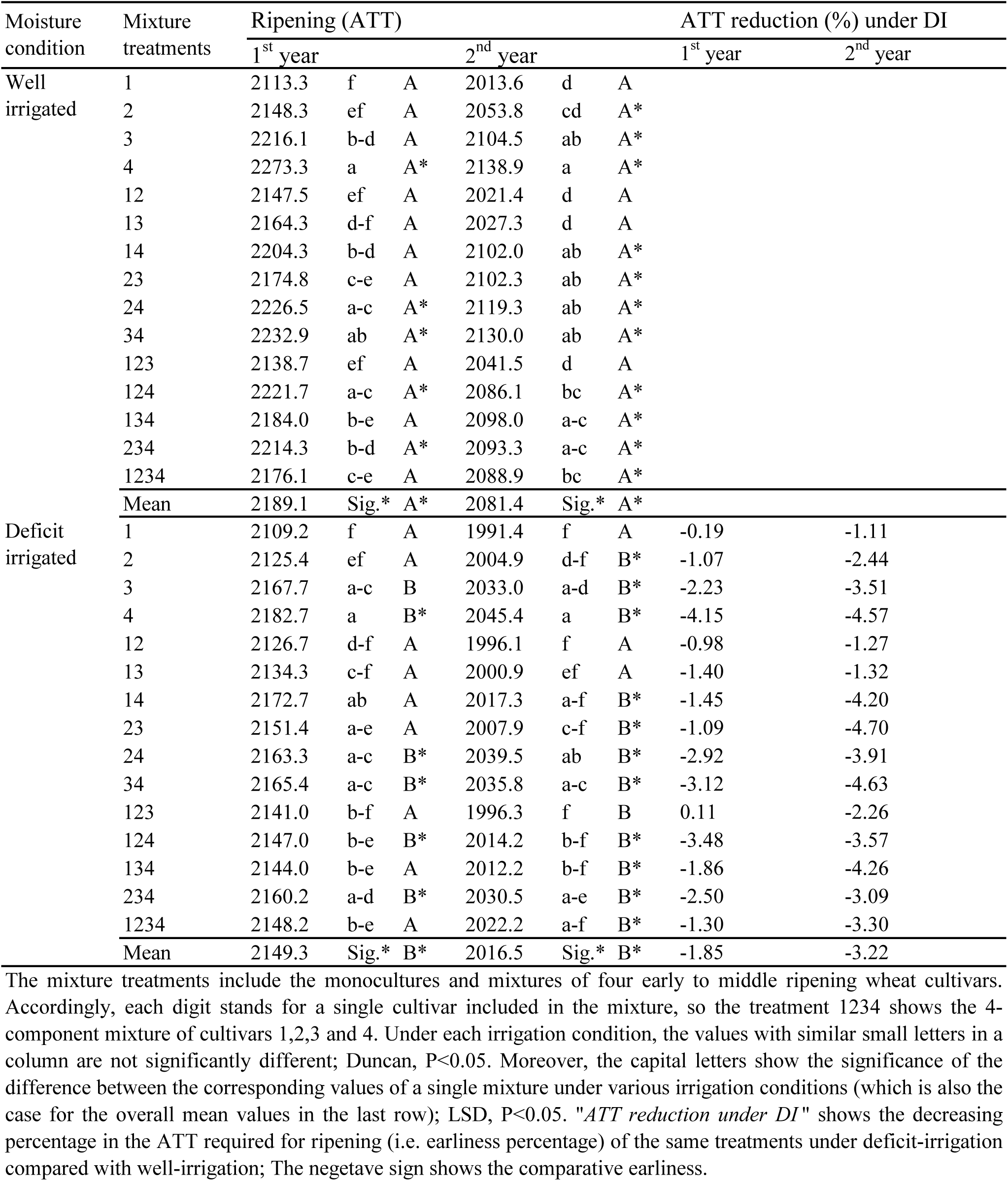
Mean comparison of accumulated thermal times required for ripening of mixture treatments under different irrigation treatments and years.

Moreover, although the effects of year and the interaction of mixture treatments × year on ripening was significant, some similar patterns were observed in the two seasons, i.e. the ripening of treatments 4, 24, 34, 124, and 234 (i.e. including the 4^th^ cultivar) occurred significantly earlier under deficit-irrigation conditions which is consistent with findings of Lopes *et al*., 2012; and Mwadzingeni *et al*., 2016, whereas in both years, the irrigation treatment had not significant effect on the ripening of treatments 1, 12, and 13. Although the accelerated ripening under deficit-irrigation was also observed in the remained treatments, the corresponding statistical significance were not similar in different seasons.

Besides, results showed that in both seasons, effect of deficit-irrigation on hastening of ripening in monocultures was enhanced from the earliest ripening cultivar (i.e. the 1^st^ cultivar) to the latest ripening ones (the 4^th^ cultivar). This might be associated with the stress avoidance mechanism in earliest ripening cultivar as has been noted by such researchers as Levitt, 1980, and Berger, 2016. These water stress-induced earliness were in average greater in the second year (%1.85 vs. %3.22). Also, it is notable that while in the first year the greatest stress-induced acceleration in ripening was observed in the monoculture of the 4^th^ cultivar, in the second year, the mixtures *23* and *34* were influenced more than this latest-ripened monoculture, and placed in the first rank. Such inconsistencies in the results are among the evidences for the considerable effect of season, even in the cultivar mixtures which are supposed to be more stable systems.

ADM analysis indicated that the range of deviation in the ripening of mixtures (compared with the corresponding monocultures) was wider under well-irrigation conditions in both seasons (Fig. 3). Moreover, averaged across the seasons and/or irrigation conditions, ripening was not markedly postponed or accelerated in mixtures compared with the corresponding monocultures (e.g. mixtures *14, 24, & 124* vs. *13 & 123*, respectively).

**Figure 3.**
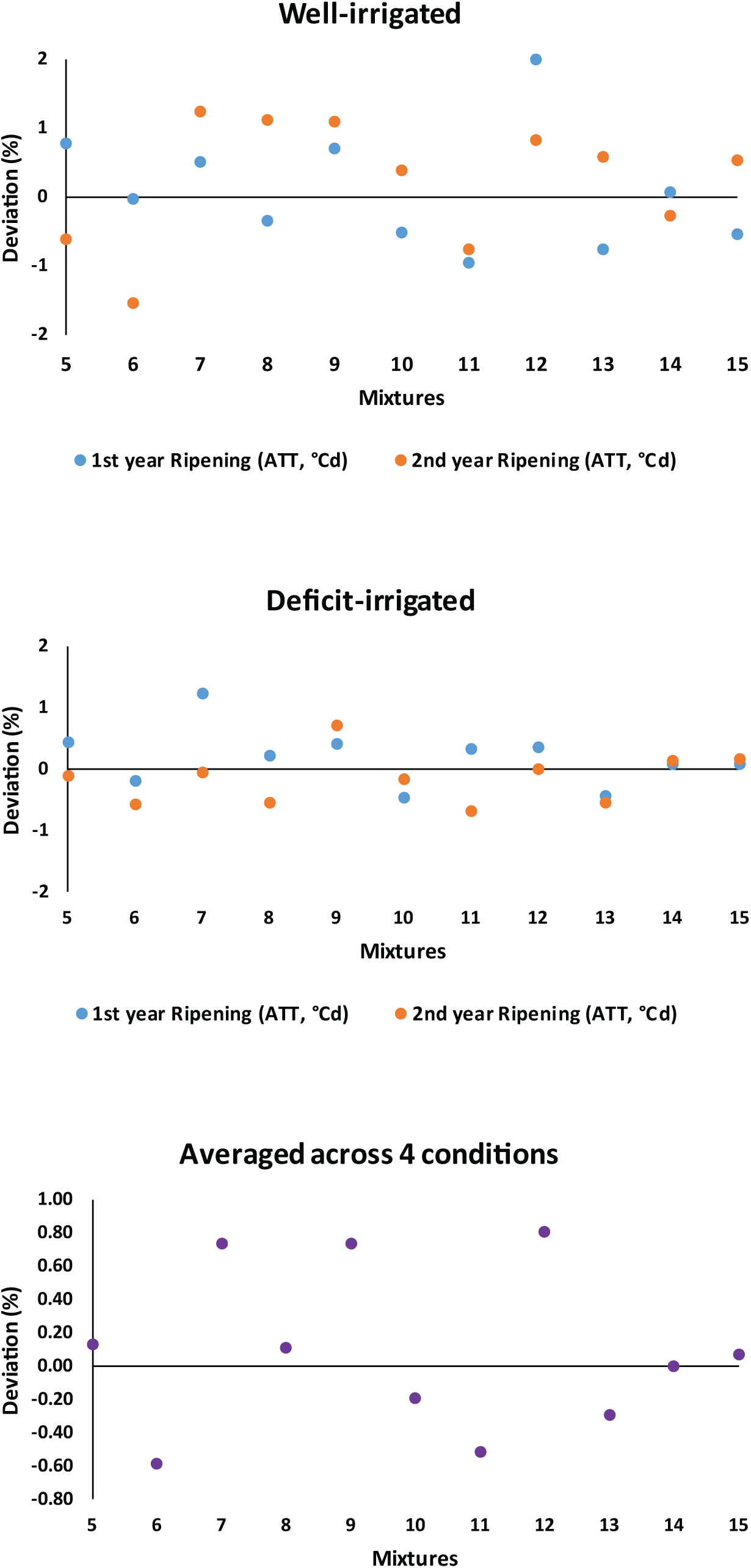
Deviation percentage of ripening (accumulated thermal time, ATT) of cultivar mixtures from mean of corresponding monocultures. Treatments 5 to 16 represent the mixtures of 12 (mixture of 1st and 2nd cultivars), 13, 14, 23, 24, 34, 123, 124, 134, 234, and 1234, respectively.

Orthogonal analysis (Tables 3 and S1) revealed that the ripening difference between mixtures and monocultures was not significant (see the comparison type contrast I). Indeed, the only considerable factors affected the ripening was inclusion or exclusion of the most early- and late-ripening cultivars (the 1^st^ vs. 4^th^ cultivars), regardless that the cropping system is mixtures or monoculture.

**Table 3.**
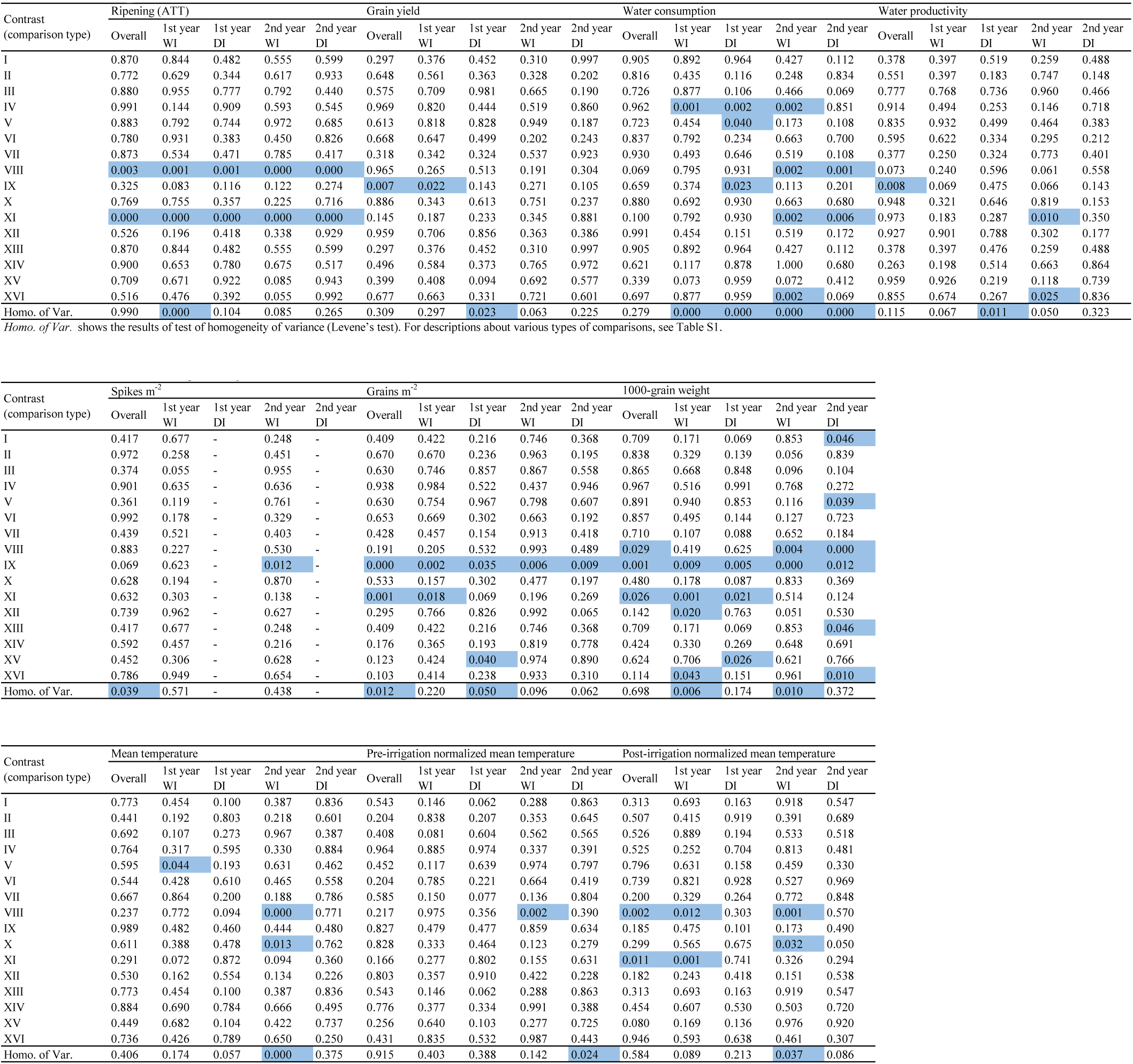
P-values of orthogonal comparisons for various traits.

Besides the quantitative image-based approach, the conventional phenology was also followed for comparing the relative growth and developmental phases of the four cultivars in monocultures. It was observed that the ATT requirement for various developmental phases also followed the similar order of 1^st^ < 2^nd^ < 3^rd^ < 4^th^ cultivar (e.g. see Fig. S2 which shows the comparative situation of early spike development in the second year).

### 3.2. Grain yield

As shown in Table 4, grain yield (GY) was significantly affected by year and irrigation, however, neither the effect of mixture treatments nor the interactions was statistically significant. Under the well-irrigation conditions in the first year, the monocultures of the earliest and latest ripening (i.e. the 1^st^ and 4^th^) cultivars produced the highest amounts of GY across the 15 mixture treatments. Indeed, these monocultures had significantly greater GY than the two other monocultures, and one of the two component mixtures (mixture *24*, Table 5). Grain yield in other mixtures were not statistically different with the maximum GY of monocultures. This result is consistent with the findings of other researchers who reported that the grain yield of wheat cultivar mixtures places in the rank between the highest and lowest yielding monocultures, and in many cases, there are some mixtures with statistically similar grain yield to the highest yielding monoculture (e.g. see Haghshenas *et al*., 2013; Fang *et al*., 2014; and Fletcher *et al*., 2019). However, under the well-irrigation condition in the second year, the situation was different, so that the 1^st^ cultivar had a significantly lower GY than the other three monocultures, which produced similar yields. Moreover, despite the first year, the highest and lowest grain yields were harvested from mixtures, rather than monocultures. Indeed, the mixture *34* had the highest GY and mixtures *23* and *24* had the lowest values, among which the latter had similarly the minimum GY in the 1^st^ year, as noted before.

**Table 4.** Analysis of variance of grain yield, yield components, water consumption, and water productivity.

| Source of variation | Grain yield |  |  |  |  | Water consumption |  |  |  | Water productivity |  |  |  |
| --- | --- | --- | --- | --- | --- | --- | --- | --- | --- | --- | --- | --- | --- |
|  | df | SS | MS | F | P-value | SS | MS | F | P-value | SS | MS | F | P-value |
| Year | 1 | 27.865 | 27.865 | 42.067 | 0.000 | 3733076.139 | 3733076 | 36.847 | 0.000 | 0.229 | 0.229 | 11.824 | 0.001 |
| Mixture treatments (Mix.) | 14 | 14.736 | 1.053 | 1.589 | 0.092 | 6287100.068 | 449078.6 | 4.433 | 0.000 | 0.375 | 0.027 | 1.383 | 0.172 |
| Irrigation (Irr.) | 1 | 46.144 | 46.144 | 69.662 | 0.000 | 90761384.25 | 90761384 | 895.845 | 0.000 | 0.223 | 0.223 | 11.505 | 0.001 |
| Replicate (Rep.) | 2 | 1.107 | 0.554 | 0.836 | 0.436 | 318370.496 | 159185.2 | 1.571 | 0.212 | 0.073 | 0.036 | 1.875 | 0.158 |
| Year × Mix. | 14 | 12.29 | 0.878 | 1.325 | 0.203 | 6452187.426 | 460870.5 | 4.549 | 0.000 | 0.346 | 0.025 | 1.275 | 0.233 |
| Year × Irr. | 1 | 0.146 | 0.146 | 0.221 | 0.639 | 7882015.084 | 7882015 | 77.798 | 0.000 | 0.124 | 0.124 | 6.412 | 0.013 |
| Mix. × Irr. | 14 | 5.862 | 0.419 | 0.632 | 0.834 | 4205767.804 | 300412 | 2.965 | 0.001 | 0.293 | 0.021 | 1.079 | 0.383 |
| Year × Mix. × Irr. | 14 | 7.712 | 0.551 | 0.832 | 0.634 | 2753000.206 | 196642.9 | 1.941 | 0.029 | 0.154 | 0.011 | 0.567 | 0.885 |
| Error | 118 | 78.162 | 0.662 | - | - | 11955018.17 | 101313.7 | - | - | 2.287 | 0.019 | - | - |
| % C.V. (coefficient of variation) |  | 16.72 |  |  |  | 14.28 |  |  |  | 14.65 |  |  |  |

Continue of Table 4. Analysis of variance of grain yield, yield components, water consumption, and water productivity.
| Source of variation | Spikes m<sup>-2</sup> |  |  |  |  | Grains m<sup>-2</sup> |  |  |  | 1000-grain weight |  |  |  |
| --- | --- | --- | --- | --- | --- | --- | --- | --- | --- | --- | --- | --- | --- |
|  | df | SS | MS | F | P-value | SS | MS | F | P-value | SS | MS | F | P-value |
| Year | 1 | 1022412 | 1022412 | 104.076 | 0.000 | 76127152.15 | 76127152 | 18.164 | 0.000 | 426.9111 | 426.9111 | 216.4481 | 1.83E-28 |
| Mixture treatments (Mix.) | 14 | 114023.5 | 8144.534 | 0.829 | 0.637 | 244100000 | 17436743 | 4.16 | 0.000 | 204.7702 | 14.62645 | 7.415752 | 6.56E-11 |
| Irrigation (Irr.) | - | - | - | - | - | 65509710.94 | 65509711 | 15.63 | 0.000 | 441.9535 | 441.9535 | 224.0748 | 4.82E-29 |
| Replicate (Rep.) | 5 | 59614.23 | 11922.85 | 1.214 | 0.306 | 529357.485 | 264678.7 | 0.063 | 0.939 | 13.08319 | 6.541597 | 3.316654 | 0.039685 |
| Year × Mix. | 14 | 212556.6 | 15182.62 | 1.546 | 0.102 | 94088368.14 | 6720598 | 1.604 | 0.088 | 57.61316 | 4.115226 | 2.08646 | 0.017137 |
| Year × Irr. | - | - | - | - | - | 36685.099 | 36685.1 | 0.009 | 0.926 | 4.171388 | 4.171388 | 2.114935 | 0.148523 |
| Mix. × Irr. | - | - | - | - | - | 37813676.36 | 2700977 | 0.644 | 0.823 | 62.08039 | 4.434313 | 2.24824 | 0.009587 |
| Year × Mix. × Irr. | - | - | - | - | - | 33900641.42 | 2421474 | 0.578 | 0.878 | 32.47142 | 2.319387 | 1.175952 | 0.302608 |
| Error | - | 1424434 | 9823.685 | - | - | 494600000 | 4191171 | - | - | 232.7371 | 1.972348 | - | - |
| % C.V. (coefficient of variation) |  | 16.81 |  |  |  | 14.87230311 |  |  |  | 7.611416 |  |  |  |

**Table 5.**
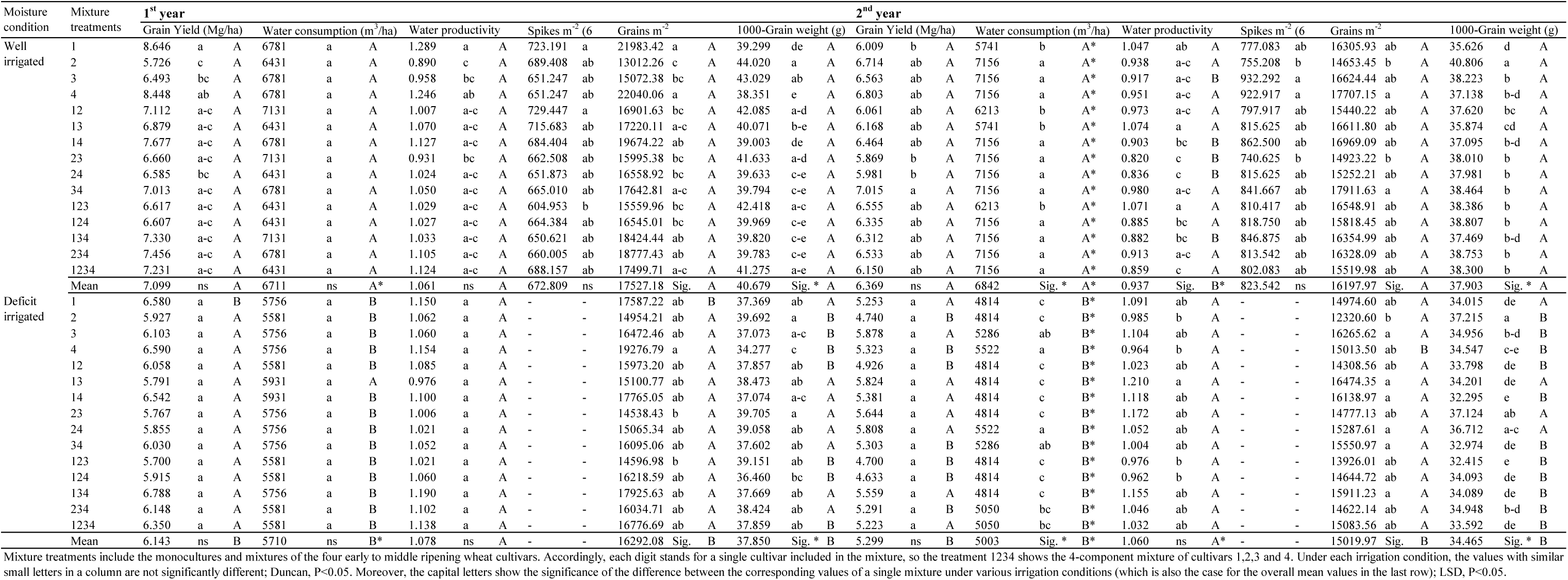
Mean comparison of grain yield, yield components, water consumption, and water productivity under different irrigation treatments and years.

Under deficit-irrigation conditions in both seasons, all monocultures and mixtures were in a single statistical group. Therefore, similar to the ripening pattern, the diversity in grain yield was considerably reduced under post-anthesis water stress. In average, deficit-irrigation reduced the GY 13.5% and 16.8% in the first and second year, respectively. Accordingly, the comparative average reductions in the GY of the monocultures and mixtures were 14.0% (mono.) vs. 13.2% (mix.), and 18.8% (mono.) vs. 16.1% (mix.) in the first and second years, respectively. Based on the susceptibility index (Table 6), grain yield in mixtures was in average about 9% more, and 15% less susceptible to the applied post-anthesis water stress compared with the monocultures, in the first and second years, respectively. This result was considerably similar to the same analysis obtained for the number of grains m^−2^.

**Table 6.**
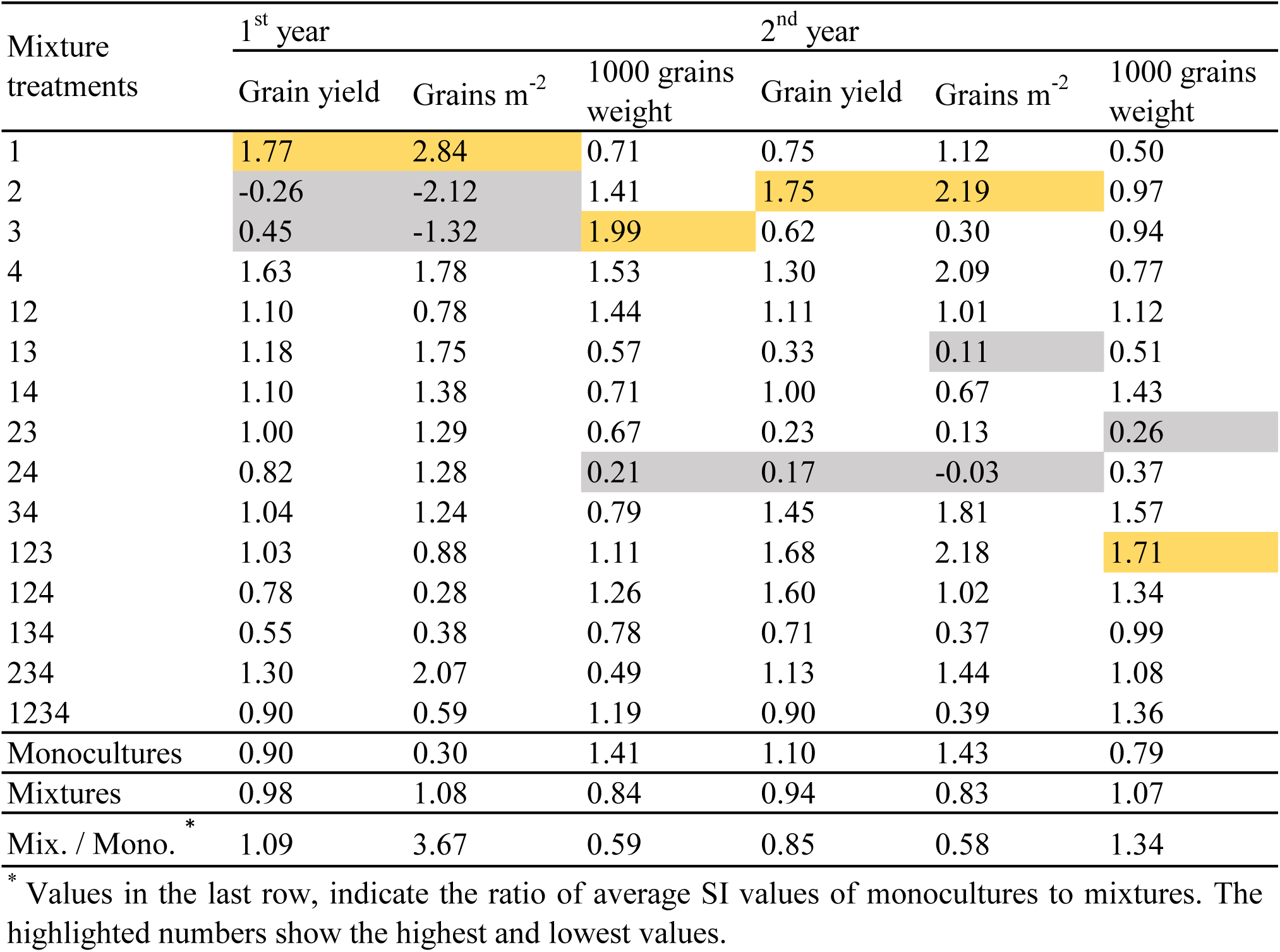
Values of Susceptibility Index (SI) calculated for grain yield and yield components.

ADM analysis showed that in most cases, GY of mixtures did not show any advantage compared to the monocultures (Fig. 4), and the rare comparative benefits gained from mixtures (e.g. the well-irrigated mixtures of *23* & *234* in the first year), did not remain stable and were declined under the other water stress condition or season. However, it seems that under deficit-irrigation, frequency of case similarities between the results of two years were relatively higher (e.g. see frequent adjacent blue and red circles in mixtures *12, 14, 34, 123, 124, 234, and 1234*, Fig. 4). Under each condition, deviations in GY of the mixtures from the corresponding monocultures were less than ±15%. Furthermore, averaged over the 4 environmental conditions (2 years and 2 irrigation treatments), 10 out of the total 11 mixtures had comparative GY less than their corresponding monoculture.

**Figure 4.**
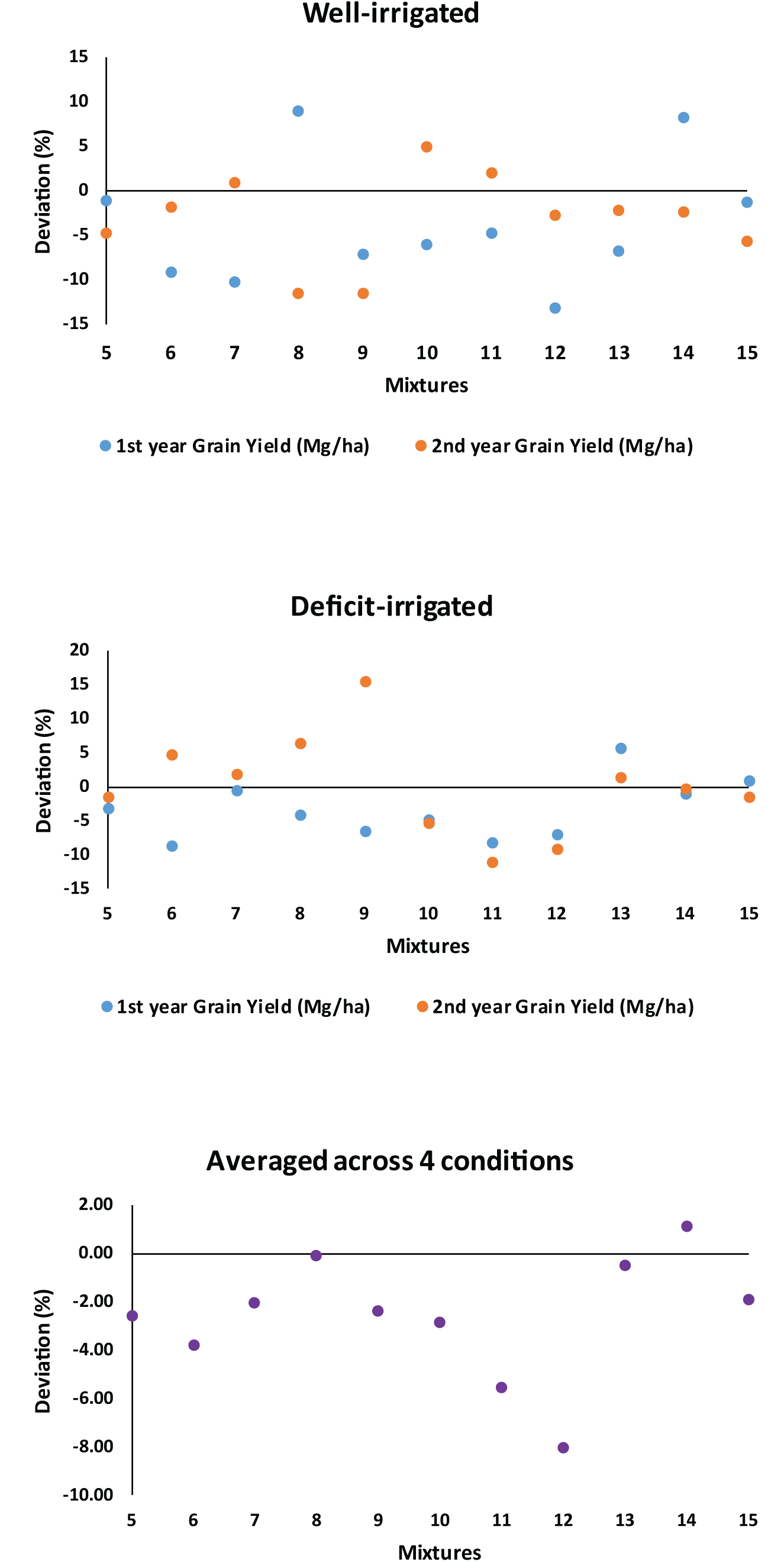
Deviation percentage of grain yield of cultivar mixtures from mean of corresponding monocultures. Treatments 5 to 16 represent the mixtures of 12 (mixture of 1st and 2nd cultivars), 13, 14, 23, 24, 34, 123, 124, 134, 234, and 1234, respectively.

Orthogonal analyses showed that there was not any significant difference between the grain yield of the two groups of monocultures and mixtures (Table 3, comparison type I). Among the various types of comparisons, the only significant difference was observed between the two sets of treatments i.e. those including vs. not-including 2^nd^ cultivar, particularly under well-irrigation condition of the first year, so that by exclusion of this cultivar, the grain yield was increased in average 10% (Table S1).

As shown in Table 7, averaged across the four conditions, all GY values ranged between the maximum and minimum extremes of the 4^th^ and 2^nd^ cultivars, respectively. Interestingly, C.V. of the highest yielding cultivars (the 4^th^ and the 1^st^ cultivars) was also maximum, which shows the comparatively higher degrees of instabilities in the grain yields of high-yielding monocultures. However, at the other side, monoculture of the 3^rd^ cultivar had the least C.V., and C.V. of the mixtures was placed at the next ranks. Similarly, based on CDI values, the first place in the list belonged to the 3^rd^ cultivar, which was followed by other mixtures (Table 7).

**Table 7.** Ranking of mixtures based on mean values, coefficient of variation (C.V.) and Consistent Desirable Index (CDI) of **measured characteristics, calculatedacross 4 environmental conditions.**

|  |  |  |  |  |  |  |
| --- | --- | --- | --- | --- | --- | --- |
| Ripening (ATT, °Cd) | Mixtures | Mean ATT | Mixtures | C.V. (%) | Mixtures | CDI index |
|  | 4 | 2160.08 | 1 | 3.08 | 4 | 206535.76 |
|  | 34 | 2141.05 | 2 | 3.16 | 34 | 205894.49 |
|  | 24 | 2137.15 | 1234 | 3.24 | 24 | 205858.72 |
|  | 3 | 2130.32 | 123 | 3.47 | 3 | 205099.14 |
|  | 234 | 2124.59 | 134 | 3.50 | 234 | 204471.82 |
|  | 14 | 2124.09 | 23 | 3.50 | 14 | 204104.07 |
|  | 124 | 2117.28 | 12 | 3.63 | 1234 | 204057.51 |
|  | 134 | 2109.56 | 24 | 3.68 | 134 | 203576.69 |
|  | 23 | 2109.11 | 3 | 3.72 | 23 | 203518.93 |
|  | 1234 | 2108.85 | 234 | 3.76 | 124 | 202900.09 |
|  | 2 | 2083.08 | 13 | 3.83 | 2 | 201720.55 |
|  | 13 | 2081.69 | 34 | 3.83 | 123 | 200715.88 |
|  | 123 | 2079.38 | 14 | 3.91 | 13 | 200192.28 |
|  | 12 | 2072.94 | 124 | 4.17 | 12 | 199761.54 |
|  | 1 | 2056.88 | 4 | 4.39 | 1 | 199342.95 |
| Grain yield (GY, Mg/ha) | Mixtures | Mean GY | Mixtures | C.V. (%) | Mixtures | CDI index |
|  | 4 | 6.79 | 3 | 5.19 | 3 | 593.42 |
|  | 1 | 6.62 | 24 | 5.93 | 134 | 574.62 |
|  | 14 | 6.52 | 23 | 7.67 | 24 | 569.80 |
|  | 134 | 6.50 | 13 | 8.20 | 13 | 566.02 |
|  | 234 | 6.36 | 134 | 11.56 | 14 | 557.80 |
|  | 34 | 6.34 | 34 | 13.13 | 23 | 552.56 |
|  | 3 | 6.26 | 1234 | 13.21 | 4 | 550.78 |
|  | 1234 | 6.24 | 2 | 14.06 | 34 | 550.77 |
|  | 13 | 6.17 | 234 | 14.12 | 234 | 545.93 |
|  | 24 | 6.06 | 14 | 14.39 | 1234 | 541.47 |
|  | 12 | 6.04 | 12 | 14.78 | 1 | 516.75 |
|  | 23 | 5.98 | 124 | 14.88 | 12 | 514.65 |
|  | 123 | 5.89 | 123 | 15.25 | 124 | 499.85 |
|  | 124 | 5.87 | 4 | 18.90 | 123 | 499.42 |
|  | 2 | 5.78 | 1 | 21.97 | 2 | 496.46 |
| Water productivity (WP) | Mixtures | Mean WP | Mixtures | C.V. (%) | Mixtures | CDI index |
|  | 1 | 1.14 | 34 | 3.47 | 1 | 103.89 |
|  | 13 | 1.08 | 123 | 3.80 | 13 | 98.62 |
|  | 4 | 1.08 | 12 | 4.59 | 34 | 98.60 |
|  | 134 | 1.07 | 2 | 7.56 | 123 | 98.53 |
|  | 14 | 1.06 | 124 | 7.86 | 12 | 97.51 |
|  | 234 | 1.04 | 3 | 8.61 | 14 | 95.54 |
|  | 1234 | 1.04 | 234 | 8.63 | 234 | 95.16 |
|  | 123 | 1.02 | 13 | 8.90 | 4 | 93.37 |
|  | 12 | 1.02 | 1 | 9.20 | 134 | 92.57 |
|  | 34 | 1.02 | 14 | 10.04 | 3 | 92.28 |
|  | 3 | 1.01 | 24 | 10.08 | 1234 | 90.98 |
|  | 124 | 0.98 | 1234 | 12.37 | 124 | 90.62 |
|  | 24 | 0.98 | 134 | 13.08 | 2 | 89.55 |
|  | 23 | 0.98 | 4 | 13.45 | 24 | 88.41 |
|  | 2 | 0.97 | 23 | 15.05 | 23 | 83.45 |
| Grains m<sup>-2</sup> (GN) | Mixtures | Mean GN | Mixtures | C.V. (%) | Mixtures | CDI index |
|  | 4 | 18509.37 | 23 | 4.28 | 14 | 1612500.39 |
|  | 1 | 17712.79 | 3 | 4.39 | 134 | 1594406.65 |
|  | 14 | 17636.83 | 24 | 4.41 | 34 | 1564497.19 |
|  | 134 | 17154.07 | 124 | 5.25 | 4 | 1557002.07 |
|  | 34 | 16800.12 | 13 | 5.47 | 13 | 1545701.97 |
|  | 234 | 16440.59 | 1234 | 6.87 | 3 | 1540235.15 |
|  | 13 | 16351.76 | 34 | 6.88 | 1234 | 1510507.20 |
|  | 1234 | 16219.98 | 12 | 6.91 | 124 | 1497701.67 |
|  | 3 | 16108.73 | 134 | 7.05 | 24 | 1485544.84 |
|  | 124 | 15806.69 | 123 | 7.55 | 234 | 1471384.32 |
|  | 12 | 15655.90 | 14 | 8.57 | 1 | 1467245.57 |
|  | 24 | 15541.02 | 2 | 9.26 | 12 | 1457356.11 |
|  | 123 | 15157.97 | 234 | 10.50 | 23 | 1441416.21 |
|  | 23 | 15058.54 | 4 | 15.88 | 123 | 1401359.54 |
|  | 2 | 13735.13 | 1 | 17.16 | 2 | 1246326.37 |
| 1000-grain weight (GW, g) | Mixtures | Mean GW | Mixtures | C.V. (%) | Mixtures | CDI index |
|  | 2 | 40.43 | 24 | 3.36 | 2 | 3761.01 |
|  | 23 | 39.12 | 23 | 5.09 | 23 | 3712.86 |
|  | 24 | 38.35 | 4 | 5.52 | 24 | 3705.93 |
|  | 3 | 38.32 | 234 | 5.53 | 234 | 3587.63 |
|  | 123 | 38.09 | 1 | 6.22 | 3 | 3490.19 |
|  | 234 | 37.98 | 134 | 6.35 | 134 | 3489.39 |
|  | 12 | 37.84 | 124 | 6.98 | 124 | 3472.58 |
|  | 1234 | 37.76 | 2 | 6.98 | 1234 | 3459.27 |
|  | 124 | 37.33 | 13 | 7.05 | 13 | 3453.39 |
|  | 134 | 37.26 | 14 | 7.87 | 12 | 3445.34 |
|  | 34 | 37.21 | 34 | 7.96 | 1 | 3430.39 |
|  | 13 | 37.15 | 1234 | 8.38 | 34 | 3424.50 |
|  | 1 | 36.58 | 3 | 8.92 | 4 | 3408.85 |
|  | 14 | 36.37 | 12 | 8.95 | 123 | 3392.31 |
|  | 4 | 36.08 | 123 | 10.95 | 14 | 3350.55 |

The slopes of color bars indicate the increasing or decreasing trends. ATT: accumulated thermal time (°Cd).

### 3.3. Grain yield components

The strongest correlation between GY and its components was observed with GN (number of grains m^−2^; Table 8), which was the most important contributor to GY, irrespective of the year or irrigation condition. This result is in accordance with our current knowledge about the main role of number of grains per unit of area (particularly compared with 1000-grain weight) in determining the final grain yield (e.g. see Slafer *et al*., 2014). In the first year, there was also a relatively considerable negative relationship between GN and 1000 GW (Table 8). Slafer *et al*. (2014) have reviewed the mechanisms beyond this negative relationship, and believed that with increasing grain number m^−2^, there is an increase in the proportion of grains of lower potential weight in distal positions within the spikelets, in secondary tiller spikes, or both. Hence a negative relationship between the number and size arises that does not necessarily involve competition for resources among grains. In general, number of spikes m^−2^ had not a considerable effect on GY, which suggest that the dependence of GY to GN is probably originating from the number of grains per spike, rather than the number of spikes m^−2^.

**Table 8.** Correlation among estimated properties under various conditions. (A) under well-irrigation condition in the 1st year.

| Variables | GY | W | WP | Spikes m<sup>-2</sup> | Grains m<sup>-2</sup> | 1000 GW | ATT | SWC 0-30 | SWC 30-60 | SWC 0-60 | LSWC 0-30 | LSWC 30-60 | LSWC 0-60 | T | Pre T | Post T |
| --- | --- | --- | --- | --- | --- | --- | --- | --- | --- | --- | --- | --- | --- | --- | --- | --- |
| Grain Yield (GY) | 1 | 0.336 | 0.937 | 0.237 | 0.984 | -0.743 | 0.104 | -0.241 | 0.070 | -0.081 | -0.241 | 0.070 | -0.081 | -0.234 | -0.178 | -0.312 |
| Water consumption (W) | 0.336 | 1 | -0.009 | 0.131 | 0.286 | -0.102 | -0.004 | -0.269 | -0.004 | -0.144 | -0.269 | -0.004 | -0.144 | -0.383 | -0.488 | -0.273 |
| Water productivity (WP) | 0.937 | -0.009 | 1 | 0.216 | 0.937 | -0.749 | 0.100 | -0.152 | 0.075 | -0.031 | -0.152 | 0.075 | -0.031 | -0.111 | -0.010 | -0.232 |
| Spikes m<sup>-2</sup> | 0.237 | 0.131 | 0.216 | 1 | 0.199 | -0.045 | -0.425 | -0.002 | 0.074 | 0.047 | -0.002 | 0.074 | 0.047 | 0.365 | 0.379 | 0.286 |
| Grains m<sup>-2</sup> | 0.984 | 0.286 | 0.937 | 0.199 | 1 | -0.847 | 0.214 | -0.261 | 0.079 | -0.086 | -0.261 | 0.079 | -0.086 | -0.185 | -0.136 | -0.338 |
| 1000-grain weight (1000 GW) | -0.743 | -0.102 | -0.749 | -0.045 | -0.847 | 1 | -0.470 | 0.231 | -0.117 | 0.045 | 0.231 | -0.117 | 0.045 | 0.043 | 0.042 | 0.355 |
| Ripening (ATT) | 0.104 | -0.004 | 0.100 | -0.425 | 0.214 | -0.470 | 1 | -0.165 | 0.326 | 0.127 | -0.165 | 0.326 | 0.127 | -0.291 | -0.190 | -0.793 |
| Soil water content 0-30 cm | -0.241 | -0.269 | -0.152 | -0.002 | -0.261 | 0.231 | -0.165 | 1 | 0.415 | 0.801 | 1.000 | 0.415 | 0.801 | -0.160 | -0.077 | 0.091 |
| Soil water content 30-60 cm | 0.070 | -0.004 | 0.075 | 0.074 | 0.079 | -0.117 | 0.326 | 0.415 | 1 | 0.877 | 0.415 | 1.000 | 0.877 | -0.224 | 0.083 | -0.516 |
| Soil water content 0-60 cm | -0.081 | -0.144 | -0.031 | 0.047 | -0.086 | 0.045 | 0.127 | 0.801 | 0.877 | 1 | 0.801 | 0.877 | 1.000 | -0.232 | 0.014 | -0.292 |
| Last sampling SWC 0-30 cm | -0.241 | -0.269 | -0.152 | -0.002 | -0.261 | 0.231 | -0.165 | 1.000 | 0.415 | 0.801 | 1 | 0.415 | 0.801 | -0.160 | -0.077 | 0.091 |
| Last sampling SWC 30-60 cm | 0.070 | -0.004 | 0.075 | 0.074 | 0.079 | -0.117 | 0.326 | 0.415 | 1.000 | 0.877 | 0.415 | 1 | 0.877 | -0.224 | 0.083 | -0.516 |
| Last sampling SWC 0-60 cm | -0.081 | -0.144 | -0.031 | 0.047 | -0.086 | 0.045 | 0.127 | 0.801 | 0.877 | 1.000 | 0.801 | 0.877 | 1 | -0.232 | 0.014 | -0.292 |
| Canopy temperature (T) | -0.234 | -0.383 | -0.111 | 0.365 | -0.185 | 0.043 | -0.291 | -0.160 | -0.224 | -0.232 | -0.160 | -0.224 | -0.232 | 1 | 0.895 | 0.645 |
| Pre-irrigation temperature (Pre T) | -0.178 | -0.488 | -0.010 | 0.379 | -0.136 | 0.042 | -0.190 | -0.077 | 0.083 | 0.014 | -0.077 | 0.083 | 0.014 | 0.895 | 1 | 0.419 |
| Post-irrigation temperature (Post T) | -0.312 | -0.273 | -0.232 | 0.286 | -0.338 | 0.355 | -0.793 | 0.091 | -0.516 | -0.292 | 0.091 | -0.516 | -0.292 | 0.645 | 0.419 | 1 |

ATT: accumulated thermal time, SWC: soil water content, LSWC: soil water content at the last sampling. Highlighted numbers show the considerable correlations.

| Variables | GY | W | WP | Grains m<sup>-2</sup> | 1000 GW | ATT | SWC 0-30 | SWC 30-60 | SWC 0-60 | LSWC 0-30 | LSWC 30-60 | LSWC 0-60 | T | Pre T | Post T |
| --- | --- | --- | --- | --- | --- | --- | --- | --- | --- | --- | --- | --- | --- | --- | --- |
| Grain Yield (GY) | 1 | 0.237 | 0.931 | 0.926 | -0.610 | 0.128 | -0.258 | 0.052 | -0.123 | -0.258 | 0.052 | -0.123 | -0.336 | -0.407 | -0.388 |
| Water consumption (W) | 0.237 | 1 | -0.130 | 0.256 | -0.183 | 0.282 | -0.024 | 0.239 | 0.111 | -0.024 | 0.239 | 0.111 | -0.031 | -0.059 | -0.163 |
| Water productivity (WP) | 0.931 | -0.130 | 1 | 0.848 | -0.553 | 0.020 | -0.280 | -0.064 | -0.197 | -0.280 | -0.064 | -0.197 | -0.327 | -0.405 | -0.326 |
| Grains m<sup>-2</sup> | 0.926 | 0.256 | 0.848 | 1 | -0.860 | 0.314 | -0.106 | 0.097 | -0.011 | -0.106 | 0.097 | -0.011 | -0.317 | -0.337 | -0.470 |
| 1000-grain weight (1000 GW) | -0.610 | -0.183 | -0.553 | -0.860 | 1 | -0.437 | -0.133 | -0.172 | -0.168 | -0.133 | -0.172 | -0.168 | 0.211 | 0.135 | 0.502 |
| Ripening (ATT) | 0.128 | 0.282 | 0.020 | 0.314 | -0.437 | 1 | 0.068 | 0.004 | 0.042 | 0.068 | 0.004 | 0.042 | -0.303 | -0.045 | -0.279 |
| Soil water content 0-30 cm | -0.258 | -0.024 | -0.280 | -0.106 | -0.133 | 0.068 | 1 | 0.630 | 0.913 | 1.000 | 0.630 | 0.913 | -0.047 | 0.597 | -0.380 |
| Soil water content 30-60 cm | 0.052 | 0.239 | -0.064 | 0.097 | -0.172 | 0.004 | 0.630 | 1 | 0.891 | 0.630 | 1.000 | 0.891 | 0.149 | 0.506 | -0.111 |
| Soil water content 0-60 cm | -0.123 | 0.111 | -0.197 | -0.011 | -0.168 | 0.042 | 0.913 | 0.891 | 1 | 0.913 | 0.891 | 1.000 | 0.051 | 0.613 | -0.280 |
| Last sampling SWC 0-30 cm | -0.258 | -0.024 | -0.280 | -0.106 | -0.133 | 0.068 | 1.000 | 0.630 | 0.913 | 1 | 0.630 | 0.913 | -0.047 | 0.597 | -0.380 |
| Last sampling SWC 30-60 cm | 0.052 | 0.239 | -0.064 | 0.097 | -0.172 | 0.004 | 0.630 | 1.000 | 0.891 | 0.630 | 1 | 0.891 | 0.149 | 0.506 | -0.111 |
| Last sampling SWC 0-60 cm | -0.123 | 0.111 | -0.197 | -0.011 | -0.168 | 0.042 | 0.913 | 0.891 | 1.000 | 0.913 | 0.891 | 1 | 0.051 | 0.613 | -0.280 |
| Canopy temperature (T) | -0.336 | -0.031 | -0.327 | -0.317 | 0.211 | -0.303 | -0.047 | 0.149 | 0.051 | -0.047 | 0.149 | 0.051 | 1 | 0.604 | 0.723 |
| Pre-irrigation temperature (Pre T) | -0.407 | -0.059 | -0.405 | -0.337 | 0.135 | -0.045 | 0.597 | 0.506 | 0.613 | 0.597 | 0.506 | 0.613 | 0.604 | 1 | 0.124 |
| Post-irrigation temperature (Post T) | -0.388 | -0.163 | -0.326 | -0.470 | 0.502 | -0.279 | -0.380 | -0.111 | -0.280 | -0.380 | -0.111 | -0.280 | 0.723 | 0.124 | 1 |

ATT: accumulated thermal time, SWC: soil water content, LSWC: soil water content at the last sampling. Highlighted numbers show the considerable correlations.

| Variables | GY | W | WP | Spikes m<sup>-2</sup> | Grains m<sup>-2</sup> | 1000 GW | ATT | SWC 0-30 | SWC 30-60 | SWC 0-60 | LSWC 0-30 | LSWC 30-60 | LSWC 0-60 | T | Pre T | Post T |
| --- | --- | --- | --- | --- | --- | --- | --- | --- | --- | --- | --- | --- | --- | --- | --- | --- |
| Grain Yield (GY) | 1 | 0.352 | 0.240 | 0.511 | 0.624 | 0.412 | 0.395 | -0.433 | -0.445 | -0.550 | -0.433 | -0.445 | -0.550 | -0.302 | -0.273 | -0.300 |
| Water consumption (W) | 0.352 | 1 | -0.823 | 0.276 | -0.049 | 0.613 | 0.858 | -0.084 | -0.237 | -0.220 | -0.084 | -0.237 | -0.220 | -0.754 | -0.713 | -0.716 |
| Water productivity (WP) | 0.240 | -0.823 | 1 | 0.019 | 0.428 | -0.381 | -0.644 | -0.178 | -0.006 | -0.093 | -0.178 | -0.006 | -0.093 | 0.587 | 0.562 | 0.548 |
| Spikes m<sup>-2</sup> | 0.511 | 0.276 | 0.019 | 1 | 0.717 | -0.194 | 0.521 | -0.446 | -0.427 | -0.543 | -0.446 | -0.427 | -0.543 | -0.321 | -0.297 | -0.311 |
| Grains m<sup>-2</sup> | 0.624 | -0.049 | 0.428 | 0.717 | 1 | -0.406 | 0.331 | -0.426 | -0.351 | -0.476 | -0.426 | -0.351 | -0.476 | 0.013 | -0.023 | 0.051 |
| 1000-grain weight (1000 GW) | 0.412 | 0.613 | -0.381 | -0.194 | -0.406 | 1 | 0.246 | 0.007 | -0.074 | -0.052 | 0.007 | -0.074 | -0.052 | -0.530 | -0.428 | -0.584 |
| Ripening (ATT) | 0.395 | 0.858 | -0.644 | 0.521 | 0.331 | 0.246 | 1 | -0.019 | -0.228 | -0.180 | -0.019 | -0.228 | -0.180 | -0.743 | -0.727 | -0.676 |
| Soil water content 0-30 cm | -0.433 | -0.084 | -0.178 | -0.446 | -0.426 | 0.007 | -0.019 | 1 | 0.251 | 0.688 | 1.000 | 0.251 | 0.688 | -0.112 | -0.097 | -0.116 |
| Soil water content 30-60 cm | -0.445 | -0.237 | -0.006 | -0.427 | -0.351 | -0.074 | -0.228 | 0.251 | 1 | 0.875 | 0.251 | 1.000 | 0.875 | 0.021 | -0.155 | 0.214 |
| Soil water content 0-60 cm | -0.550 | -0.220 | -0.093 | -0.543 | -0.476 | -0.052 | -0.180 | 0.688 | 0.875 | 1 | 0.688 | 0.875 | 1.000 | -0.040 | -0.165 | 0.103 |
| Last sampling SWC 0-30 cm | -0.433 | -0.084 | -0.178 | -0.446 | -0.426 | 0.007 | -0.019 | 1.000 | 0.251 | 0.688 | 1 | 0.251 | 0.688 | -0.112 | -0.097 | -0.116 |
| Last sampling SWC 30-60 cm | -0.445 | -0.237 | -0.006 | -0.427 | -0.351 | -0.074 | -0.228 | 0.251 | 1.000 | 0.875 | 0.251 | 1 | 0.875 | 0.021 | -0.155 | 0.214 |
| Last sampling SWC 0-60 cm | -0.550 | -0.220 | -0.093 | -0.543 | -0.476 | -0.052 | -0.180 | 0.688 | 0.875 | 1.000 | 0.688 | 0.875 | 1 | -0.040 | -0.165 | 0.103 |
| Canopy temperature (T) | -0.302 | -0.754 | 0.587 | -0.321 | 0.013 | -0.530 | -0.743 | -0.112 | 0.021 | -0.040 | -0.112 | 0.021 | -0.040 | 1 | 0.952 | 0.941 |
| Pre-irrigation temperature (Pre T) | -0.273 | -0.713 | 0.562 | -0.297 | -0.023 | -0.428 | -0.727 | -0.097 | -0.155 | -0.165 | -0.097 | -0.155 | -0.165 | 0.952 | 1 | 0.791 |
| Post-irrigation temperature (Post T) | -0.300 | -0.716 | 0.548 | -0.311 | 0.051 | -0.584 | -0.676 | -0.116 | 0.214 | 0.103 | -0.116 | 0.214 | 0.103 | 0.941 | 0.791 | 1 |

ATT: accumulated thermal time, SWC: soil water content, LSWC: soil water content at the last sampling. Highlighted numbers show the considerable correlations.

| Variables | GY | W | WP | Grains m<sup>-2</sup> | 1000 GW | ATT | SWC 0-30 | SWC 30-60 | SWC 0-60 | LSWC 0-30 | LSWC 30-60 | LSWC 0-60 | T | Pre T | Post T |
| --- | --- | --- | --- | --- | --- | --- | --- | --- | --- | --- | --- | --- | --- | --- | --- |
| Grain Yield (GY) | 1 | 0.410 | 0.749 | 0.759 | 0.251 | 0.388 | 0.021 | -0.343 | -0.312 | 0.021 | -0.343 | -0.312 | -0.313 | 0.130 | -0.473 |
| Water consumption (W) | 0.410 | 1 | -0.294 | 0.238 | 0.186 | 0.892 | -0.140 | -0.288 | -0.355 | -0.140 | -0.288 | -0.355 | 0.035 | 0.341 | -0.279 |
| Water productivity (WP) | 0.749 | -0.294 | 1 | 0.624 | 0.122 | -0.239 | 0.160 | -0.155 | -0.053 | 0.160 | -0.155 | -0.053 | -0.342 | -0.122 | -0.271 |
| Grains m<sup>-2</sup> | 0.759 | 0.238 | 0.624 | 1 | -0.362 | 0.308 | 0.068 | -0.203 | -0.153 | 0.068 | -0.203 | -0.153 | -0.506 | -0.064 | -0.510 |
| 1000-grain weight (1000 GW) | 0.251 | 0.186 | 0.122 | -0.362 | 1 | 0.139 | 0.206 | -0.407 | -0.264 | 0.206 | -0.407 | -0.264 | -0.007 | -0.023 | 0.013 |
| Ripening (ATT) | 0.388 | 0.892 | -0.239 | 0.308 | 0.139 | 1 | -0.125 | -0.498 | -0.545 | -0.125 | -0.498 | -0.545 | -0.218 | 0.244 | -0.473 |
| Soil water content 0-30 cm | 0.021 | -0.140 | 0.160 | 0.068 | 0.206 | -0.125 | 1 | -0.218 | 0.381 | 1.000 | -0.218 | 0.381 | -0.168 | -0.544 | 0.319 |
| Soil water content 30-60 cm | -0.343 | -0.288 | -0.155 | -0.203 | -0.407 | -0.498 | -0.218 | 1 | 0.819 | -0.218 | 1.000 | 0.819 | 0.497 | -0.093 | 0.647 |
| Soil water content 0-60 cm | -0.312 | -0.355 | -0.053 | -0.153 | -0.264 | -0.545 | 0.381 | 0.819 | 1 | 0.381 | 0.819 | 1.000 | 0.372 | -0.408 | 0.800 |
| Last sampling SWC 0-30 cm | 0.021 | -0.140 | 0.160 | 0.068 | 0.206 | -0.125 | 1.000 | -0.218 | 0.381 | 1 | -0.218 | 0.381 | -0.168 | -0.544 | 0.319 |
| Last sampling SWC 30-60 cm | -0.343 | -0.288 | -0.155 | -0.203 | -0.407 | -0.498 | -0.218 | 1.000 | 0.819 | -0.218 | 1 | 0.819 | 0.497 | -0.093 | 0.647 |
| Last sampling SWC 0-60 cm | -0.312 | -0.355 | -0.053 | -0.153 | -0.264 | -0.545 | 0.381 | 0.819 | 1.000 | 0.381 | 0.819 | 1 | 0.372 | -0.408 | 0.800 |
| Canopy temperature (T) | -0.313 | 0.035 | -0.342 | -0.506 | -0.007 | -0.218 | -0.168 | 0.497 | 0.372 | -0.168 | 0.497 | 0.372 | 1 | 0.542 | 0.619 |
| Pre-irrigation temperature (Pre T) | 0.130 | 0.341 | -0.122 | -0.064 | -0.023 | 0.244 | -0.544 | -0.093 | -0.408 | -0.544 | -0.093 | -0.408 | 0.542 | 1 | -0.324 |
| Post-irrigation temperature (Post T) | -0.473 | -0.279 | -0.271 | -0.510 | 0.013 | -0.473 | 0.319 | 0.647 | 0.800 | 0.319 | 0.647 | 0.800 | 0.619 | -0.324 | 1 |

ATT: accumulated thermal time, SWC: soil water content, LSWC: soil water content at the last sampling. Highlighted numbers show the considerable correlations.

Continue of Table 8. Correlation among estimated properties averaged across the 4 conditions.
| Variables | GY | W | WP | Spikes m<sup>-2</sup> | Grains m<sup>-2</sup> | 1000 GW | ATT | SWC 0-30 | SWC 30-60 | SWC 0-60 | LSWC 0-30 | LSWC 30-60 | LSWC 0-60 | T | Pre T | Post T |
| --- | --- | --- | --- | --- | --- | --- | --- | --- | --- | --- | --- | --- | --- | --- | --- | --- |
| Grain Yield (GY) | 1 | 0.709 | 0.316 | -0.364 | 0.862 | 0.574 | 0.687 | 0.380 | 0.164 | 0.304 | 0.380 | 0.164 | 0.304 | -0.646 | -0.494 | -0.657 |
| Water consumption (W) | 0.709 | 1 | -0.439 | -0.071 | 0.404 | 0.681 | 0.569 | 0.463 | 0.411 | 0.509 | 0.463 | 0.411 | 0.509 | -0.399 | -0.473 | -0.561 |
| Water productivity (WP) | 0.316 | -0.439 | 1 | -0.348 | 0.553 | -0.198 | 0.086 | -0.153 | -0.343 | -0.304 | -0.153 | -0.343 | -0.304 | -0.264 | 0.003 | -0.078 |
| Spikes m<sup>-2</sup> | -0.364 | -0.071 | -0.348 | 1 | -0.261 | -0.460 | -0.642 | -0.286 | 0.243 | 0.012 | -0.286 | 0.243 | 0.012 | 0.785 | 0.341 | 0.437 |
| Grains m<sup>-2</sup> | 0.862 | 0.404 | 0.553 | -0.261 | 1 | 0.116 | 0.519 | 0.164 | 0.061 | 0.125 | 0.164 | 0.061 | 0.125 | -0.477 | -0.364 | -0.550 |
| 1000-grain weight (1000 GW) | 0.574 | 0.681 | -0.198 | -0.460 | 0.116 | 1 | 0.671 | 0.514 | 0.077 | 0.316 | 0.514 | 0.077 | 0.316 | -0.684 | -0.423 | -0.444 |
| Ripening (ATT) | 0.687 | 0.569 | 0.086 | -0.642 | 0.519 | 0.671 | 1 | 0.386 | -0.071 | 0.153 | 0.386 | -0.071 | 0.153 | -0.893 | -0.454 | -0.715 |
| Soil water content 0-30 cm | 0.380 | 0.463 | -0.153 | -0.286 | 0.164 | 0.514 | 0.386 | 1 | 0.445 | 0.809 | 1.000 | 0.445 | 0.809 | -0.407 | -0.288 | -0.428 |
| Soil water content 30-60 cm | 0.164 | 0.411 | -0.343 | 0.243 | 0.061 | 0.077 | -0.071 | 0.445 | 1 | 0.886 | 0.445 | 1.000 | 0.886 | 0.179 | -0.094 | -0.215 |
| Soil water content 0-60 cm | 0.304 | 0.509 | -0.304 | 0.012 | 0.125 | 0.316 | 0.153 | 0.809 | 0.886 | 1 | 0.809 | 0.886 | 1.000 | -0.093 | -0.210 | -0.363 |
| Last sampling SWC 0-30 cm | 0.380 | 0.463 | -0.153 | -0.286 | 0.164 | 0.514 | 0.386 | 1.000 | 0.445 | 0.809 | 1 | 0.445 | 0.809 | -0.407 | -0.288 | -0.428 |
| Last sampling SWC 30-60 cm | 0.164 | 0.411 | -0.343 | 0.243 | 0.061 | 0.077 | -0.071 | 0.445 | 1.000 | 0.886 | 0.445 | 1 | 0.886 | 0.179 | -0.094 | -0.215 |
| Last sampling SWC 0-60 cm | 0.304 | 0.509 | -0.304 | 0.012 | 0.125 | 0.316 | 0.153 | 0.809 | 0.886 | 1.000 | 0.809 | 0.886 | 1 | -0.093 | -0.210 | -0.363 |
| Canopy temperature (T) | -0.646 | -0.399 | -0.264 | 0.785 | -0.477 | -0.684 | -0.893 | -0.407 | 0.179 | -0.093 | -0.407 | 0.179 | -0.093 | 1 | 0.576 | 0.676 |
| Pre-irrigation temperature (Pre T) | -0.494 | -0.473 | 0.003 | 0.341 | -0.364 | -0.423 | -0.454 | -0.288 | -0.094 | -0.210 | -0.288 | -0.094 | -0.210 | 0.576 | 1 | 0.531 |
| Post-irrigation temperature (Post T) | -0.657 | -0.561 | -0.078 | 0.437 | -0.550 | -0.444 | -0.715 | -0.428 | -0.215 | -0.363 | -0.428 | -0.215 | -0.363 | 0.676 | 0.531 | 1 |

ATT: accumulated thermal time, SWC: soil water content, LSWC: soil water content at the last sampling. Highlithed numbers show the considerable correlations.

Number of grains m^−2^ was very significantly affected by year, mixture treatment, and irrigation (Table 4). Post-anthesis deficit-irrigation had reduced the average GN by 7.05% and 7.23% in the first and second years, respectively (Table 5). The 2^nd^ cultivar had produced the lowest GN among the monocultures under every 4 conditions (2 seasons and 2 irrigation treatments), which can explain its less GY. Among the consistent cases in the mixtures were treatments 23 and 14 which had produced low and high amounts of GN under all 4 conditions, similar to GY. Result of ADM analysis for GN was differed considerably between the two seasons (Fig. 5); for instance, deficit-irrigated mixtures had generally produced higher GN in the 2^nd^ year, compared with the average of their corresponding monocultures; while the trend was opposite in the first year. Averaged across the 4 conditions, mixtures 23 and 234 had a slightly (1.3% and 2%, respectively) more GN compared with their corresponding monocultures. Based on the orthogonal comparisons, there was no significant difference between GN values of monocultures and mixtures (comparison type I), and almost similar to GY (which described before), the only meaningful contrast was referred to inclusion or exclusion of the low GN producing component i.e. the 2^nd^ cultivar (Table S1). Results of CDI was almost similar to those described for GY (Table 7), except that the first three treatments with the highest CDI values in the list were three mixtures (excluding the 2^nd^ cultivar). It is among the rare results that indicates a relatively advantage of mixtures compared with monocultures. Furthermore, based on susceptibility index, average GN of mixtures had influenced by the deficit-irrigation about 3.67 fold higher than monocultures in the first year; however, in the second year, mixtures showed 42% less susceptibility to water stress compared with monocultures. It might be another evidence for instability in the results.

**Figure 5.**
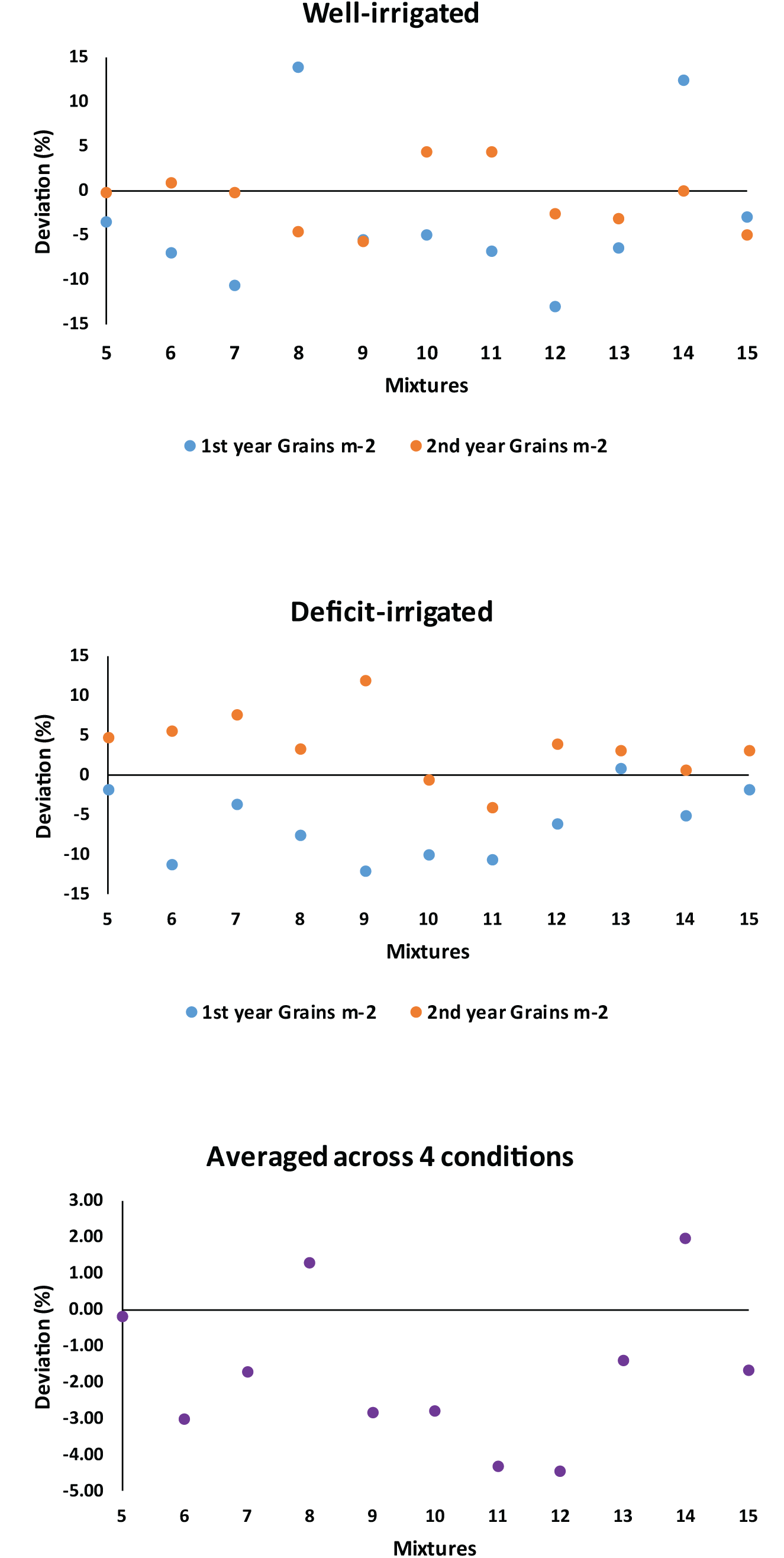
Deviation percentage of number of grains per m2 of cultivar mixtures from mean of corresponding monocultures. Treatments 5 to 16 represent the mixtures of 12 (mixture of 1st and 2nd cultivars), 13, 14, 23, 24, 34, 123, 124, 134, 234, and 1234, respectively.

Number of spikes m^−2^ was affected only by the season (Table 4), as in average it increased 22.4% in the second year (24.8% in mixtures and 21.5% in monocultures). This difference may be associated with the comparatively longer tillering period or probably more desirable conditions during this phase in the second year. ADM analysis indicated that in general, mixed cropping of cultivars had reduced number of spikes m^−2^ compared with the corresponding monocultures, irrespective of the season (at most about 12%, Fig. 6). The only exception was the mixture 12 which produced 3.28% and 4.15% higher spikes m^−2^ in the first and second seasons, respectively, compared with its corresponding monocultures. Orthogonal analysis revealed that there was no significant difference between mixtures and monocultures in number of spikes m^−2^ (Table 3, comparison type I). Besides, no other considerable difference was observed, except for a comparison between inclusion and exclusion of the 2^nd^ cultivars in the second year, as produced 7.9% more spikes m^−2^ in the absence of this cultivar in the canopy (Table S1).

**Figure 6.**
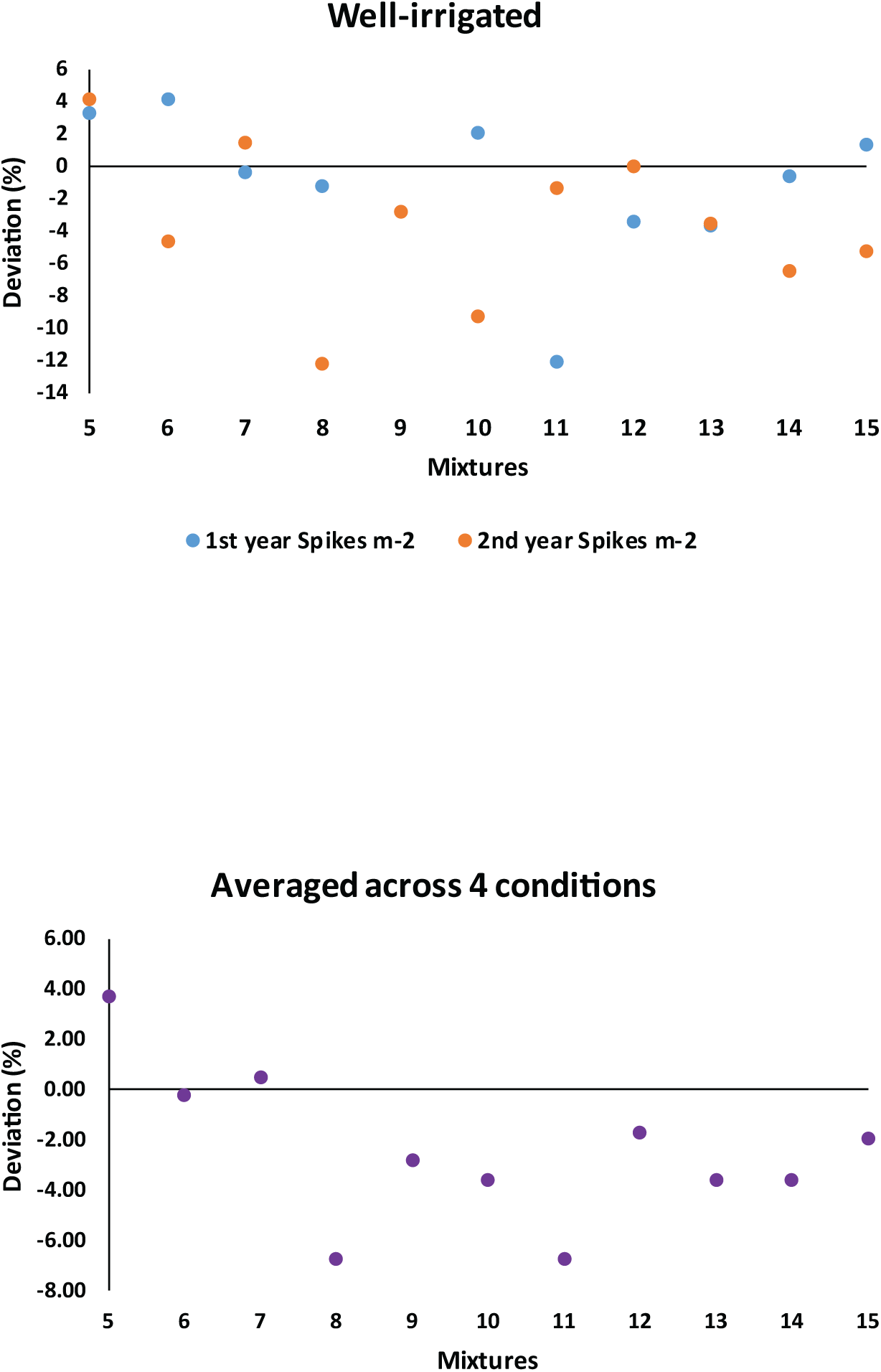
Deviation percentage of number of spikes per m2 of cultivar mixtures from mean of corresponding monocultures. Treatments 5 to 16 represent the mixtures of 12 (mixture of 1st and 2nd cultivars), 13, 14, 23, 24, 34, 123, 124, 134, 234, and 1234, respectively.

In the present study, 1000-grain weight had relatively less contribution to the grain yield and as noted before, it had a strong negative relationship with GN. Effects of year, mixture treatment, irrigation, and the interactions of Year × Mixture and Mixture × Irrigation on this trait were significant (Table 4). Post-anthesis deficit-irrigation reduced 1000-grain weight 7.0% and 9.1% in the first and second year, respectively. The 1000-GW in monoculture of the 1^st^ cultivar along with the mixtures 13, 23, and 24 was not affected significantly by deficit-irrigation, while it was significantly reduced in all other treatments (Table 5). Regardless of the year and irrigation treatment, 1000-GW was significantly lower in the 1^st^ and 4^th^ cultivars compared with other cultivars. This was the reverse of the trend found in GY. Under deficit-irrigation in both seasons, mixtures 23 and 24 had relatively higher 1000-GWs; however, particularly under well-irrigation condition, there was not any other mixture with stable high amounts of this trait. The deviation percentages of 1000-GW of mixtures from their corresponding monocultures ranged between −4.8% to +2.51% under well-irrigation, and between −8.4% to +5.6% under deficit-irrigation condition (Fig. 7). When averaged across the four conditions, this range narrowed, i.e. was between −1.85% to +0.75%. Although similar to other traits, there were inconsistent results across the years, yet there were some consistent cases among them. For instance, mixtures 23 and 24 had lower 1000-GWs compared with their corresponding monocultures under well-irrigation conditions in both seasons; however, they produced comparatively higher 1000-grain weights under deficit-irrigation (Fig. 7). As shown in Table 6, exactly opposite to the GN, the ratio of mixture/ monoculture SI was lower and higher in the first and second seasons, respectively. However, the extreme values were observed in different treatments in each season. Orthogonal comparisons indicated that there was no difference between average 1000-GW of monocultures and mixtures (Table 3), except under deficit-irrigation condition of the second year; where monocultures had higher values of 1000-grain weight. Although some other significant differences were observed through orthogonal comparisons, mostly they were not consistent across all conditions; however, in an overall view, despite the 1^st^ and 4^th^ cultivars, the inclusion of the 2^nd^ cultivar in the canopy, significantly improved 1000-GW (Table S1). Similarly, based on CDI (Table 7), the 2^nd^ cultivar and its corresponding mixtures included it had the highest and most stable values of 1000-GW; while the high yielding monocultures of the 1^st^ and 4^th^ cultivars placed at the second half of the list.

**Figure 7.**
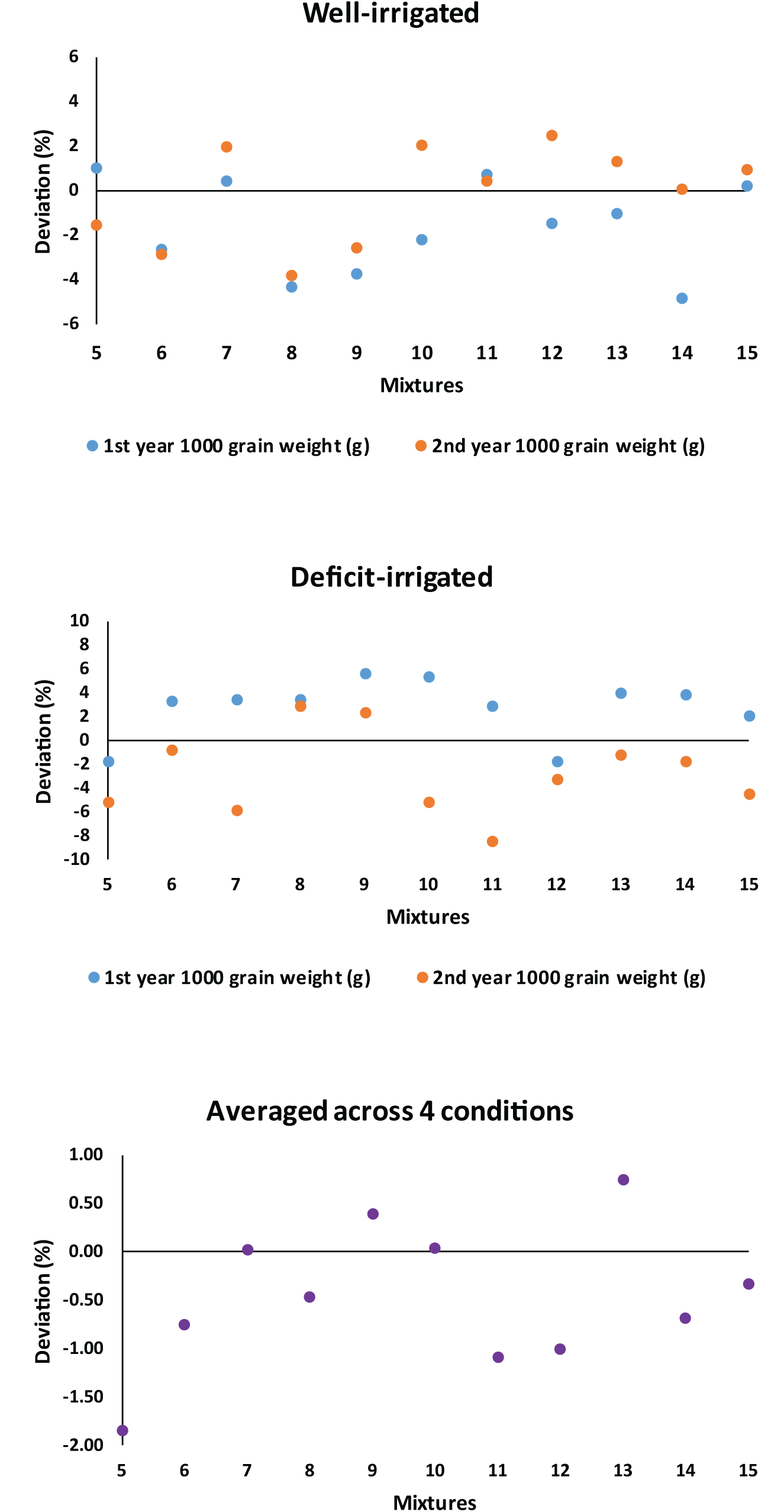
Deviation percentage of 1000-grain weight of cultivar mixtures from mean of corresponding monocultures. Treatments 5 to 16 represent the mixtures of 12 (mixture of 1st and 2nd cultivars), 13, 14, 23, 24, 34, 123, 124, 134, 234, and 1234, respectively.

### 3.4. Water consumption, water productivity, and water use efficiency

Effects of year, mixture treatment, irrigation, and their dual interactions on water consumption (i.e. the summation of effective rainfall and irrigation water) were significant (Table 4). Based on the designed treatments, amount of water consumption was expected to be affected by two main factors, i.e. amount of water estimated based on irrigation treatment, and taking the last irrigation into the account depended on the degree of canopy ripening. Under deficit-irrigation, water consumption was declined significantly by 15% and 27% in the first and second years, respectively (Table 5). Such reduction for monocultures were equal to 14.7% and 24.9% against 15% and 27.6% in the mixtures, in the first and second years, respectively. Based on the comparison type I of the orthogonal analyses (Tables 3 and S1), there was no difference between water consumption of the monocultures and mixtures. Meaningful results of other orthogonal comparisons were also unstable across the four conditions. For instance, under both irrigation conditions of the 1^st^ year, the 4-component mixture (i.e. treatment 1234) on average consumed significantly lower water compared with other mixtures and monocultures, while in the second year, the condition was the reverse. As another example, in the second year, the treatments included the earliest ripening cultivar (i.e. the 1^st^ one) had comparatively consumed less water than others; despite for the presence of the latest ripening cultivar (the 4^th^ one) in the canopy which required higher water volume, particularly due to the estimations of the last irrigation in the season. However, as noted before, these trends were limited to only one season, and it is difficult to extrapolate or recommend such results for other conditions and climates.

Water productivity (WP) was significantly affected by year, irrigation treatment, and their interaction (Table 4). In the first year, there was no significant difference between the WP of the well- and deficit-irrigated treatments (Table 5), while in the second year, it was increased significantly by about 13% under deficit-irrigation condition. As shown in Fig. 8, deviation of WP of the mixtures from their corresponding monocultures ranged between ±10% (under well-irrigation) and −7.5% to 6.8% (under deficit-irrigation), among which mixtures 134 and 24 had a consistent increase in both years, under well- and deficit-irrigation conditions, respectively. Besides, averaged across the four conditions, none of the mixtures had WP higher than 2% compared with the respective monocultures, while in the case of comparative reductions, there were mixtures (i.e. treatments 13 and 123) with a slightly larger deviation (more than 4%) compared with monocultures. Orthogonal analyses showed that there was not any significant difference between monocultures and mixtures (Table 3, comparison type I). Overall across the four conditions, inclusion of the 2^nd^ cultivar had significantly reduced WP, but this effect was not observed in individual comparisons within each year. Furthermore, only under well-irrigation condition in the second year, water productivity was significantly and negatively influenced by presence of the 4^th^ cultivar, and also maximum extension in the duration of single phenological events in the canopies (i.e. simultaneously inclusion of the 1^st^ and 4^th^ cultivars in mixtures). Averaged across the 4 conditions, the maximum and minimum WP values was found in the 1^st^ and 2^nd^ cultivars, respectively (Table 7), and other treatments placed between these extremums. The highest CDI for water productivity was also observed in the 1^st^ cultivars, which followed by mixtures 13, 34, and 123. On the other hand, mixture 23 had the least WP. Based on result of Principal Component Analysis, variations in water productivity was much more associated with GY than water consumption, irrespective of the environmental condition (data not shown).

**Figure 8.**
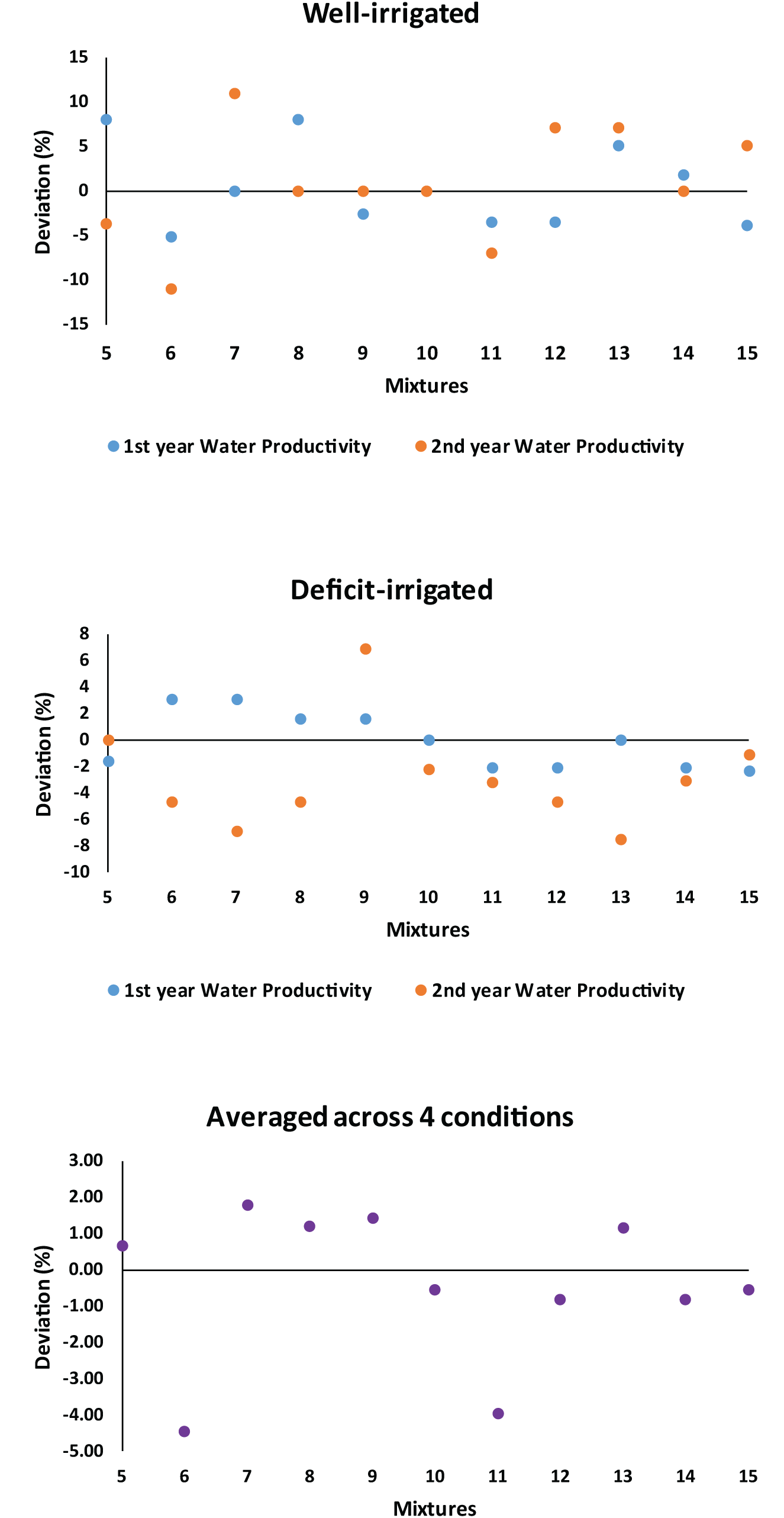
Deviation percentage of water productivity of cultivar mixtures from mean of corresponding monocultures. Treatments 5 to 16 represent the mixtures of 12 (mixture of 1st and 2nd cultivars), 13, 14, 23, 24, 34, 123, 124, 134, 234, and 1234, respectively.

Although due to the limitations in the number of replication and the depth of soil water sampling (see M&M) the focus of water demand analyses was put on the model-based *ET_C_* estimation and WP, from the physiological point of view, *ET_C_*′ and WUE might reflect more actual status of water extraction and productivity in the canopy; due to including the actual (sampled) soil water content in the calculations. Results of *ET_C_*′ and WUE analyses are reported briefly (see supplementary Tables S2, S3, S4 and Fig. S4 & S5). As a whole, mixture treatment did not make any difference neither in *ET*_*C*_′ nor in WUE (Tables S3 and S4, contrast I). Moreover, ADM analysis showed that averaged across the 4 conditions, WUE was lowered in the majority of mixtures compared with their corresponding monocultures (Fig. S5). As a notable instance of detrimental effect of mixtures, water use efficiency of the mixture 12 was consistently lower than the average of the respective monocultures, under all the conditions.

### 3.5. Canopy temperature

As shown in Table 9, effects of year, mixture treatment, and irrigation treatment on the triple canopy temperatures (i.e. mean, pre-irrigation, and post-irrigation temperatures) were significant, except for the effect of mixture treatment on the pre-irrigation canopy temperature. Excluding the pre-irrigation temperature in the 2^nd^ year, deficit-irrigation increased canopy temperature significantly (Table 10), which is in accordance with the results reported by Blum *et* al, 1982; Blad *et* al., 1988; and Gontia & Tiwari, 2008. Under deficit-irrigation conditions in both seasons, pre-irrigation canopy temperatures were statistically similar; however, post-irrigation temperatures were significantly different and heterogeneous. This finding which was consistent between the two seasons, may be another evidence for reduction of heterogeneity in crop performance under stressful conditions, and suggest that it is under desirable conditions (i.e. after irrigation) that the potential differences among mixture treatments are detectable. Under well-irrigation condition, pre-irrigation canopy temperatures had also significantly diverse patterns, regardless of the year (Table 10).

**Table 9.**
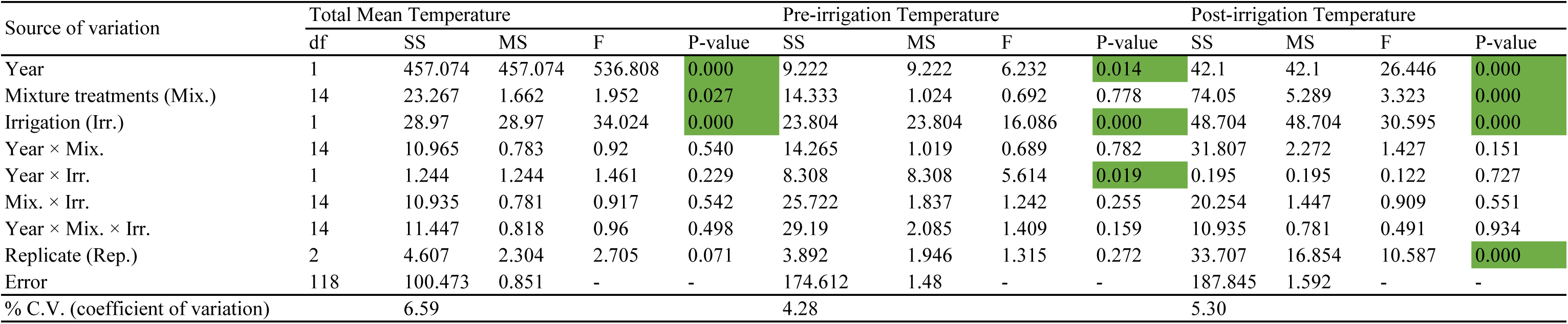
Analysis of variance of canopy temperature during post-anthesis period of two years.

| Source of variation | Total Mean Temperature |  |  |  |  | Pre-irrigation Temperature |  |  |  | Post-irrigation Temperature |  |  |  |
| --- | --- | --- | --- | --- | --- | --- | --- | --- | --- | --- | --- | --- | --- |
|  | df | SS | MS | F | P-value | SS | MS | F | P-value | SS | MS | F | P-value |
| Year | 1 | 457.074 | 457.074 | 536.808 | 0.000 | 9.222 | 9.222 | 6.232 | 0.014 | 42.1 | 42.1 | 26.446 | 0.000 |
| Mixture treatments (Mix.) | 14 | 23.267 | 1.662 | 1.952 | 0.027 | 14.333 | 1.024 | 0.692 | 0.778 | 74.05 | 5.289 | 3.323 | 0.000 |
| Irrigation (Irr.) | 1 | 28.97 | 28.97 | 34.024 | 0.000 | 23.804 | 23.804 | 16.086 | 0.000 | 48.704 | 48.704 | 30.595 | 0.000 |
| Year × Mix. | 14 | 10.965 | 0.783 | 0.92 | 0.540 | 14.265 | 1.019 | 0.689 | 0.782 | 31.807 | 2.272 | 1.427 | 0.151 |
| Year × Irr. | 1 | 1.244 | 1.244 | 1.461 | 0.229 | 8.308 | 8.308 | 5.614 | 0.019 | 0.195 | 0.195 | 0.122 | 0.727 |
| Mix. × Irr. | 14 | 10.935 | 0.781 | 0.917 | 0.542 | 25.722 | 1.837 | 1.242 | 0.255 | 20.254 | 1.447 | 0.909 | 0.551 |
| Year × Mix. × Irr. | 14 | 11.447 | 0.818 | 0.96 | 0.498 | 29.19 | 2.085 | 1.409 | 0.159 | 10.935 | 0.781 | 0.491 | 0.934 |
| Replicate (Rep.) | 2 | 4.607 | 2.304 | 2.705 | 0.071 | 3.892 | 1.946 | 1.315 | 0.272 | 33.707 | 16.854 | 10.587 | 0.000 |
| Error | 118 | 100.473 | 0.851 | - | - | 174.612 | 1.48 | - | - | 187.845 | 1.592 | - | - |
| % C.V. (coefficient of variation) |  | 6.59 |  |  |  | 4.28 |  |  |  | 5.30 |  |  |  |

**Table 10.**
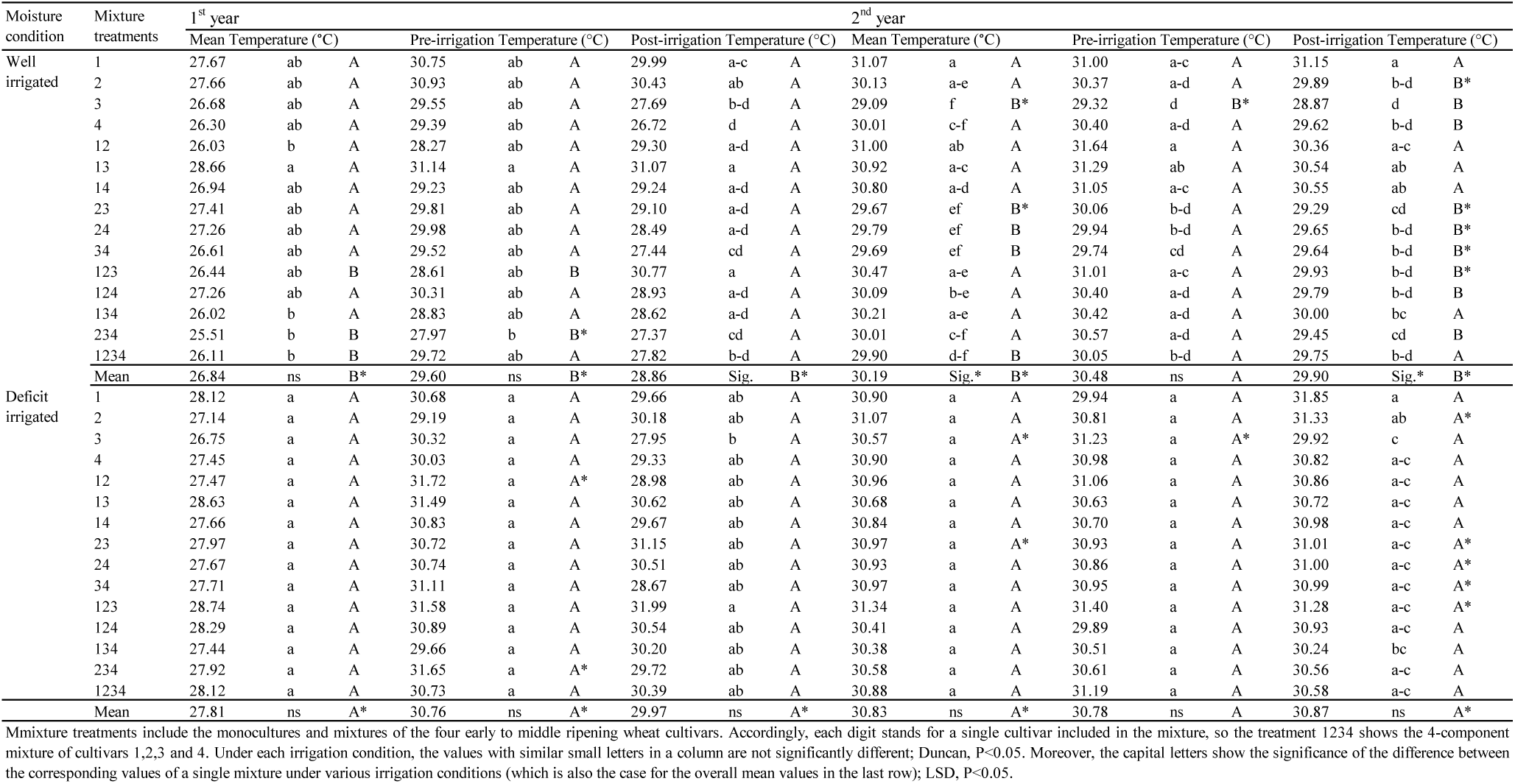
Mean comparison of canopy temperatures recorded under well- and deficit-irrigation during two years.

As there were negative correlations between GY and the triplet canopy temperatures (e.g. see R values of correlations estimated as average of the 4 conditions, Table 8), the relatively lower temperatures were more focused in the analyses. As shown in Fig. 9 mixtures 13 & 134 had higher and lower canopy mean temperatures, respectively, compared with their corresponding monocultures. Such trend was also observed for pre-irrigation temperature (Fig. 10). However, after irrigation, the trend was completely different, and post-irrigation canopy temperatures of mixtures were generally higher (up to 1.1 °C) than their corresponding monocultures (Fig. 11). The only exception which had a comparatively lower post-irrigation canopy temperature, was mixture 12 (i.e. had a cooler canopy around 0.7 °C). These results suggest that probably the canopy of monocultures have a higher degrees of transpiration under the post-irrigation stressless situations, compared with mixtures, e.g. due to better aerodynamic conditions, root system distribution, or other associated factors.

**Figure 9.**
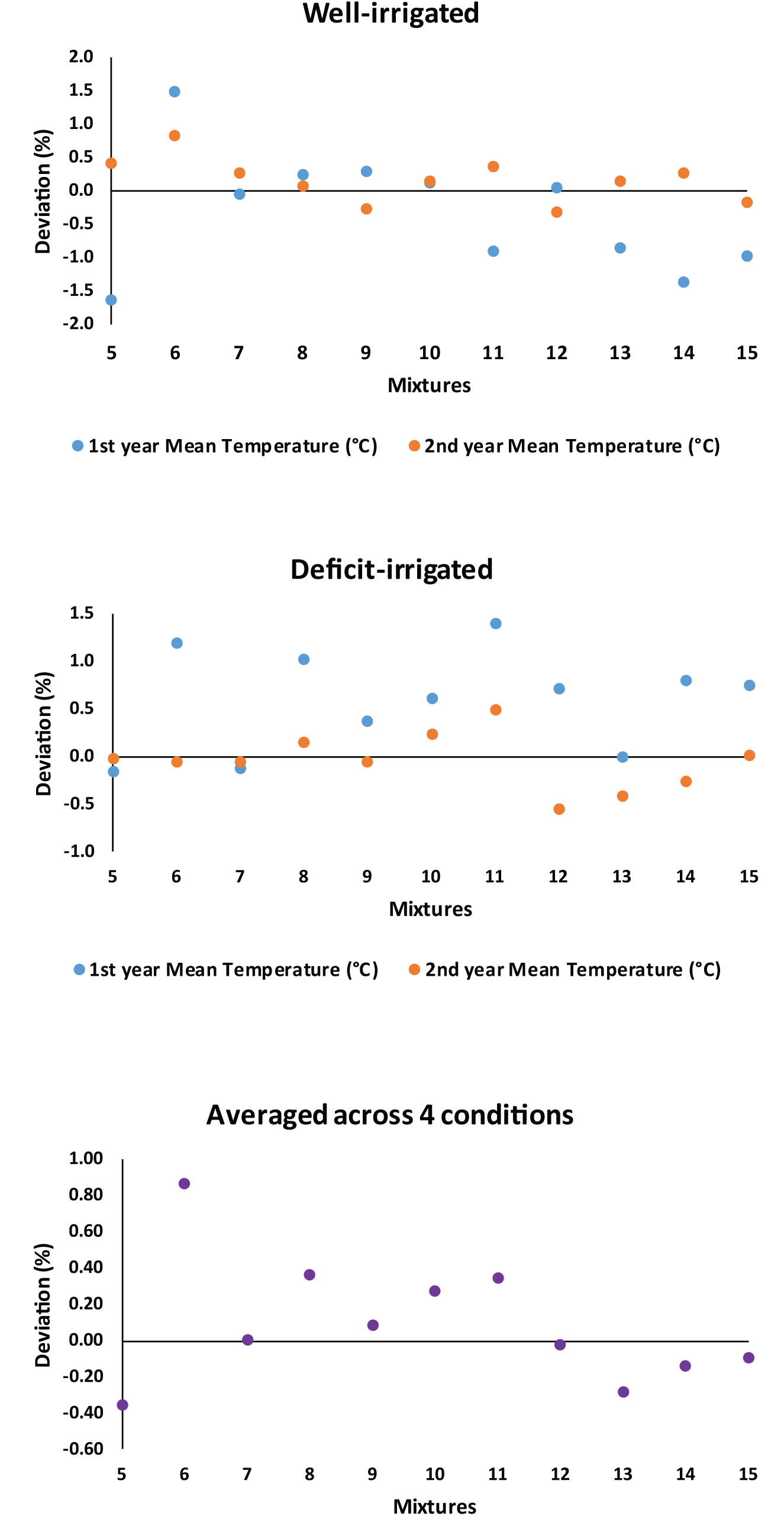
Deviation percentage of mean canopy temperature of cultivar mixtures from mean of corresponding monocultures. Treatments 5 to 16 represent the mixtures of 12 (mixture of 1st and 2nd cultivars), 13, 14, 23, 24, 34, 123, 124, 134, 234, and 1234, respectively.

**Figure 10.**
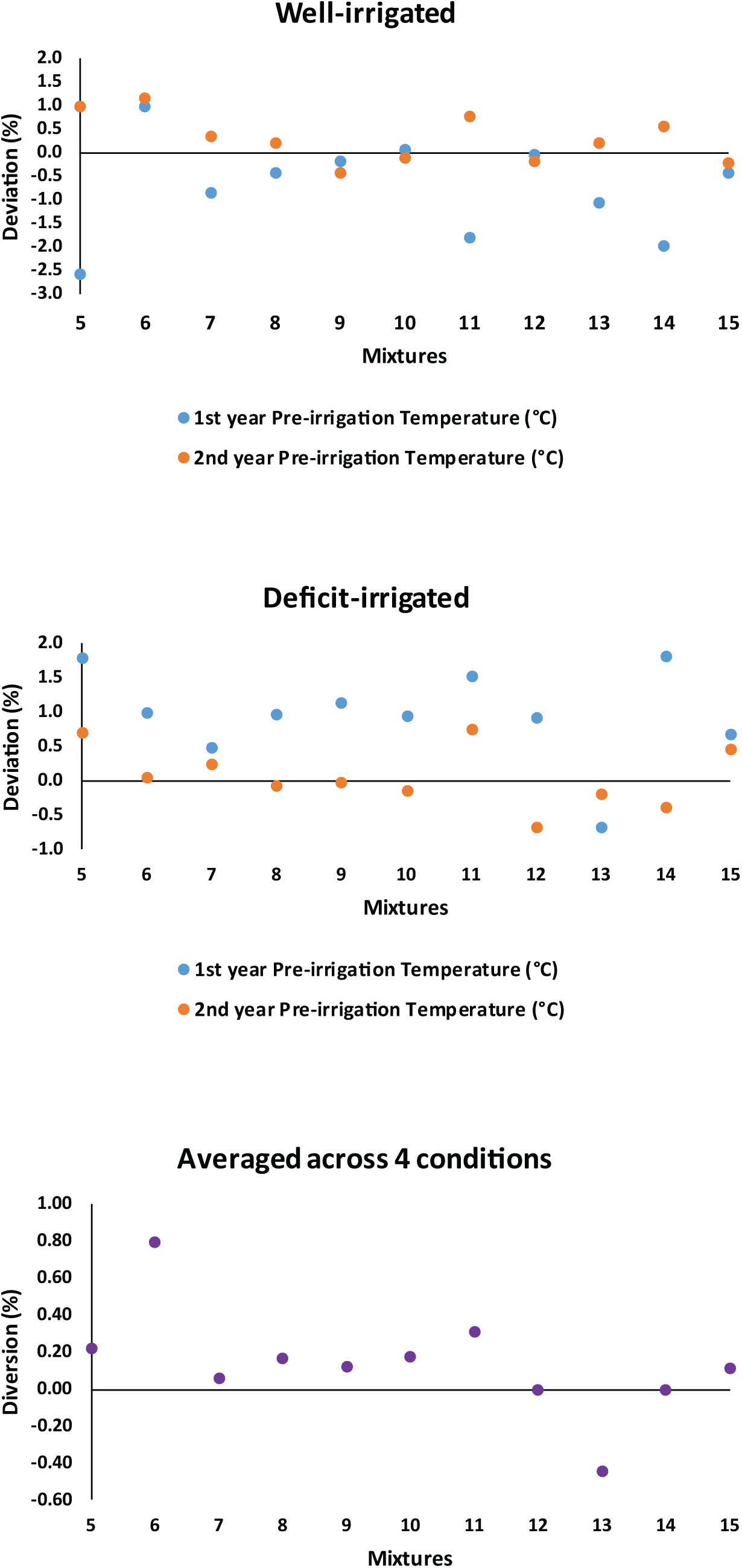
Deviation percentage of pre-irrigation canopy temperature of cultivar mixtures from mean of corresponding monocultures. Treatments 5 to 16 represent the mixtures of 12 (mixture of 1st and 2nd cultivars), 13, 14, 23, 24, 34, 123, 124, 134, 234, and 1234, respectively.

**Figure 11.**
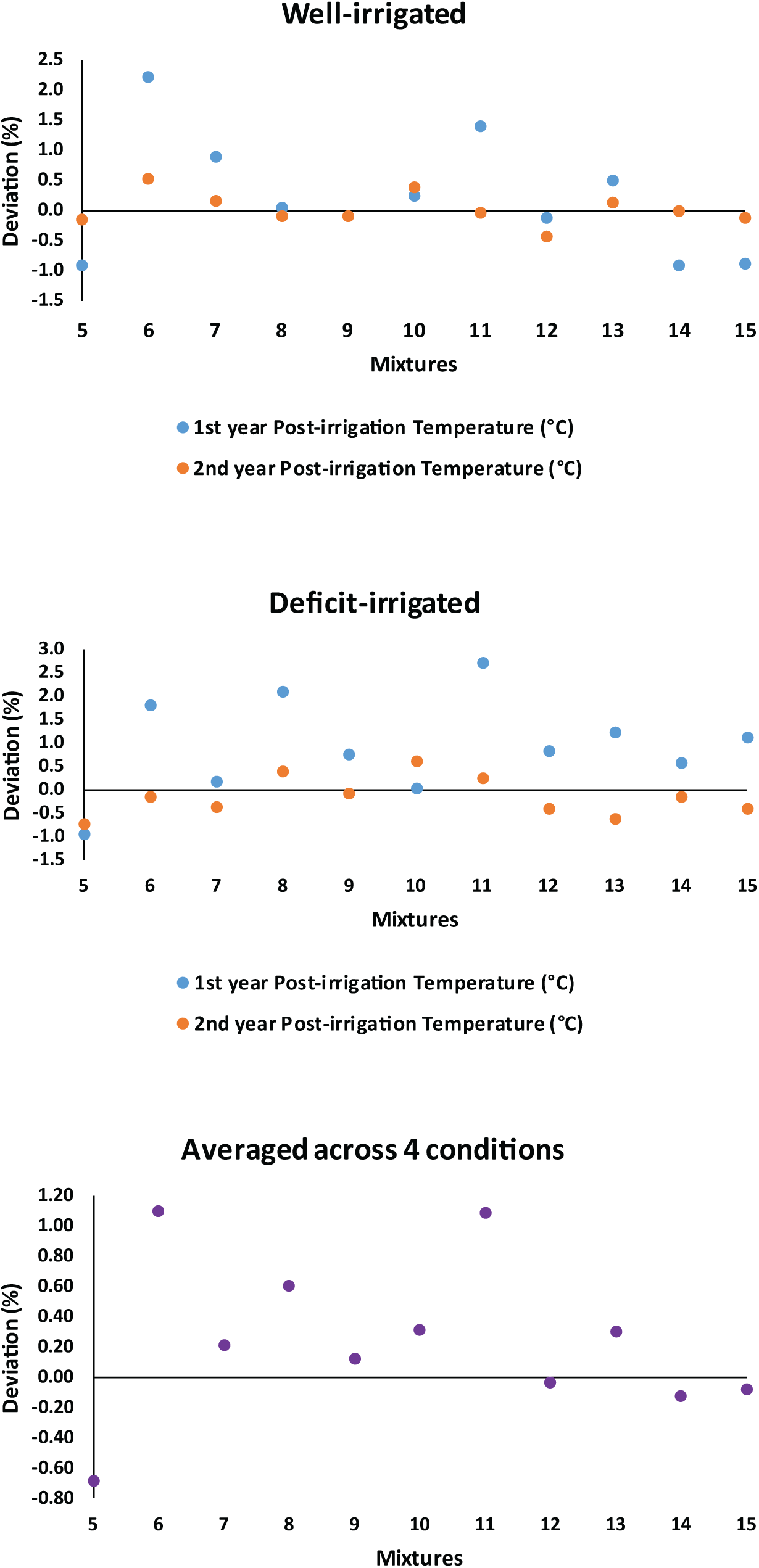
Deviation percentage of post-irrigation canopy temperature of cultivar mixtures from mean of corresponding monocultures. Treatments 5 to 16 represent the mixtures of 12 (mixture of 1st and 2nd cultivars), 13, 14, 23, 24, 34, 123, 124, 134, 234, and 1234, respectively.

Averaged across the 4 conditions (Table 11), mixture 13 had the highest pre-irrigation, and mean canopy temperatures, and also was among the high-temperature treatments based on the post-irrigation values. At the other hand, mixtures 134 and 234 along with monocultures 3 and 4 generally had the lowest triplets of canopy temperature.

**Table 11.**
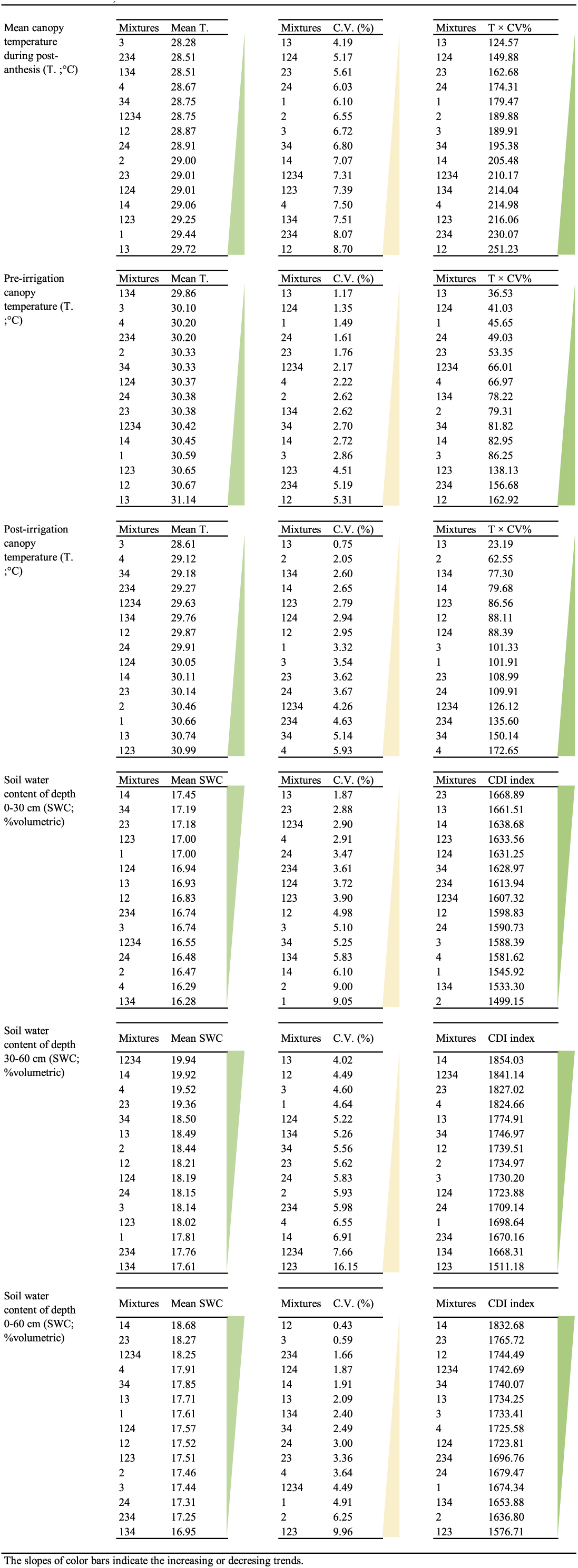
Ranking of mixtures based on mean values, coefficient of variation (C.V.) and Consistent Desirable Index (CDI) of measured characteristics, calculated across 4 environmental conditions.

Orthogonal analysis showed that there was no significant difference between monocultures and mixtures in canopy temperature, including pre- or post-irrigation, or mean temperatures (comparison type I, Table 3). The only significant result of orthogonal analyses of pre-irrigation canopy temperature was found under well-irrigation condition in the 2^nd^ year, where the average value of treatments included the 1^st^ cultivar was higher than those excluded it (Table S1). Also, similar trend was observed for the post-irrigation canopy temperature. Moreover, in the same condition (i.e. well-irrigation in the second year), inclusion of the 3^rd^ cultivar decreased the post-irrigation canopy temperature. Averaged across all conditions, especially under well-irrigation in the 1^st^ year, presence of the latest ripening cultivar (4^th^ cultivar) in the canopy also reduced post-anthesis canopy temperature, which in addition to the previously described results may show the cooler canopies of late ripening cultivars after irrigation, probably due to having a greener stand late in the season.

### 3.6. Soil water content

As mentioned before, since it was only possible to carrying out soil sampling for two out of the three replicates (i.e. from 60 experimental plots), ANOVA and orthogonal analyses were not applicable for soil water content; however, some minor assessments were carried out. ADM analysis (Fig. 12) showed that the deviation of SWC_0-30_ (soil water content in the depth of 0-30 cm) of mixtures from their corresponding monocultures ranged between −0.91 to +2.07 and between −1.14 to +2.33 volumetric percentage under well- and deficit-irrigation, respectively. For SWC_30-60_, these ranges were −2.84% to 2.37% and −1.46% to 1.7%, respectively (Fig. 13). It seems that soil water uptake in the 0-30 cm depth in mixtures was relatively lower than their corresponding monocultures, particularly under deficit-irrigation of the first year (also see average values of 4 conditions, Fig. 12). However, such overall condition was not observed in the case of SWC_30-60_. In both depth, and under all conditions, mixture 14 had comparatively a high level of SWC; whereas mixture 134 always had least values, which implies a greater water uptake in this mixture. Eventually, based on the averaged SWC (SWC_0-60_), mixtures 14, 23, and 1234 had positive and mixtures 24, 134, and 234 had negative deviations compared with their corresponding monocultures; of course in the mentioned range i.e. less than 1% volumetric water content (Fig. 14). Overall across all conditions (Table 11), SWC values of both depths in mixtures placed at the first ranks, either based on the water content percentage or related CDIs.

**Figure 12.**
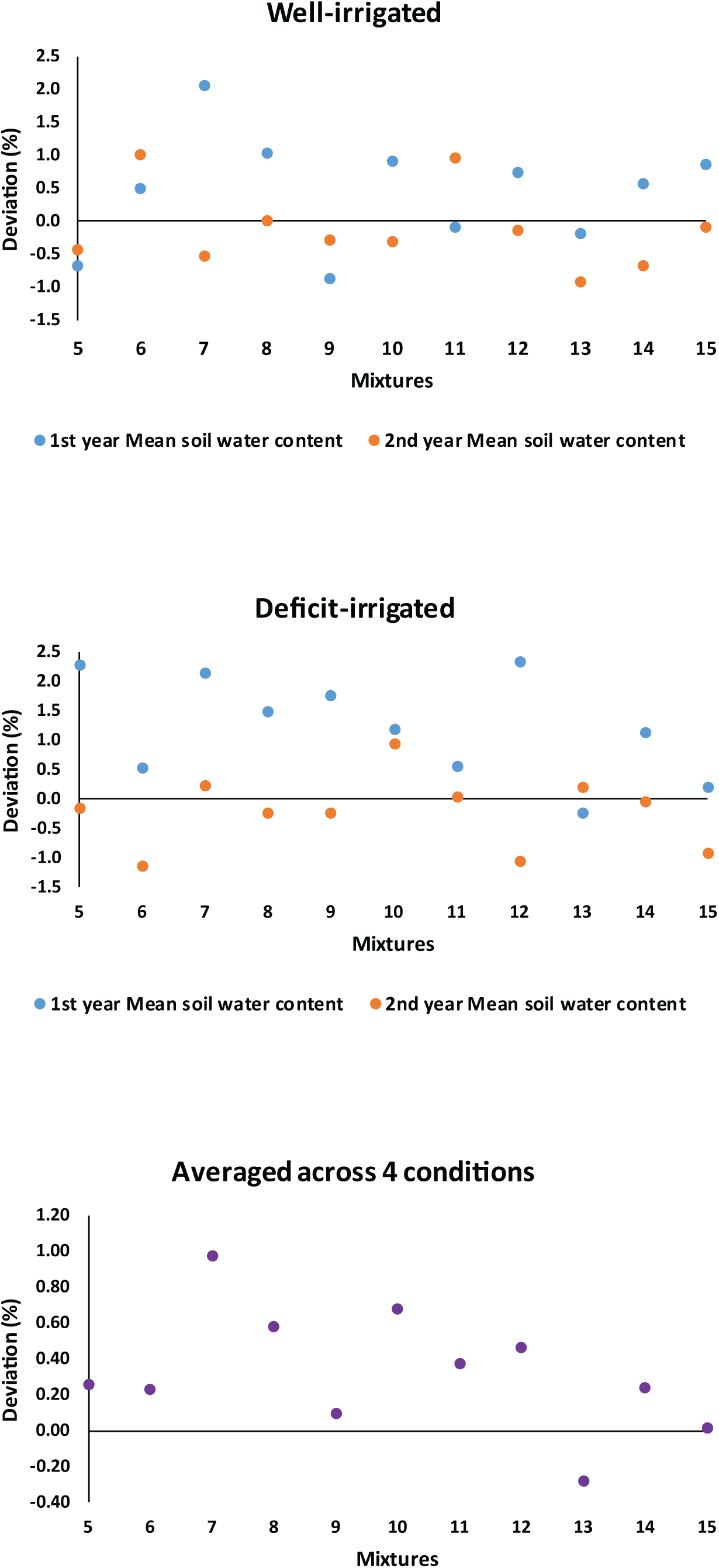
Deviation percentage of soil water content in the 0-30 cm depth of cultivar mixtures from mean of corresponding monocultures. Treatments 5 to 16 represent the mixtures of 12 (mixture of 1st and 2nd cultivars), 13, 14, 23, 24, 34, 123, 124, 134, 234, and 1234, respectively.

**Figure 13.**
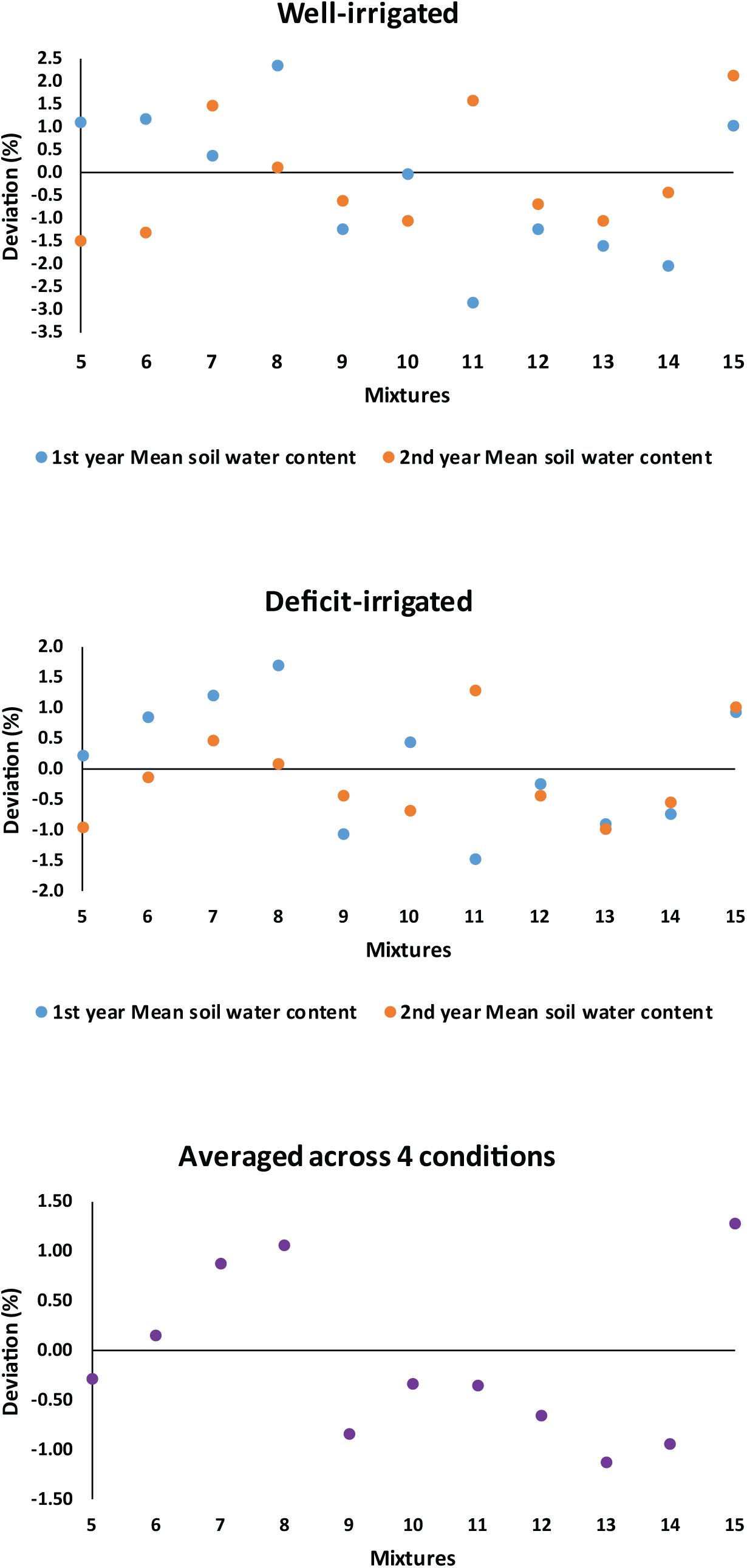
Deviation percentage of soil water content in the 30-60 cm depth of cultivar mixtures from mean of corresponding monocultures. Treatments 5 to 16 represent the mixtures of 12 (mixture of 1st and 2nd cultivars), 13, 14, 23, 24, 34, 123, 124, 134, 234, and 1234, respectively.

**Figure 14.**
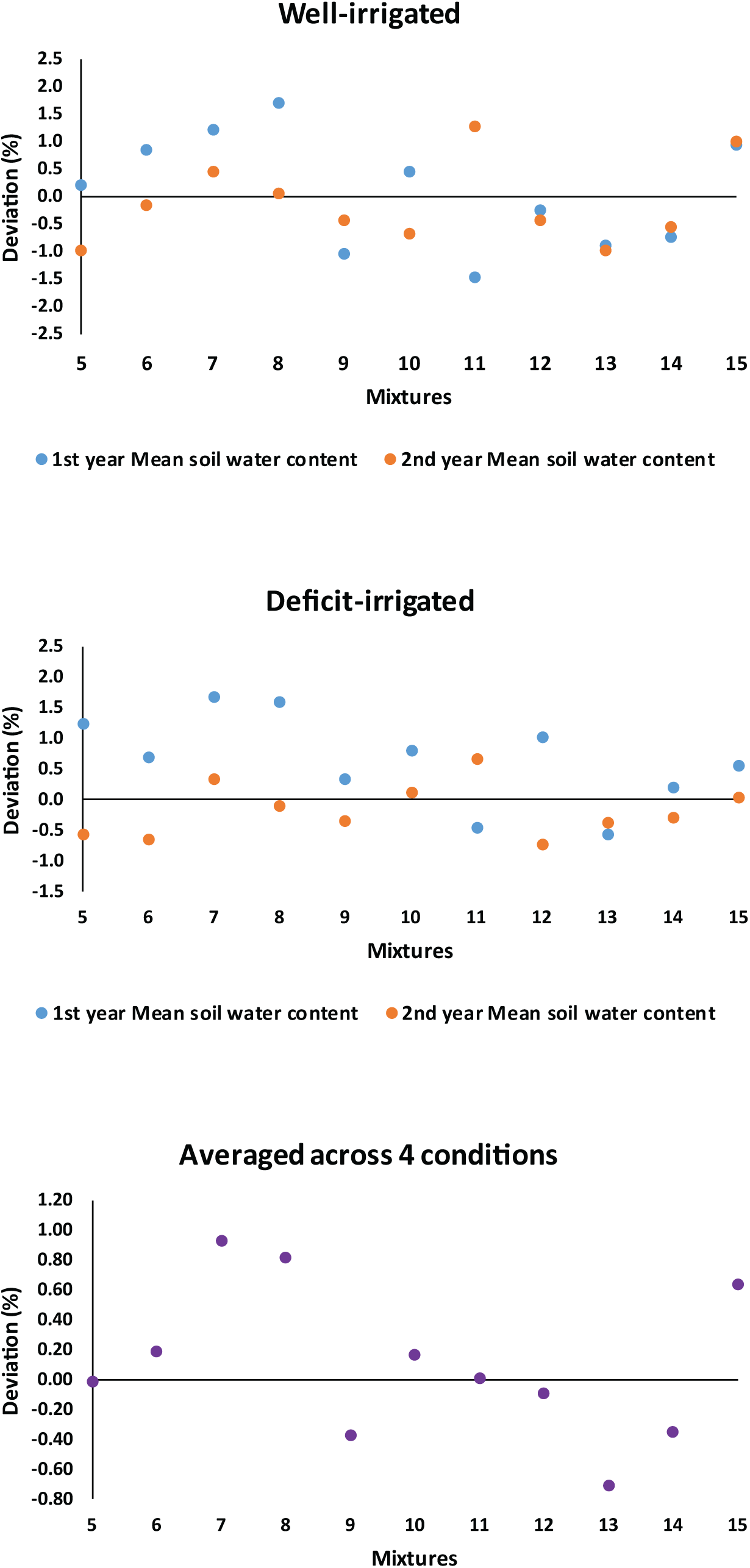
Deviation percentage of soil water content in the 0-60 cm depth of cultivar mixtures from mean of corresponding monocultures. Treatments 5 to 16 represent the mixtures of 12 (mixture of 1st and 2nd cultivars), 13, 14, 23, 24, 34, 123, 124, 134, 234, and 1234, respectively.

Although considering the maximum depth of wheat root system, it is necessary to measure soil water content at greater depths (e.g. see Guo *et al*., 2016), it is known that at least under non-stress conditions, the main portion of root distribution and soil water extraction by wheat occur in the top 30 cm of soil (Sharma & Chaudhary, 1983; Asseng *et al*., 1998).

## 4. Discussion

The present study investigated the consequence of enhancing heterogeneity in occurrence of late season phenological events within conventional wheat cropping system, through a collection of ripening patterns provided by cultivar mixtures. Monocultures of 4 early- to middle ripening cultivars created the expected gradient of ripening (Fig. 2), which shows that the cultivars were selected appropriately. Mixing such cultivars produced a diverse pattern of ripening. Ripening dates were in the range of ripening of the earliest- and the latest-ripening monocultures. However, as one of the most important outcomes of the current experiment, it was revealed that post-anthesis drought stress considerably waived the schemed diversity among the treatments. Although accelerated crop senescence under water stress largely due to enhanced remobilization is a well-known phenomenon (e.g. see Yang *et al*., 2001; Munné-Bosch *et al*., 2004; Yang & Zhang, 2005; Bazargani *et al*., 2011), it seems that the effect of stressful conditions on temporal diversity has rarely been investigated in agroecosystems. However, the negative influence of water stress on temporal heterogeneity described here may be among the first evidences in cropping systems, and be in association with results of ecological studies (conducted at other scales and ecosystems) which have evaluated the mutual relationship between various kinds of stresses and biological diversity in ecosystems (e.g. see Steudel *et al*., 2012; Baert *et al*., 2016; Smeti *et al*., 2019).

In general, the increased similarities in the ripening under stressful conditions were also observed for grain yield and other measured traits; the evidence that indicates the heterogeneity of measured crop properties and mechanisms contribute to grain yield follow the pattern of accelerated ripening. Indeed, both the expected diversity of mixtures in ripening pattern and grain yield was negatively affected by deficit-irrigation, among which the first one was the schemed mechanism for sustaining the latter. As indicated by ADM analyses and various orthogonal comparisons, the hypothesized idea of inducing heterogeneous phenology in late season (Fig. S1) did not lead to an overall consistent improve in evaluated traits. Even under well-irrigated condition, where the designed diversity was relatively occurred, no considerable advantage was observed in the main traits (i.e. grain yield and water productivity) of mixtures compared with monocultures. Therefore, under conditions of the present study, monoculture of the high yielding cultivars with the highest degrees of homogeneity in the late season phenology seems to be remained recommendable. Fletcher *et al*. (2019) reported an average over-yielding of 6.4% for mixtures of wheat cultivars with contrasting phenology, only in one out of the three experiments conducted at different locations. They believed that the value of cultivar mixtures with differing phenology depends on location, climate and sowing date. Similarly, according to the results of the present study, we also suggest that the effect of season may highly affect the comparative performance of mixtures vs. monocultures. Obviously, this is the case for many results of the study including values of susceptibility index of mixtures which showed a completely different trend i.e. +9% vs. −15% of deviation from monoculture in the 1^st^ and 2^nd^ seasons, respectively.

It was relatively a frequent observation in the present study that the only significant difference among various comparisons, was presence or absence of a specific cultivar in the stand, rather than the properties associated with mixing and the structure of canopy e.g. number of component, degree of diversity in phenological events, etc. For instance, consider the significant differences in GY, yield components, and water productivity, made by inclusion or exclusion of the 2^nd^ cultivar. Moreover, only presence or absence of individual cultivars made the significant differences in pre-and post-irrigation canopy temperatures, and mixing properties or complexity of the canopy structure did not lead to considerable variations.

In contrast, as a promising evidence for benefits of mixing cultivars with contrasting phenology, it was observed that under both irrigation conditions in the first year, the 4-component mixture had a significantly lower water requirement; however, this potentially considerable advantage did not repeat in the second year. Besides, only under the well-irrigation condition of the 2^nd^ year, water productivity was significantly reduced in the mixtures with maximum degree of extension in ripening (i.e. those simultaneously included the 1^st^ and 4^th^ cultivars). Among the other rare notable effects of mixtures, was the results of soil water content, which was higher in the first 30 cm of soil layer of the mixtures, compared with monocultures. Interestingly, maximum and minimum SWCs of both depth was recorded in two mixtures, under every conditions.

As the second goal of the study, individual mixtures were screened for potential advantages in grain yield, water productivity, and other measured traits. The results showed that in all conditions, no mixture had grain yields out of the range determined by monocultures, as the lower and higher extremes. However, always (under every 4 conditions) there were mixtures with statistically similar grain yields and some other properties with the superior monoculture. This pattern seems to be a well-known behavior of cultivar mixtures, as frequently have been reported in various studies (e.g. see Haghshenas *et al*., 2013; Fang *et al*., 2014; and Fletcher *et al*., 2019). According to various types of analyses carried out, in many cases there were disadvantageous mixture those led to considerably reduction in the values of the investigated traits (sometimes number of such undesirable blends was higher than the beneficial ones; e.g. see the results of ADM analyses on GY). Indeed, if various aspects of mixing the given cultivars are not understood enough, it may contrarily result in unfavorable outcomes. Therefore, it seems that the general recommendation of utilizing mixtures for enhancing the agronomic or agroecological aspects without running exclusive local experiments (and relying only on the theoretical expectation of increased ecological services in cultivar mixtures) is highly risky.

While the focus of the present study was on describing the consequences of heterogeneous patterns of phenology during late season, the inevitable role of inter-component interaction in earlier stages should not be neglected. Particularly concerning the grain yield, the most important component i.e. number of grains m^−2^ is determined at most few days after anthesis (e.g. see the relative contribution of pre-anthesis days to formation of the grain yield components, reviewed by Fischer, 2011; and also see Slafer *et al*., 2014). Moreover, although determination of mean grain weight is mainly associated with the grain filling period, its potential may be determined by events up to about a week before or after anthesis; Fischer, 2011). Thus, pre-anthesis events should also be considered enough in the performance interpretation of cultivar mixture, particularly those affect grain number m^−2^.

Although the few cases of beneficial effects of mixtures on various traits were not consistent across the seasons or irrigation conditions, they may be a clue in future studies for designing desirable cultivar mixtures (e.g. by selecting or breeding appropriate components). As a conclusion for this part (i.e. screening the potential superior mixtures for readily recommending to growers) no option was found with a significant and consistent advantage compared with monocultures; however, under specific production situations, e.g. when sufficient amounts of seeds of superior cultivar are not available in an area and/or in a season, higher-ranking mixtures which have not significant difference with monocultures may be considered to compensate for the shortage.

Methodologically, evaluation of the complex canopies of cultivar mixtures may require novel and more efficient approaches, as for instance, quantified tracking of ripening in the present study required a transition from the conventional qualitative phenology to a quantitative image-based framework. Moreover, it seems that considering the spatial diversity in the canopies of mixtures, a generally greater area should be sampled to avoid the potential errors which may be raised from taking the samples from a homogenous patch within the stand. Another example of challenges which might be problematic in the current study was making a reasonable and scientific decision about applying (or taking into account) the last irrigation in the season, which might be extremely lead to miscalculations, due to the contrasting patches of still-green to dried areas in mixtures. Therefore, despite the simple and unexpansive practices of mixing and growing cultivars (which almost carried out similar to monocultures), the limitations in the experimental techniques and/or computations should be known, prior to conducting the study.

## 5. Conclusion

In the present study, the option of utilizing the cultivar mixtures with contrasting ripening patterns with the aim of improving grain yield, yield stability, water productivity, and some other agronomic properties was tested under different irrigation regimes. The results indicated that under the stressful condition of post-anthesis deficit-irrigation, the expected heterogeneities in the ripening pattern of mixtures were declined. Consequently, dissimilarities in grain yields as well as various agronomic characters of mixture treatments were also lessened. This may be an evidence for the negative effect of water shortage stress on heterogeneity within agroecosystems. Also, it was observed that the effect of year on most of the measured traits was significant, even in the diversified canopies of mixtures, which were expected to have higher degrees of stability. Although cultivar mixtures showed some casual advantages for various characters, such beneficial effects were not consistent across all conditions; as for instance, values of susceptibility index calculated for the effect of post-anthesis deficit-irrigation on grain yield were +9% vs. −15% in the 1^st^ and 2^nd^ seasons, respectively. No cultivar mixture produced grain yield higher than the maximum monoculture. It is also notable that despite the general expectation of enhancing ecological services and probably agronomic properties, in many cases even disadvantageous blends were found which led to considerable reductions in the grain yield and some measured traits. Therefore, it seems that unless the performances, and preferably the involved mechanisms, of cultivar mixtures are not fully understood, using blends as an alternative for conventional high-input wheat cropping systems may lead to adverse results.

---

**Supplementary Figure S1.**
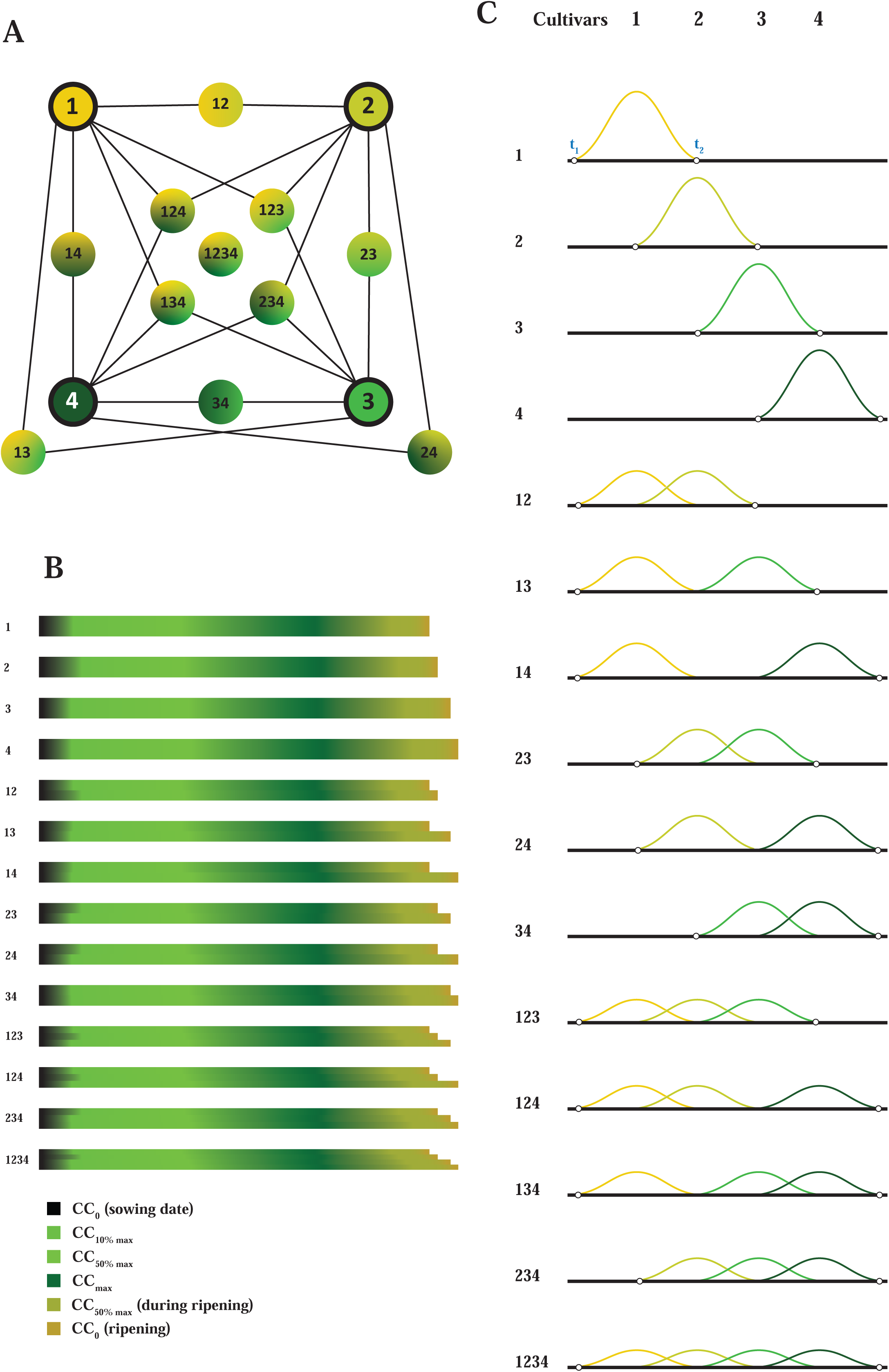
Schematic illustration of the hypothesis of inducing temporal heterogeneity in wheat canopy using cultivar mixtures with diverse ripening patterns. **A**. Mixture treatments included 4 early- to middle ripening cultivars and all of possible 2-, 3-, and 4-component mixtures. The 1^st^, 2^nd^, 3^rd^, and 4^th^ cultivars are the early- to middle-ripening cultivars, which is also shown by yellow to green colors. **B**. The schemed temporal heterogeneity in phenology of monocultures and mixtures, according to the image-based method (described in chapter I). The color gradients of bars show the phenological transitions in the canopy, and also the thicknesses of bars represent the frequency of a given phenological pattern in the canopy. It is notable that the patterns of the first 4 bars (monocultures) are taken from the results of image-based evaluation of phenology of the 4 monocultures in the 2^nd^ year, and the patterns of the remained bars are the theoretical expectation of mixing the cultivars with different phenology; however, the actual image-based analyses carried out in the present study, provide the overall pattern of ripening in each mixture as a single bar, instead of several adjacent bars for the mixture components (as illustrated theoretically in this figure). **C**. The hypothesized comparative occurrence of a given phenological event (e.g. anthesis) in the four monocultures and mixture canopies. The phenological event initiates at *t*_1_ and terminates at *t*_2_. The under-curve-areas show the total number of plants entered the intendent phase (although the shape of the curve may be another, in the lack of information in the literature, the schematic bell-shaped curves are drawn). Consider the comparative lengths of duration and fluctuations of the phenological phase. Each digit in the name of mixture treatment, shows the presence of the respected cultivar in the mixture; e.g. mixture 124 is consisted of the 1^st^, 2^nd^, and 4^th^ cultivars.

**Supplementary Figure S2.**
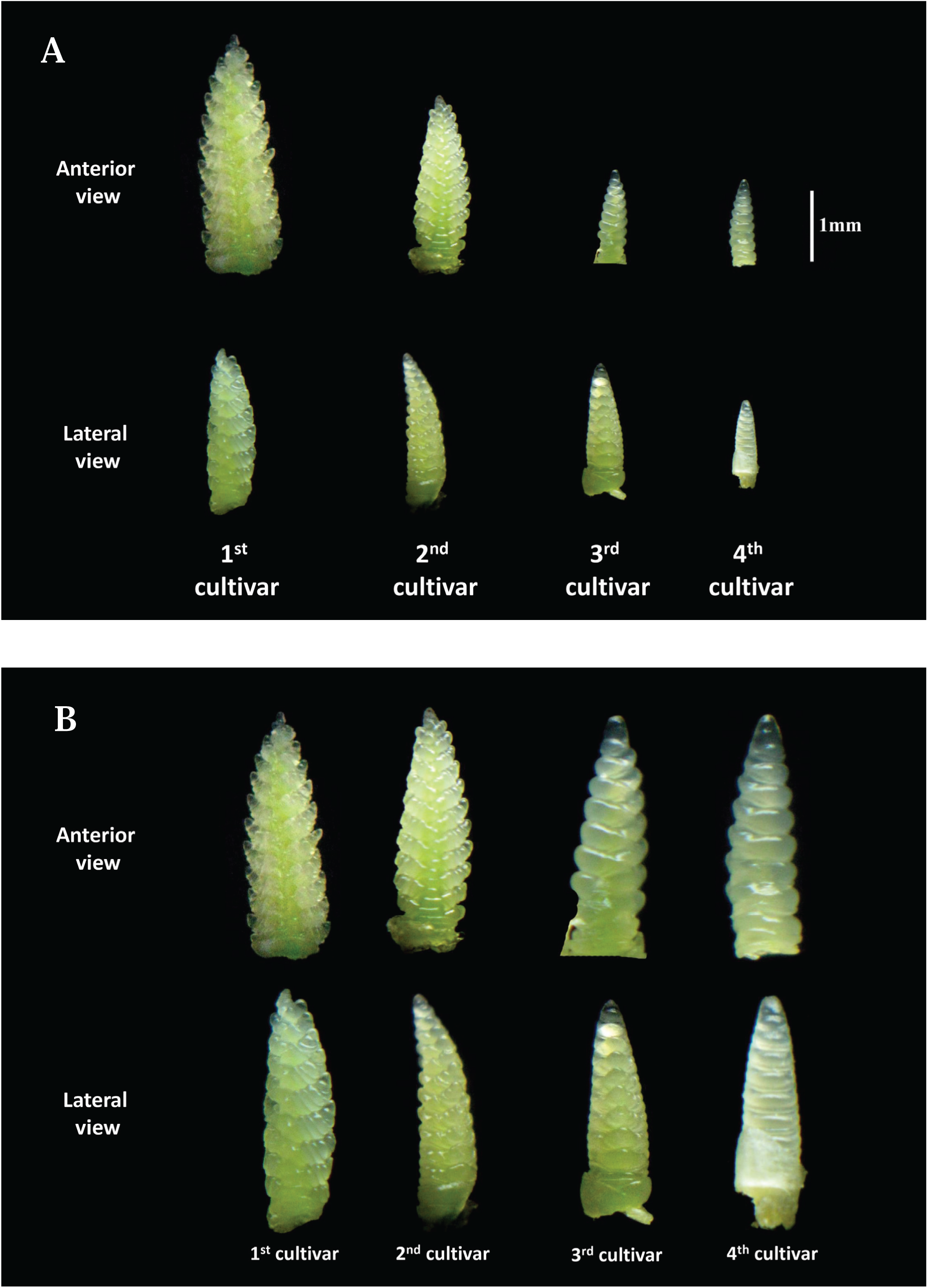
Comparative spike develompent of 4 early- to middle ripening wheat cultivars, 125 days after sowing in the second season. **A**. Spikes are shown in real relative size; **B**. Spikes are shown in normalized equal size; consider the developmental phase irrespective the size.

**Supplementary Figure S3.**
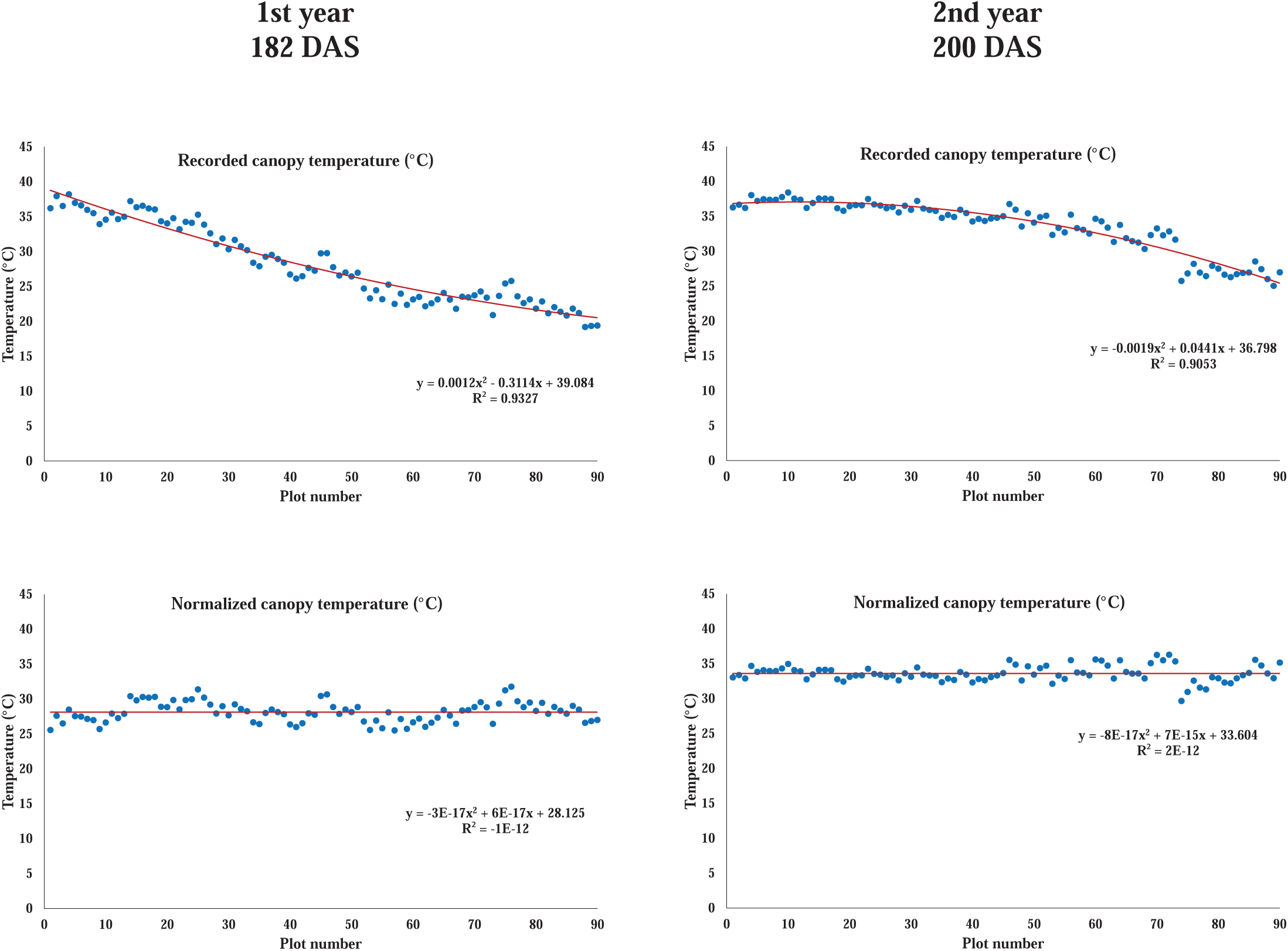
Normalization of canopy temperatures for two sampling dates in the 1^st^ and 2^nd^ years. Recorded temperatures in all other samling dates were also normalized in a same way. DAS: days after sowing.

**Supplementary Figure S4.**
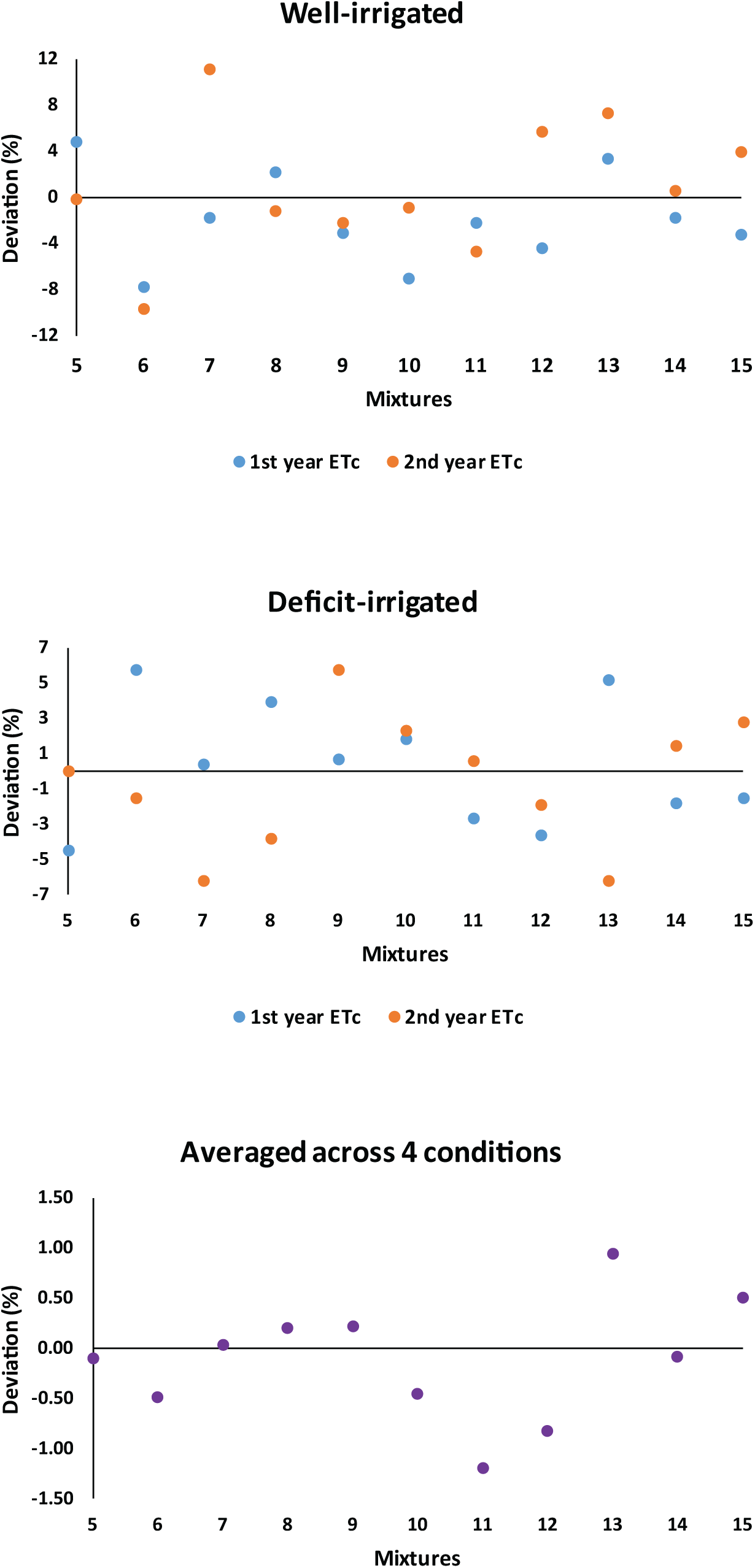
Deviation percentage of evapotranspiration of cultivar mixtures from mean of corresponding monocultures. Treatments 5 to 16 represent the mixtures of 12 (mixture of 1st and 2nd cultivars), 13, 14, 23, 24, 34, 123, 124, 134, 234, and 1234, respectively.

**Supplementary Figure S5.**
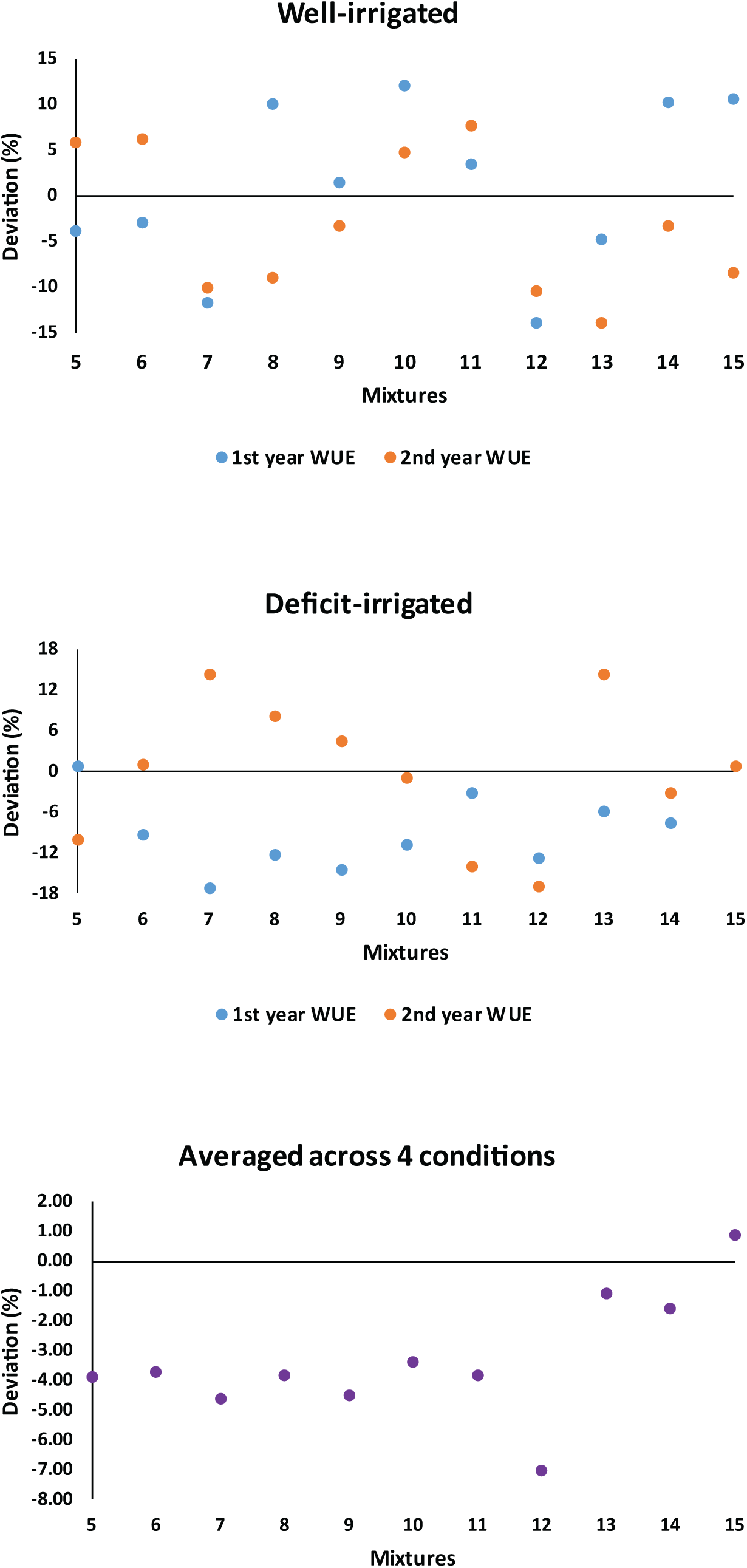
Deviation percentage of water use efficiency of cultivar mixtures from mean of corresponding monocultures. Treatments 5 to 16 represent the mixtures of 12 (mixture of 1st and 2nd cultivars), 13, 14, 23, 24, 34, 123, 124, 134, 234, and 1234, respectively.

**Table S1.**
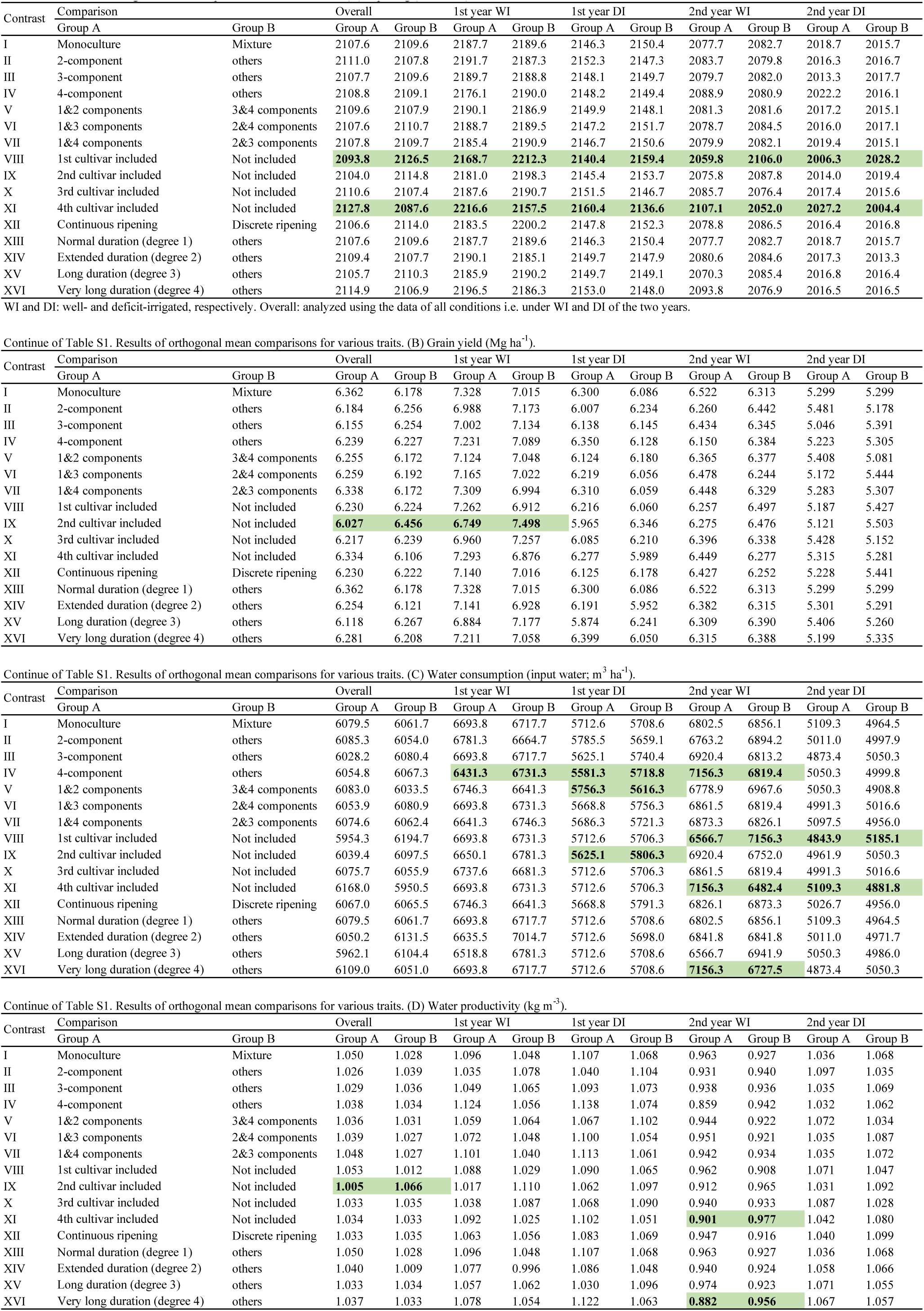
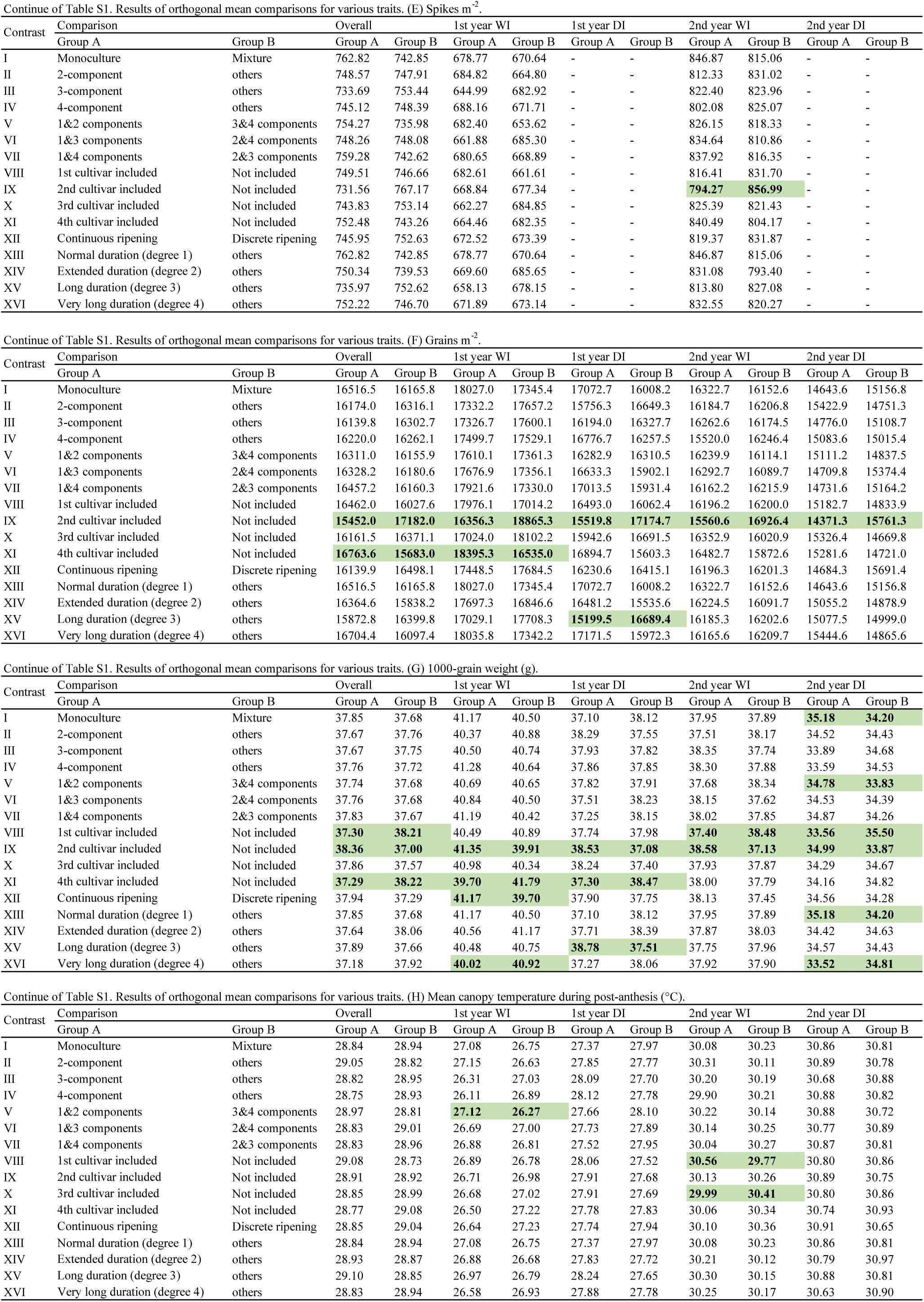
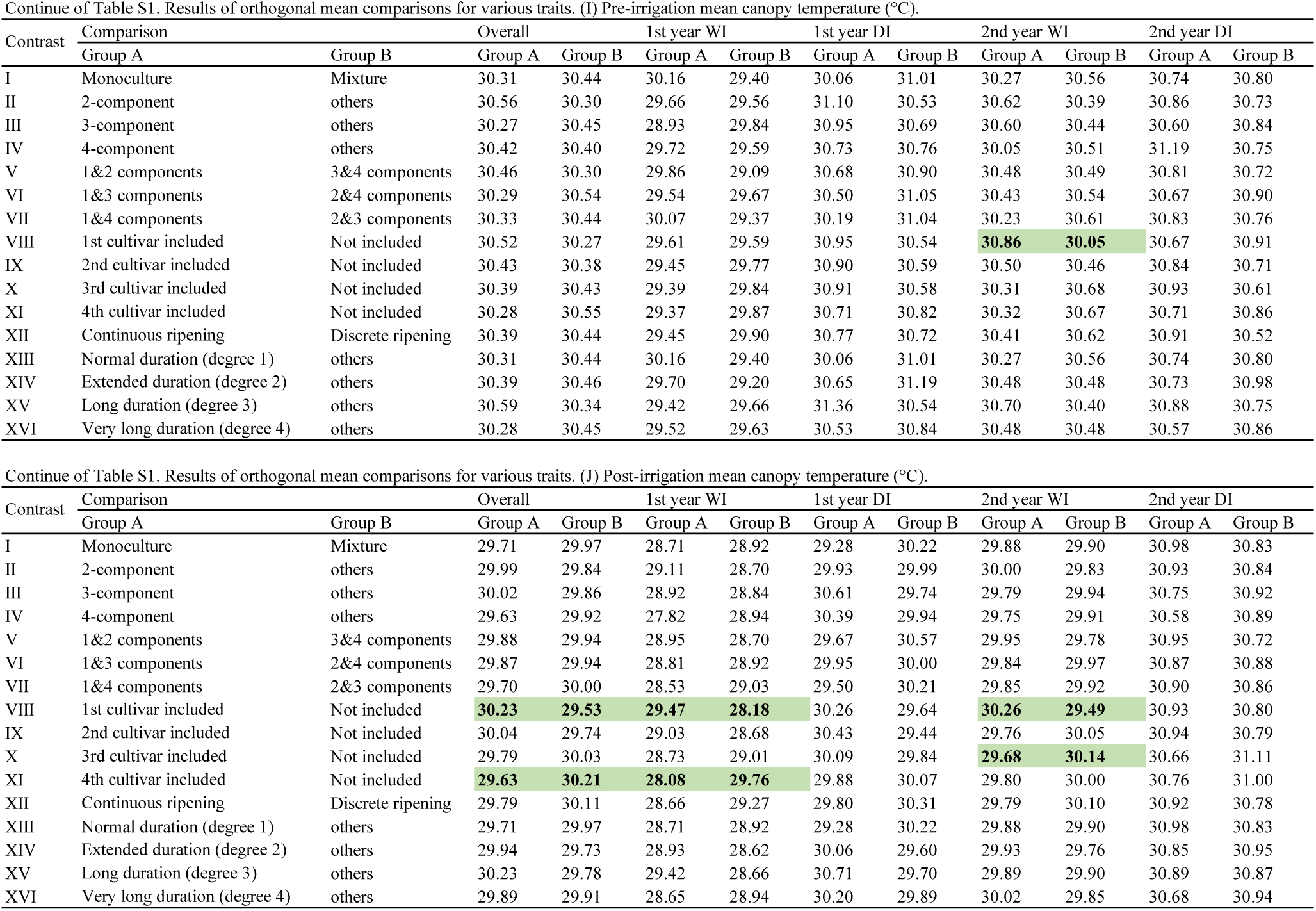
Results of orthogonal mean comparisons for various traits. (A) Ripening (ATT, °Cd).

| Contrast | Comparison |  | Overall |  | 1st year WI |  | 1st year DI |  | 2nd year WI |  | 2nd year DI |  |
| --- | --- | --- | --- | --- | --- | --- | --- | --- | --- | --- | --- | --- |
|  | Group A | Group B | Group A | Group B | Group A | Group B | Group A | Group B | Group A | Group B | Group A | Group B |
| I | Monoculture | Mixture | 2107.6 | 2109.6 | 2187.7 | 2189.6 | 2146.3 | 2150.4 | 2077.7 | 2082.7 | 2018.7 | 2015.7 |
| II | 2-component | others | 2111.0 | 2107.8 | 2191.7 | 2187.3 | 2152.3 | 2147.3 | 2083.7 | 2079.8 | 2016.3 | 2016.7 |
| III | 3-component | others | 2107.7 | 2109.6 | 2189.7 | 2188.8 | 2148.1 | 2149.7 | 2079.7 | 2082.0 | 2013.3 | 2017.7 |
| IV | 4-component | others | 2108.8 | 2109.1 | 2176.1 | 2190.0 | 2148.2 | 2149.4 | 2088.9 | 2080.9 | 2022.2 | 2016.1 |
| V | 1&2 components | 3&4 components | 2109.6 | 2107.9 | 2190.1 | 2186.9 | 2149.9 | 2148.1 | 2081.3 | 2081.6 | 2017.2 | 2015.1 |
| VI | 1&3 components | 2&4 components | 2107.6 | 2110.7 | 2188.7 | 2189.5 | 2147.2 | 2151.7 | 2078.7 | 2084.5 | 2016.0 | 2017.1 |
| VII | 1&4 components | 2&3 components | 2107.8 | 2109.7 | 2185.4 | 2190.9 | 2146.7 | 2150.6 | 2079.9 | 2082.1 | 2019.4 | 2015.1 |
| VIII | 1st cultivar included | Not included | **2093.8** | **2126.5** | **2168.7** | **2212.3** | **2140.4** | **2159.4** | **2059.8** | **2106.0** | **2006.3** | **2028.2** |
| IX | 2nd cultivar included | Not included | 2104.0 | 2114.8 | 2181.0 | 2198.3 | 2145.4 | 2153.7 | 2075.8 | 2087.8 | 2014.0 | 2019.4 |
| X | 3rd cultivar included | Not included | 2110.6 | 2107.4 | 2187.6 | 2190.7 | 2151.5 | 2146.7 | 2085.7 | 2076.4 | 2017.4 | 2015.6 |
| XI | 4th cultivar included | Not included | **2127.8** | **2087.6** | **2216.6** | **2157.5** | **2160.4** | **2136.6** | **2107.1** | **2052.0** | **2027.2** | **2004.4** |
| XII | Continuous ripening | Discrete ripening | 2106.6 | 2114.0 | 2183.5 | 2200.2 | 2147.8 | 2152.3 | 2078.8 | 2086.5 | 2016.4 | 2016.8 |
| XIII | Normal duration (degree 1) | others | 2107.6 | 2109.6 | 2187.7 | 2189.6 | 2146.3 | 2150.4 | 2077.7 | 2082.7 | 2018.7 | 2015.7 |
| XIV | Extended duration (degree 2) | others | 2109.4 | 2107.7 | 2190.1 | 2185.1 | 2149.7 | 2147.9 | 2080.6 | 2084.6 | 2017.3 | 2013.3 |
| XV | Long duration (degree 3) | others | 2105.7 | 2110.3 | 2185.9 | 2190.2 | 2149.7 | 2149.1 | 2070.3 | 2085.4 | 2016.8 | 2016.4 |
| XVI | Very long duration (degree 4) | others | 2114.9 | 2106.9 | 2196.5 | 2186.3 | 2153.0 | 2148.0 | 2093.8 | 2076.9 | 2016.5 | 2016.5 |

| Contrast | Comparison |  | Overall |  | 1st year WI |  | 1st year DI |  | 2nd year WI |  | 2nd year DI |  |
| --- | --- | --- | --- | --- | --- | --- | --- | --- | --- | --- | --- | --- |
|  | Group A | Group B | Group A | Group B | Group A | Group B | Group A | Group B | Group A | Group B | Group A | Group B |
| I | Monoculture | Mixture | 6.362 | 6.178 | 7.328 | 7.015 | 6.300 | 6.086 | 6.522 | 6.313 | 5.299 | 5.299 |
| II | 2-component | others | 6.184 | 6.256 | 6.988 | 7.173 | 6.007 | 6.234 | 6.260 | 6.442 | 5.481 | 5.178 |
| III | 3-component | others | 6.155 | 6.254 | 7.002 | 7.134 | 6.138 | 6.145 | 6.434 | 6.345 | 5.046 | 5.391 |
| IV | 4-component | others | 6.239 | 6.227 | 7.231 | 7.089 | 6.350 | 6.128 | 6.150 | 6.384 | 5.223 | 5.305 |
| V | 1&2 components | 3&4 components | 6.255 | 6.172 | 7.124 | 7.048 | 6.124 | 6.180 | 6.365 | 6.377 | 5.408 | 5.081 |
| VI | 1&3 components | 2&4 components | 6.259 | 6.192 | 7.165 | 7.022 | 6.219 | 6.056 | 6.478 | 6.244 | 5.172 | 5.444 |
| VII | 1&4 components | 2&3 components | 6.338 | 6.172 | 7.309 | 6.994 | 6.310 | 6.059 | 6.448 | 6.329 | 5.283 | 5.307 |
| VIII | 1st cultivar included | Not included | 6.230 | 6.224 | 7.262 | 6.912 | 6.216 | 6.060 | 6.257 | 6.497 | 5.187 | 5.427 |
| IX | 2nd cultivar included | Not included | **6.027** | **6.456** | **6.749** | **7.498** | 5.965 | 6.346 | 6.275 | 6.476 | 5.121 | 5.503 |
| X | 3rd cultivar included | Not included | 6.217 | 6.239 | 6.960 | 7.257 | 6.085 | 6.210 | 6.396 | 6.338 | 5.428 | 5.152 |
| XI | 4th cultivar included | Not included | 6.334 | 6.106 | 7.293 | 6.876 | 6.277 | 5.989 | 6.449 | 6.277 | 5.315 | 5.281 |
| XII | Continuous ripening | Discrete ripening | 6.230 | 6.222 | 7.140 | 7.016 | 6.125 | 6.178 | 6.427 | 6.252 | 5.228 | 5.441 |
| XIII | Normal duration (degree 1) | others | 6.362 | 6.178 | 7.328 | 7.015 | 6.300 | 6.086 | 6.522 | 6.313 | 5.299 | 5.299 |
| XIV | Extended duration (degree 2) | others | 6.254 | 6.121 | 7.141 | 6.928 | 6.191 | 5.952 | 6.382 | 6.315 | 5.301 | 5.291 |
| XV | Long duration (degree 3) | others | 6.118 | 6.267 | 6.884 | 7.177 | 5.874 | 6.241 | 6.309 | 6.390 | 5.406 | 5.260 |
| XVI | Very long duration (degree 4) | others | 6.281 | 6.208 | 7.211 | 7.058 | 6.399 | 6.050 | 6.315 | 6.388 | 5.199 | 5.335 |

| Contrast | Comparison |  | Overall |  | 1st year WI |  | 1st year DI |  | 2nd year WI |  | 2nd year DI |  |
| --- | --- | --- | --- | --- | --- | --- | --- | --- | --- | --- | --- | --- |
|  | Group A | Group B | Group A | Group B | Group A | Group B | Group A | Group B | Group A | Group B | Group A | Group B |
| I | Monoculture | Mixture | 6079.5 | 6061.7 | 6693.8 | 6717.7 | 5712.6 | 5708.6 | 6802.5 | 6856.1 | 5109.3 | 4964.5 |
| II | 2-component | others | 6085.3 | 6054.0 | 6781.3 | 6664.7 | 5785.5 | 5659.1 | 6763.2 | 6894.2 | 5011.0 | 4997.9 |
| III | 3-component | others | 6028.2 | 6080.4 | 6693.8 | 6717.7 | 5625.1 | 5740.4 | 6920.4 | 6813.2 | 4873.4 | 5050.3 |
| IV | 4-component | others | 6054.8 | 6067.3 | **6431.3** | **6731.3** | **5581.3** | **5718.8** | **7156.3** | **6819.4** | 5050.3 | 4999.8 |
| V | 1&2 components | 3&4 components | 6083.0 | 6033.5 | 6746.3 | 6641.3 | **5756.3** | **5616.3** | 6778.9 | 6967.6 | 5050.3 | 4908.8 |
| VI | 1&3 components | 2&4 components | 6053.9 | 6080.9 | 6693.8 | 6731.3 | 5668.8 | 5756.3 | 6861.5 | 6819.4 | 4991.3 | 5016.6 |
| VII | 1&4 components | 2&3 components | 6074.6 | 6062.4 | 6641.3 | 6746.3 | 5686.3 | 5721.3 | 6873.3 | 6826.1 | 5097.5 | 4956.0 |
| VIII | 1st cultivar included | Not included | 5954.3 | 6194.7 | 6693.8 | 6731.3 | 5712.6 | 5706.3 | **6566.7** | **7156.3** | **4843.9** | **5185.1** |
| IX | 2nd cultivar included | Not included | 6039.4 | 6097.5 | 6650.1 | 6781.3 | **5625.1** | **5806.3** | 6920.4 | 6752.0 | 4961.9 | 5050.3 |
| X | 3rd cultivar included | Not included | 6075.7 | 6055.9 | 6737.6 | 6681.3 | 5712.6 | 5706.3 | 6861.5 | 6819.4 | 4991.3 | 5016.6 |
| XI | 4th cultivar included | Not included | 6168.0 | 5950.5 | 6693.8 | 6731.3 | 5712.6 | 5706.3 | **7156.3** | **6482.4** | **5109.3** | **4881.8** |
| XII | Continuous ripening | Discrete ripening | 6067.0 | 6065.5 | 6746.3 | 6641.3 | 5668.8 | 5791.3 | 6826.1 | 6873.3 | 5026.7 | 4956.0 |
| XIII | Normal duration (degree 1) | others | 6079.5 | 6061.7 | 6693.8 | 6717.7 | 5712.6 | 5708.6 | 6802.5 | 6856.1 | 5109.3 | 4964.5 |
| XIV | Extended duration (degree 2) | others | 6050.2 | 6131.5 | 6635.5 | 7014.7 | 5712.6 | 5698.0 | 6841.8 | 6841.8 | 5011.0 | 4971.7 |
| XV | Long duration (degree 3) | others | 5962.1 | 6104.4 | 6518.8 | 6781.3 | 5712.6 | 5708.6 | 6566.7 | 6941.9 | 5050.3 | 4986.0 |
| XVI | Very long duration (degree 4) | others | 6109.0 | 6051.0 | 6693.8 | 6717.7 | 5712.6 | 5708.6 | **7156.3** | **6727.5** | 4873.4 | 5050.3 |

| Contrast | Comparison |  | Overall |  | 1st year WI |  | 1st year DI |  | 2nd year WI |  | 2nd year DI |  |
| --- | --- | --- | --- | --- | --- | --- | --- | --- | --- | --- | --- | --- |
|  | Group A | Group B | Group A | Group B | Group A | Group B | Group A | Group B | Group A | Group B | Group A | Group B |
| I | Monoculture | Mixture | 1.050 | 1.028 | 1.096 | 1.048 | 1.107 | 1.068 | 0.963 | 0.927 | 1.036 | 1.068 |
| II | 2-component | others | 1.026 | 1.039 | 1.035 | 1.078 | 1.040 | 1.104 | 0.931 | 0.940 | 1.097 | 1.035 |
| III | 3-component | others | 1.029 | 1.036 | 1.049 | 1.065 | 1.093 | 1.073 | 0.938 | 0.936 | 1.035 | 1.069 |
| IV | 4-component | others | 1.038 | 1.034 | 1.124 | 1.056 | 1.138 | 1.074 | 0.859 | 0.942 | 1.032 | 1.062 |
| V | 1&2 components | 3&4 components | 1.036 | 1.031 | 1.059 | 1.064 | 1.067 | 1.102 | 0.944 | 0.922 | 1.072 | 1.034 |
| VI | 1&3 components | 2&4 components | 1.039 | 1.027 | 1.072 | 1.048 | 1.100 | 1.054 | 0.951 | 0.921 | 1.035 | 1.087 |
| VII | 1&4 components | 2&3 components | 1.048 | 1.027 | 1.101 | 1.040 | 1.113 | 1.061 | 0.942 | 0.934 | 1.035 | 1.072 |
| VIII | 1st cultivar included | Not included | 1.053 | 1.012 | 1.088 | 1.029 | 1.090 | 1.065 | 0.962 | 0.908 | 1.071 | 1.047 |
| IX | 2nd cultivar included | Not included | **1.005** | **1.066** | 1.017 | 1.110 | 1.062 | 1.097 | 0.912 | 0.965 | 1.031 | 1.092 |
| X | 3rd cultivar included | Not included | 1.033 | 1.035 | 1.038 | 1.087 | 1.068 | 1.090 | 0.940 | 0.933 | 1.087 | 1.028 |
| XI | 4th cultivar included | Not included | 1.034 | 1.033 | 1.092 | 1.025 | 1.102 | 1.051 | **0.901** | **0.977** | 1.042 | 1.080 |
| XII | Continuous ripening | Discrete ripening | 1.033 | 1.035 | 1.063 | 1.056 | 1.083 | 1.069 | 0.947 | 0.916 | 1.040 | 1.099 |
| XIII | Normal duration (degree 1) | others | 1.050 | 1.028 | 1.096 | 1.048 | 1.107 | 1.068 | 0.963 | 0.927 | 1.036 | 1.068 |
| XIV | Extended duration (degree 2) | others | 1.040 | 1.009 | 1.077 | 0.996 | 1.086 | 1.048 | 0.940 | 0.924 | 1.058 | 1.066 |
| XV | Long duration (degree 3) | others | 1.033 | 1.034 | 1.057 | 1.062 | 1.030 | 1.096 | 0.974 | 0.923 | 1.071 | 1.055 |
| XVI | Very long duration (degree 4) | others | 1.037 | 1.033 | 1.078 | 1.054 | 1.122 | 1.063 | **0.882** | **0.956** | 1.067 | 1.057 |

Continue of Table S1. Results of orthogonal mean comparisons for various traits. (E) Spikes m<sup>-2</sup>.
| Contrast | Comparison |  | Overall |  | 1st year WI |  | 1st year DI |  | 2nd year WI |  | 2nd year DI |  |
| --- | --- | --- | --- | --- | --- | --- | --- | --- | --- | --- | --- | --- |
|  | Group A | Group B | Group A | Group B | Group A | Group B | Group A | Group B | Group A | Group B | Group A | Group B |
| I | Monoculture | Mixture | 762.82 | 742.85 | 678.77 | 670.64 | - | - | 846.87 | 815.06 | - | - |
| II | 2-component | others | 748.57 | 747.91 | 684.82 | 664.80 | - | - | 812.33 | 831.02 | - | - |
| III | 3-component | others | 733.69 | 753.44 | 644.99 | 682.92 | - | - | 822.40 | 823.96 | - | - |
| IV | 4-component | others | 745.12 | 748.39 | 688.16 | 671.71 | - | - | 802.08 | 825.07 | - | - |
| V | 1&2 components | 3&4 components | 754.27 | 735.98 | 682.40 | 653.62 | - | - | 826.15 | 818.33 | - | - |
| VI | 1&3 components | 2&4 components | 748.26 | 748.08 | 661.88 | 685.30 | - | - | 834.64 | 810.86 | - | - |
| VII | 1&4 components | 2&3 components | 759.28 | 742.62 | 680.65 | 668.89 | - | - | 837.92 | 816.35 | - | - |
| VIII | 1st cultivar included | Not included | 749.51 | 746.66 | 682.61 | 661.61 | - | - | 816.41 | 831.70 | - | - |
| IX | 2nd cultivar included | Not included | 731.56 | 767.17 | 668.84 | 677.34 | - | - | **794.27** | **856.99** | - | - |
| X | 3rd cultivar included | Not included | 743.83 | 753.14 | 662.27 | 684.85 | - | - | 825.39 | 821.43 | - | - |
| XI | 4th cultivar included | Not included | 752.48 | 743.26 | 664.46 | 682.35 | - | - | 840.49 | 804.17 | - | - |
| XII | Continuous ripening | Discrete ripening | 745.95 | 752.63 | 672.52 | 673.39 | - | - | 819.37 | 831.87 | - | - |
| XIII | Normal duration (degree 1) | others | 762.82 | 742.85 | 678.77 | 670.64 | - | - | 846.87 | 815.06 | - | - |
| XIV | Extended duration (degree 2) | others | 750.34 | 739.53 | 669.60 | 685.65 | - | - | 831.08 | 793.40 | - | - |
| XV | Long duration (degree 3) | others | 735.97 | 752.62 | 658.13 | 678.15 | - | - | 813.80 | 827.08 | - | - |
| XVI | Very long duration (degree 4) | others | 752.22 | 746.70 | 671.89 | 673.14 | - | - | 832.55 | 820.27 | - | - |

Continue of Table S1. Results of orthogonal mean comparisons for various traits. (F) Grains m<sup>-2</sup>.
| Contrast | Comparison |  | Overall |  | 1st year WI |  | 1st year DI |  | 2nd year WI |  | 2nd year DI |  |
| --- | --- | --- | --- | --- | --- | --- | --- | --- | --- | --- | --- | --- |
|  | Group A | Group B | Group A | Group B | Group A | Group B | Group A | Group B | Group A | Group B | Group A | Group B |
| I | Monoculture | Mixture | 16516.5 | 16165.8 | 18027.0 | 17345.4 | 17072.7 | 16008.2 | 16322.7 | 16152.6 | 14643.6 | 15156.8 |
| II | 2-component | others | 16174.0 | 16316.1 | 17332.2 | 17657.2 | 15756.3 | 16649.3 | 16184.7 | 16206.8 | 15422.9 | 14751.3 |
| III | 3-component | others | 16139.8 | 16302.7 | 17326.7 | 17600.1 | 16194.0 | 16327.7 | 16262.6 | 16174.5 | 14776.0 | 15108.7 |
| IV | 4-component | others | 16220.0 | 16262.1 | 17499.7 | 17529.1 | 16776.7 | 16257.5 | 15520.0 | 16246.4 | 15083.6 | 15015.4 |
| V | 1&2 components | 3&4 components | 16311.0 | 16155.9 | 17610.1 | 17361.3 | 16282.9 | 16310.5 | 16239.9 | 16114.1 | 15111.2 | 14837.5 |
| VI | 1&3 components | 2&4 components | 16328.2 | 16180.6 | 17676.9 | 17356.1 | 16633.3 | 15902.1 | 16292.7 | 16089.7 | 14709.8 | 15374.4 |
| VII | 1&4 components | 2&3 components | 16457.2 | 16160.3 | 17921.6 | 17330.0 | 17013.5 | 15931.4 | 16162.2 | 16215.9 | 14731.6 | 15164.2 |
| VIII | 1st cultivar included | Not included | 16462.0 | 16027.6 | 17976.1 | 17014.2 | 16493.0 | 16062.4 | 16196.2 | 16200.0 | 15182.7 | 14833.9 |
| IX | 2nd cultivar included | Not included | **15452.0** | **17182.0** | **16356.3** | **18865.3** | **15519.8** | **17174.7** | **15560.6** | **16926.4** | **14371.3** | **15761.3** |
| X | 3rd cultivar included | Not included | 16161.5 | 16371.1 | 17024.0 | 18102.2 | 15942.6 | 16691.5 | 16352.9 | 16020.9 | 15326.4 | 14669.8 |
| XI | 4th cultivar included | Not included | **16763.6** | **15683.0** | **18395.3** | **16535.0** | 16894.7 | 15603.3 | 16482.7 | 15872.6 | 15281.6 | 14721.0 |
| XII | Continuous ripening | Discrete ripening | 16139.9 | 16498.1 | 17448.5 | 17684.5 | 16230.6 | 16415.1 | 16196.3 | 16201.3 | 14684.3 | 15691.4 |
| XIII | Normal duration (degree 1) | others | 16516.5 | 16165.8 | 18027.0 | 17345.4 | 17072.7 | 16008.2 | 16322.7 | 16152.6 | 14643.6 | 15156.8 |
| XIV | Extended duration (degree 2) | others | 16364.6 | 15838.2 | 17697.3 | 16846.6 | 16481.2 | 15535.6 | 16224.5 | 16091.7 | 15055.2 | 14878.9 |
| XV | Long duration (degree 3) | others | 15872.8 | 16399.8 | 17029.1 | 17708.3 | **15199.5** | **16689.4** | 16185.3 | 16202.6 | 15077.5 | 14999.0 |
| XVI | Very long duration (degree 4) | others | 16704.4 | 16097.4 | 18035.8 | 17342.2 | 17171.5 | 15972.3 | 16165.6 | 16209.7 | 15444.6 | 14865.6 |

Continue of Table S1. Results of orthogonal mean comparisons for various traits. (G) 1000-grain weight (g).
| Contrast | Comparison |  | Overall |  | 1st year WI |  | 1st year DI |  | 2nd year WI |  | 2nd year DI |  |
| --- | --- | --- | --- | --- | --- | --- | --- | --- | --- | --- | --- | --- |
|  | Group A | Group B | Group A | Group B | Group A | Group B | Group A | Group B | Group A | Group B | Group A | Group B |
| I | Monoculture | Mixture | 37.85 | 37.68 | 41.17 | 40.50 | 37.10 | 38.12 | 37.95 | 37.89 | **35.18** | **34.20** |
| II | 2-component | others | 37.67 | 37.76 | 40.37 | 40.88 | 38.29 | 37.55 | 37.51 | 38.17 | 34.52 | 34.43 |
| III | 3-component | others | 37.67 | 37.75 | 40.50 | 40.74 | 37.93 | 37.82 | 38.35 | 37.74 | 33.89 | 34.68 |
| IV | 4-component | others | 37.76 | 37.72 | 41.28 | 40.64 | 37.86 | 37.85 | 38.30 | 37.88 | 33.59 | 34.53 |
| V | 1&2 components | 3&4 components | 37.74 | 37.68 | 40.69 | 40.65 | 37.82 | 37.91 | 37.68 | 38.34 | **34.78** | **33.83** |
| VI | 1&3 components | 2&4 components | 37.76 | 37.68 | 40.84 | 40.50 | 37.51 | 38.23 | 38.15 | 37.62 | 34.53 | 34.39 |
| VII | 1&4 components | 2&3 components | 37.83 | 37.67 | 41.19 | 40.42 | 37.25 | 38.15 | 38.02 | 37.85 | 34.87 | 34.26 |
| VIII | 1st cultivar included | Not included | **37.30** | **38.21** | 40.49 | 40.89 | 37.74 | 37.98 | **37.40** | **38.48** | **33.56** | **35.50** |
| IX | 2nd cultivar included | Not included | **38.36** | **37.00** | **41.35** | **39.91** | **38.53** | **37.08** | **38.58** | **37.13** | **34.99** | **33.87** |
| X | 3rd cultivar included | Not included | 37.86 | 37.57 | 40.98 | 40.34 | 38.24 | 37.40 | 37.93 | 37.87 | 34.29 | 34.67 |
| XI | 4th cultivar included | Not included | **37.29** | **38.22** | **39.70** | **41.79** | **37.30** | **38.47** | 38.00 | 37.79 | 34.16 | 34.82 |
| XII | Continuous ripening | Discrete ripening | 37.94 | 37.29 | **41.17** | **39.70** | 37.90 | 37.75 | 38.13 | 37.45 | 34.56 | 34.28 |
| XIII | Normal duration (degree 1) | others | 37.85 | 37.68 | 41.17 | 40.50 | 37.10 | 38.12 | 37.95 | 37.89 | **35.18** | **34.20** |
| XIV | Extended duration (degree 2) | others | 37.64 | 38.06 | 40.56 | 41.17 | 37.71 | 38.39 | 37.87 | 38.03 | 34.42 | 34.63 |
| XV | Long duration (degree 3) | others | 37.89 | 37.66 | 40.48 | 40.75 | **38.78** | **37.51** | 37.75 | 37.96 | 34.57 | 34.43 |
| XVI | Very long duration (degree 4) | others | 37.18 | 37.92 | **40.02** | **40.92** | 37.27 | 38.06 | 37.92 | 37.90 | **33.52** | **34.81** |

| Contrast | Comparison |  | Overall |  | 1st year WI |  | 1st year DI |  | 2nd year WI |  | 2nd year DI |  |
| --- | --- | --- | --- | --- | --- | --- | --- | --- | --- | --- | --- | --- |
|  | Group A | Group B | Group A | Group B | Group A | Group B | Group A | Group B | Group A | Group B | Group A | Group B |
| I | Monoculture | Mixture | 28.84 | 28.94 | 27.08 | 26.75 | 27.37 | 27.97 | 30.08 | 30.23 | 30.86 | 30.81 |
| II | 2-component | others | 29.05 | 28.82 | 27.15 | 26.63 | 27.85 | 27.77 | 30.31 | 30.11 | 30.89 | 30.78 |
| III | 3-component | others | 28.82 | 28.95 | 26.31 | 27.03 | 28.09 | 27.70 | 30.20 | 30.19 | 30.68 | 30.88 |
| IV | 4-component | others | 28.75 | 28.93 | 26.11 | 26.89 | 28.12 | 27.78 | 29.90 | 30.21 | 30.88 | 30.82 |
| V | 1&2 components | 3&4 components | 28.97 | 28.81 | **27.12** | **26.27** | 27.66 | 28.10 | 30.22 | 30.14 | 30.88 | 30.72 |
| VI | 1&3 components | 2&4 components | 28.83 | 29.01 | 26.69 | 27.00 | 27.73 | 27.89 | 30.14 | 30.25 | 30.77 | 30.89 |
| VII | 1&4 components | 2&3 components | 28.83 | 28.96 | 26.88 | 26.81 | 27.52 | 27.95 | 30.04 | 30.27 | 30.87 | 30.81 |
| VIII | 1st cultivar included | Not included | 29.08 | 28.73 | 26.89 | 26.78 | 28.06 | 27.52 | **30.56** | **29.77** | 30.80 | 30.86 |
| IX | 2nd cultivar included | Not included | 28.91 | 28.92 | 26.71 | 26.98 | 27.91 | 27.68 | 30.13 | 30.26 | 30.89 | 30.75 |
| X | 3rd cultivar included | Not included | 28.85 | 28.99 | 26.68 | 27.02 | 27.91 | 27.69 | **29.99** | **30.41** | 30.80 | 30.86 |
| XI | 4th cultivar included | Not included | 28.77 | 29.08 | 26.50 | 27.22 | 27.78 | 27.83 | 30.06 | 30.34 | 30.74 | 30.93 |
| XII | Continuous ripening | Discrete ripening | 28.85 | 29.04 | 26.64 | 27.23 | 27.74 | 27.94 | 30.10 | 30.36 | 30.91 | 30.65 |
| XIII | Normal duration (degree 1) | others | 28.84 | 28.94 | 27.08 | 26.75 | 27.37 | 27.97 | 30.08 | 30.23 | 30.86 | 30.81 |
| XIV | Extended duration (degree 2) | others | 28.93 | 28.87 | 26.88 | 26.68 | 27.83 | 27.72 | 30.21 | 30.12 | 30.79 | 30.97 |
| XV | Long duration (degree 3) | others | 29.10 | 28.85 | 26.97 | 26.79 | 28.24 | 27.65 | 30.30 | 30.15 | 30.88 | 30.81 |
| XVI | Very long duration (degree 4) | others | 28.83 | 28.94 | 26.58 | 26.93 | 27.88 | 27.78 | 30.25 | 30.17 | 30.63 | 30.90 |
|  | Group A | Group B | Group A | Group B | Group A | Group B | Group A | Group B | Group A | Group B | Group A | Group B |
| I | Monoculture | Mixture | 30.31 | 30.44 | 30.16 | 29.40 | 30.06 | 31.01 | 30.27 | 30.56 | 30.74 | 30.80 |
| II | 2-component | others | 30.56 | 30.30 | 29.66 | 29.56 | 31.10 | 30.53 | 30.62 | 30.39 | 30.86 | 30.73 |
| III | 3-component | others | 30.27 | 30.45 | 28.93 | 29.84 | 30.95 | 30.69 | 30.60 | 30.44 | 30.60 | 30.84 |
| IV | 4-component | others | 30.42 | 30.40 | 29.72 | 29.59 | 30.73 | 30.76 | 30.05 | 30.51 | 31.19 | 30.75 |
| V | 1&2 components | 3&4 components | 30.46 | 30.30 | 29.86 | 29.09 | 30.68 | 30.90 | 30.48 | 30.49 | 30.81 | 30.72 |
| VI | 1&3 components | 2&4 components | 30.29 | 30.54 | 29.54 | 29.67 | 30.50 | 31.05 | 30.43 | 30.54 | 30.67 | 30.90 |
| VII | 1&4 components | 2&3 components | 30.33 | 30.44 | 30.07 | 29.37 | 30.19 | 31.04 | 30.23 | 30.61 | 30.83 | 30.76 |
| VIII | 1st cultivar included | Not included | 30.52 | 30.27 | 29.61 | 29.59 | 30.95 | 30.54 | **30.86** | **30.05** | 30.67 | 30.91 |
| IX | 2nd cultivar included | Not included | 30.43 | 30.38 | 29.45 | 29.77 | 30.90 | 30.59 | 30.50 | 30.46 | 30.84 | 30.71 |
| X | 3rd cultivar included | Not included | 30.39 | 30.43 | 29.39 | 29.84 | 30.91 | 30.58 | 30.31 | 30.68 | 30.93 | 30.61 |
| XI | 4th cultivar included | Not included | 30.28 | 30.55 | 29.37 | 29.87 | 30.71 | 30.82 | 30.32 | 30.67 | 30.71 | 30.86 |
| XII | Continuous ripening | Discrete ripening | 30.39 | 30.44 | 29.45 | 29.90 | 30.77 | 30.72 | 30.41 | 30.62 | 30.91 | 30.52 |
| XIII | Normal duration (degree 1) | others | 30.31 | 30.44 | 30.16 | 29.40 | 30.06 | 31.01 | 30.27 | 30.56 | 30.74 | 30.80 |
| XIV | Extended duration (degree 2) | others | 30.39 | 30.46 | 29.70 | 29.20 | 30.65 | 31.19 | 30.48 | 30.48 | 30.73 | 30.98 |
| XV | Long duration (degree 3) | others | 30.59 | 30.34 | 29.42 | 29.66 | 31.36 | 30.54 | 30.70 | 30.40 | 30.88 | 30.75 |
| XVI | Very long duration (degree 4) | others | 30.28 | 30.45 | 29.52 | 29.63 | 30.53 | 30.84 | 30.48 | 30.48 | 30.57 | 30.86 |

| Contrast | Comparison |  | Overall |  | 1st year WI |  | 1st year DI |  | 2nd year WI |  | 2nd year DI |  |
| --- | --- | --- | --- | --- | --- | --- | --- | --- | --- | --- | --- | --- |
|  | Group A | Group B | Group A | Group B | Group A | Group B | Group A | Group B | Group A | Group B | Group A | Group B |
| I | Monoculture | Mixture | 29.71 | 29.97 | 28.71 | 28.92 | 29.28 | 30.22 | 29.88 | 29.90 | 30.98 | 30.83 |
| II | 2-component | others | 29.99 | 29.84 | 29.11 | 28.70 | 29.93 | 29.99 | 30.00 | 29.83 | 30.93 | 30.84 |
| III | 3-component | others | 30.02 | 29.86 | 28.92 | 28.84 | 30.61 | 29.74 | 29.79 | 29.94 | 30.75 | 30.92 |
| IV | 4-component | others | 29.63 | 29.92 | 27.82 | 28.94 | 30.39 | 29.94 | 29.75 | 29.91 | 30.58 | 30.89 |
| V | 1&2 components | 3&4 components | 29.88 | 29.94 | 28.95 | 28.70 | 29.67 | 30.57 | 29.95 | 29.78 | 30.95 | 30.72 |
| VI | 1&3 components | 2&4 components | 29.87 | 29.94 | 28.81 | 28.92 | 29.95 | 30.00 | 29.84 | 29.97 | 30.87 | 30.88 |
| VII | 1&4 components | 2&3 components | 29.70 | 30.00 | 28.53 | 29.03 | 29.50 | 30.21 | 29.85 | 29.92 | 30.90 | 30.86 |
| VIII | 1st cultivar included | Not included | **30.23** | **29.53** | **29.47** | **28.18** | 30.26 | 29.64 | **30.26** | **29.49** | 30.93 | 30.80 |
| IX | 2nd cultivar included | Not included | 30.04 | 29.74 | 29.03 | 28.68 | 30.43 | 29.44 | 29.76 | 30.05 | 30.94 | 30.79 |
| X | 3rd cultivar included | Not included | 29.79 | 30.03 | 28.73 | 29.01 | 30.09 | 29.84 | **29.68** | **30.14** | 30.66 | 31.11 |
| XI | 4th cultivar included | Not included | **29.63** | **30.21** | **28.08** | **29.76** | 29.88 | 30.07 | 29.80 | 30.00 | 30.76 | 31.00 |
| XII | Continuous ripening | Discrete ripening | 29.79 | 30.11 | 28.66 | 29.27 | 29.80 | 30.31 | 29.79 | 30.10 | 30.92 | 30.78 |
| XIII | Normal duration (degree 1) | others | 29.71 | 29.97 | 28.71 | 28.92 | 29.28 | 30.22 | 29.88 | 29.90 | 30.98 | 30.83 |
| XIV | Extended duration (degree 2) | others | 29.94 | 29.73 | 28.93 | 28.62 | 30.06 | 29.60 | 29.93 | 29.76 | 30.85 | 30.95 |
| XV | Long duration (degree 3) | others | 30.23 | 29.78 | 29.42 | 28.66 | 30.71 | 29.70 | 29.89 | 29.90 | 30.89 | 30.87 |
| XVI | Very long duration (degree 4) | others | 29.89 | 29.91 | 28.65 | 28.94 | 30.20 | 29.89 | 30.02 | 29.85 | 30.68 | 30.94 |

**Table S2.**
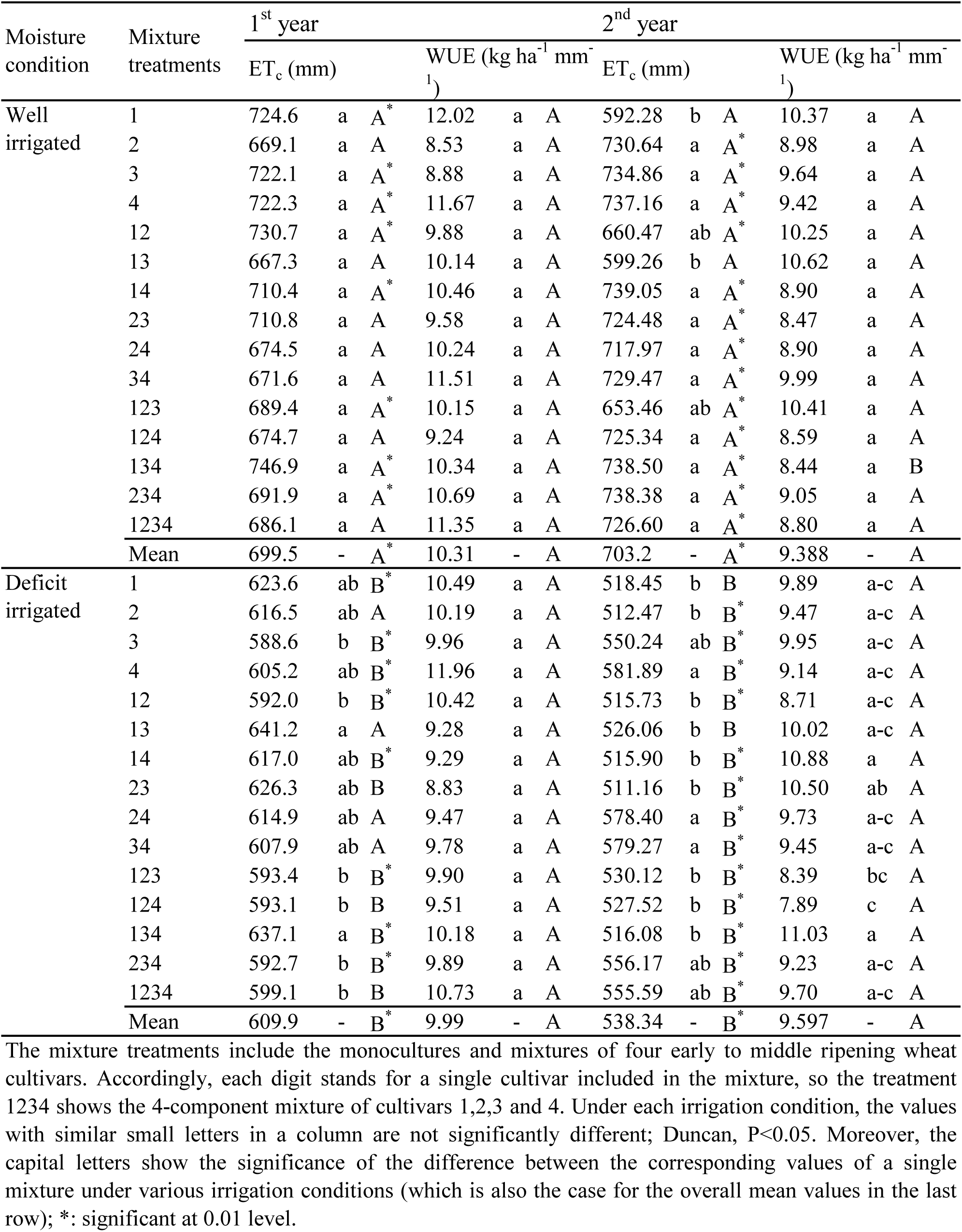
Evapotranspiration (ET_c_′) and water use efficiency (WUE) of mixture treatments under different irrigation conditions and years.

**Table S3.** Orthogonal analyses of ET_c_’ (mm).

| Contrast | Comparison |  | Overall |  | 1<sup>st</sup> year WI |  | 1<sup>st</sup> year DI |  | 2<sup>nd</sup> year WI |  | 2<sup>nd</sup> year DI |  |
| --- | --- | --- | --- | --- | --- | --- | --- | --- | --- | --- | --- | --- |
|  | Group A | Group B | Group A | Group B | Group A | Group B | Group A | Group B | Group A | Group B | Group A | Group B |
| I | Monoculture | Mixture | 639.4 | 637.1 | 709.5 | 695.8 | 608.4 | 610.4 | 698.7 | 704.8 | 540.8 | 537.5 |
| II | 2-component | others | 635.9 | 638.9 | 694.2 | 703.0 | 616.6 | 605.5 | 695.1 | 708.6 | 537.8 | 538.7 |
| III | 3-component | others | 637.8 | 637.7 | 700.7 | 699.0 | 604.1 | 612.0 | 713.9 | 699.3 | 532.5 | 540.5 |
| IV | 4-component | others | 641.8 | 637.4 | 686.1 | 700.5 | 599.1 | 610.7 | 726.6 | 701.5 | 555.6 | 537.1 |
| V | 1&2 components | 3&4 components | 637.3 | 638.6 | 700.3 | 697.8 | 613.3 | 603.1 | 696.6 | 716.5 | 539.0 | 537.1 |
| VI | 1&3 components | 2&4 components | 638.6 | 636.8 | 705.1 | 693.1 | 606.3 | 614.1 | 706.3 | 699.6 | 536.6 | 540.3 |
| VII | 1&4 components | 2&3 components | 639.9 | 636.7 | 704.8 | 696.8 | 606.6 | 611.6 | 704.3 | 702.6 | 543.7 | 535.6 |
| VIII | 1st cultivar included | Not included | 630.2 | 646.3 | 703.8 | 694.6 | 612.1 | 607.4 | 679.4 | 730.4 | **525.7** | **552.8** |
| IX | 2nd cultivar included | Not included | 635.0 | 640.9 | 690.9 | 709.3 | 603.5 | 617.2 | 709.7 | 695.8 | 535.9 | 541.1 |
| X | 3rd cultivar included | Not included | 638.8 | 636.5 | 698.3 | 700.9 | 610.8 | 608.9 | 705.6 | 700.4 | 540.6 | 535.8 |
| XI | 4th cultivar included | Not included | 647.1 | 627.0 | 697.3 | 702.0 | 608.4 | 611.7 | 731.6 | 670.8 | **551.4** | **523.5** |
| XII | Continuous ripening | Discrete ripening | 637.6 | 638.1 | 701.9 | 694.8 | **604.5** | **620.7** | 702.8 | 704.0 | 541.1 | 532.8 |
| XIII | Normal duration (degree 1) | others | 639.4 | 637.1 | 709.5 | 695.8 | 608.4 | 610.4 | 698.7 | 704.8 | 540.8 | 537.5 |
| XIV | Extended duration (degree 2) | others | 637.6 | 638.3 | 698.3 | 704.4 | 610.2 | 608.8 | 702.8 | 704.8 | 539.1 | 535.4 |
| XV | Long duration (degree 3) | others | 629.1 | 640.9 | 680.8 | 706.3 | 610.6 | 609.7 | 677.3 | 712.6 | 547.7 | 534.9 |
| XVI | Very long duration (degree 4) | others | 644.3 | 635.3 | 704.5 | 697.7 | 611.6 | 609.3 | **732.4** | **692.6** | 528.8 | 541.8 |

WI and DI: well- and deficit-irrigated, respectively. Overall: analyzed using the data of all conditions i.e. under WI and DI of the two years.

**Table S4.** . Orthogonal analyses of WUE (kg ha^−1^ mm^−1^).

| Contrast | Comparison |  | Overall |  | 1<sup>st</sup> year WI |  | 1<sup>st</sup> year DI |  | 2<sup>nd</sup> year WI |  | 2<sup>nd</sup> year DI |  |
| --- | --- | --- | --- | --- | --- | --- | --- | --- | --- | --- | --- | --- |
|  | Group A | Group B | Group A | Group B | Group A | Group B | Group A | Group B | Group A | Group B | Group A | Group B |
| I | Monoculture | Mixture | 10.03 | 9.74 | 10.27 | 10.32 | 10.65 | 9.75 | 9.60 | 9.31 | 9.61 | 9.59 |
| II | 2-component | others | 9.80 | 9.83 | 10.30 | 10.32 | 9.51 | 10.31 | 9.52 | 9.30 | 9.88 | 9.41 |
| III | 3-component | others | 9.56 | 9.92 | 10.10 | 10.39 | 9.87 | 10.04 | 9.12 | 9.49 | 9.13 | 9.77 |
| IV | 4-component | others | 10.14 | 9.80 | 11.35 | 10.24 | 10.73 | 9.94 | 8.80 | 9.43 | 9.70 | 9.59 |
| V | 1&2 components | 3&4 components | 9.90 | 9.67 | 10.29 | 10.35 | 9.97 | 10.04 | 9.55 | 9.06 | 9.77 | 9.25 |
| VI | 1&3 components | 2&4 components | 9.80 | 9.85 | 10.19 | 10.45 | 10.26 | 9.68 | 9.36 | 9.42 | 9.37 | 9.85 |
| VII | 1&4 components | 2&3 components | 10.06 | 9.70 | 10.49 | 10.22 | 10.67 | 9.65 | 9.44 | 9.36 | 9.63 | 9.58 |
| VIII | 1st cultivar included | Not included | 9.88 | 9.75 | 10.45 | 10.16 | 9.97 | 10.01 | 9.55 | 9.21 | 9.56 | 9.64 |
| IX | 2nd cultivar included | Not included | **9.55** | **10.13** | 9.96 | 10.72 | 9.87 | 10.13 | 9.18 | 9.63 | 9.20 | 10.05 |
| X | 3rd cultivar included | Not included | 9.84 | 9.80 | 10.33 | 10.29 | 9.82 | 10.19 | 9.43 | 9.35 | 9.78 | 9.38 |
| XI | 4th cultivar included | Not included | 9.86 | 9.78 | 10.69 | 9.88 | 10.10 | 9.87 | 9.01 | 9.82 | 9.63 | 9.56 |
| XII | Continuous ripening | Discrete ripening | 9.90 | 9.66 | 10.43 | 10.08 | 10.21 | 9.54 | 9.54 | 9.09 | 9.44 | 9.91 |
| XIII | Normal duration (degree 1) | others | 10.03 | 9.74 | 10.27 | 10.32 | 10.65 | 9.75 | 9.60 | 9.31 | 9.61 | 9.59 |
| XIV | Extended duration (degree 2) | others | 9.83 | 9.78 | 10.31 | 10.32 | 10.07 | 9.68 | 9.34 | 9.57 | 9.61 | 9.55 |
| XV | Long duration (degree 3) | others | 9.76 | 9.85 | 10.31 | 10.31 | 9.63 | 10.12 | 9.74 | 9.26 | 9.34 | 9.69 |
| XVI | Very long duration (degree 4) | others | 9.71 | 9.86 | 10.35 | 10.30 | 9.92 | 10.02 | **8.68** | **9.64** | 9.87 | 9.50 |

WI and DI: well- and deficit-irrigated, respectively. Overall: analyzed using the data of all conditions i.e. under WI and DI of the two years.

